# Proposal of *Patescibacterium danicum* gen. nov., sp. nov. in the ubiquitous ultrasmall bacterial phylum *Patescibacteriota* phyl. nov.

**DOI:** 10.1101/2024.11.01.618662

**Authors:** Zuzanna Dutkiewicz, Caitlin Margaret Singleton, Mantas Sereika, Juan C. Villada, Aaron J. Mussig, Maria Chuvochina, Mads Albertsen, Frederik Schulz, Tanja Woyke, Per Halkjær Nielsen, Philip Hugenholtz, Christian Rinke

## Abstract

*Candidatus* Patescibacteria is a diverse bacterial phylum that is notable for members with ultrasmall cell size, reduced genomes, limited metabolic capabilities and dependence on other prokaryotic hosts. Despite the prevalence of the name *Ca.* Patescibacteria in the scientific literature, it is not officially recognized under the International Code of Nomenclature of Prokaryotes (ICNP) and lacks a nomenclatural type. Here, we rectify this situation by describing two closely related circular metagenome-assembled genomes (MAGs) and by proposing one of them (ABY1^TS^) to serve as the nomenclatural type for the species *Patescibacterium danicum*^TS^ gen. nov., sp. nov. according to the rules of the SeqCode. Rank-normalized phylogenomic inference confirmed the stable placement of *P. danicum*^TS^ in the class ABY1 within *Ca.* Patescibacteria. Based on these results, we propose *Patescibacterium* gen. nov. to serve as the type genus for associated higher taxa, including the phylum *Patescibacteriota* phyl. nov. We complement our proposal with a genomic characterization, metabolic reconstruction, and biogeographical analysis of *Patescibacterium*. Our results confirm small genome sizes (< 1Mbp), low GC content (>36%), and the occurrence of long gene coding insertions in the 23S rRNA sequences, along with reduced metabolic potential, inferred symbiotic lifestyle, and a global distribution. In summary, this effort, focused on the fourth-largest phylum in the bacterial domain, aims to provide nomenclatural stability in the scientific literature.

## Introduction

The name *Candidatus* Patescibacteria was originally proposed for a bacterial superphylum comprising the lineages *Ca.* Microgenomates (formerly OP11) (Hugenholtz et al. 1998), *Ca.* Parcubacteria (formerly OD1) (Harris, Kelley, and Pace 2004), and *Ca.* Gracilibacteria (formerly GN02 or BD1-5) (L. Li, Kato, and Horikoshi 1999; Ley et al. 2006) based on phylogenomic inferences of single amplified and metagenome-assembled genomes (Rinke et al. 2013). Derived from *patesco* (Latin), meaning bare, the name *Ca.* Patescibacteria was chosen to reflect the limited metabolic capacities, that are characteristic for this group (Rinke et al. 2013). The discovery of additional major lineages within *Ca.* Patescibacteria led to the proposal of the Candidate Phyla Radiation (CPR), described as a group of bacteria with consistently small genomes and cell sizes that comprises over 15% of the bacterial domain (Castelle and Banfield 2018; Brown et al. 2015). The names *Ca.* Patescibacteria and CPR are currently used as synonyms, noting that neither are validly published under the International Code of Nomenclature of Prokaryotes (ICNP; see below), because the former has *Candidatus* status and the latter is a vernacular name (LPSN 2024). More recently, *Ca.* Patescibacteria/CPR were classified as a single bacterial phylum in the Genome Taxonomy Database (GTDB) based on rank normalization using relative evolutionary divergence (Avise and Johns 1999; Parks et al. 2018). The name *Ca.* Patescibacteria currently serves as a provisional name for this phylum in GTDB (Parks et al. 2018). However, this name cannot be validated without a nomenclatural type, a problem we address in the present study by proposing a type genus under the SeqCode based on a high-quality circular genome sequence obtained from a metagenomic survey of Danish wastewater treatment plants.

The International Committee on Systematics of Prokaryotes (ICSP) voted to include the rank of phylum in the ICNP (Parker, Tindall, and Garrity 2019) in 2021, based on proposals by Oren et al. (Oren et al. 2015) and Whitman et al. (Whitman et al. 2018). The ICSP further decided that the nomenclatural type of a phylum is one of the contained genera. This ruling means that all phylum names must be based on the name of a genus as its nomenclatural type, and names of phyla are formed by the addition of the suffix -*ota* to the stem of the name of the type genus (Oren and Garrity 2021). The subsequent proposal to accept DNA sequences as nomenclatural types, which would have allowed the inclusion of uncultured prokaryotes in the ICNP, was rejected by the ICSP and triggered the development of the SeqCode (Hedlund et al. 2022). The SeqCode accepts genome sequences as type material for uncultivated and cultivated prokaryotes and follows naming rules for higher taxa similar to those of the ICNP (Whitman et al. 2024). Here, we follow the SeqCode and describe the genome serving as the nomenclatural type for the novel species *Patescibacterium danicum* gen. nov. sp. nov., and propose the genus *Patescibacterium* gen. nov. as the nomenclatural type of associated higher taxa, including the phylum *Patescibacteriota* phyl. nov. All proposed names have been submitted for validation under the SeqCode to provide a genus, family, order, class, and phylum name with standing in nomenclature, a measure that supports taxonomic congruence in scientific literature.

In addition, we inferred lifestyle, biogeography and metabolic capabilities of the proposed type genus *Patescibacterium*. In general, a symbiotic or parasitic lifestyle has been assumed for many *Patescibacteriota*. Reduced genomes and the associated small cell sizes, e.g., cell diameters of less than 200nm (Luef et al. 2015; He et al. 2015), in combination with the lack of essential biosynthetic and metabolic pathways (Rinke et al. 2013; He et al. 2015; Brown et al. 2015) support this assumption. A limited number of cultures has since confirmed host dependencies for members of this phylum, e.g. through co-cultures of Saccharimonadia (TM7) (Ibrahim et al. 2021; He et al. 2015) or Absconditabacterales (Yakimov et al. 2022) and their respective hosts. Probable free-living members have also been reported, e.g., a study visualizing fresh water *Patescibacteriota* found many cells not to be attached to other organisms or to be associated with organic particles, suggesting a host independent lifestyle (Chiriac et al. 2022). Here, we applied a machine learning based classifier to explore host dependencies of the type genus *Patescibacterium*, resulting in a predicted symbiotic lifestyle. We found a global distribution of this genus and a preference for aquatic habitats, in accordance with previous reports for members of this phylum (Tian et al. 2020; Haro-Moreno et al. 2023). Our metabolic reconstruction suggests a lack of essential biosynthesis pathways and characterizes the genus *Patescibacterium* as a fermentative organotroph.

In summary, our proposal of *Patescibacterium danicum* and the associated higher ranks, up to phylum level, should contribute to taxonomic stability of *Patescibacteriota*, one of the most diverse phyla in the bacterial domain.

## Material and methods

### Genome recovery

The closed metagenome assembled genome (MAG) ‘ABY1’ was obtained via differential coverage binning of Oxford Nanopore Technology (ONT) long read assemblies polished by Illumina short read data from Mariagerfjord wastewater treatment plant (WWTP) (SAMN14825711) and published previously (Singleton et al. 2021). The closed genome ‘Fred.cMAG.1’ was recovered via re-assembly by first performing taxonomic classification of the contigs from the Fredericia WWTP assembly (SAMN14825693) using mmseqs2 v.14.7e284 (Steinegger and Söding 2017) to the NCBI nr database (release 2023-09-11). Next, ONT (SRR11673978) and Illumina (SRR11674035) reads of the same sample were mapped to the contigs with Minimap2 v22.26 (H. Li 2018) and Samtools v1.18 (Danecek et al. 2021) using the settings “-ax map-ont” for ONT and “-ax sr” for short reads. The mapped reads were then subsetted to select reads mapping to contigs taxonomically classified, using NCBI taxonomy, as "Candidatus *Falkowbacteria*" by mmseqs2. The subsetted ONT reads were assembled using Flye v2.9.3 (Kolmogorov et al. 2020) with the following settings: “--nano-raw”, “--meta”, “--extra-params min_read_cov_cutoff=20”. The assembled circular contig was then polished with the subsetted Illumina reads using Racon v1.5.0 (Vaser et al. 2017) resulting in the ‘Fred.cMAG.1’ genome.

Four genomes closely related to “ABY1’ and ‘Fred.cMAG.1’ were derived from assembled contigs obtained from the IMG/M data management and analysis system (Chen et al. 2023), that were labeled as metagenomes (IMGM prefix in MAGs) projects. Only contigs with a length of equal or greater than 5 kbp were considered. Binning was performed on each sample with MetaBAT2 (parameters: --minContig 5000 --minClsSize 20000 --cvExt). Initial gene calling was performed on each bin with Prodigal (v2.6.3), to create FAA input files for taxonomic assignments by GTDB-Tk v2.3.2 (Chaumeil et al. 2022) and for symbiont classification by symcla (see “*Lifestyle predictions*” below).

An additional four public genomes (GCA_002344425.1, GCA_002433955.1, GCA_002293885.1, GCA_002343995.1) were selected based on their taxonomic assignment to genus UBA919 in the Genome Taxonomy Database (GTDB) (Chuvochina et al. 2023) and were subsequently obtained from NCBI RefSeq (Haft et al. 2018).

Completeness and contamination estimates of all MAGs were calculated with CheckM2 (v1.0.1) (Chklovski et al. 2022). FastANI (Jain et al. 2018) was used to calculate the Average Nucleotide Identity (ANI) between genomes, using default settings, e.g. --fragLen 3,000 and –minFraction 0.2. Estimated sizes of complete genomes were calculated (=genome size/estimated completeness by CheckM2 (v1.0.1) * 100) based on all ‘Patescibacteria’ in GTDB Release 09-RS220 (24th April 2024).

Prediction of the secondary structures of ribosomal 5S and 23S RNA genes were made with the webserver StructRNAfinder (Arias-Carrasco et al. 2018).

### Phylogeny

Phylogenomic inferences were calculated from the bacterial GTDB alignment comprising 120 marker proteins, whereby the alignment was created with GTDB-Tk (v2.3.2; GTDB-Tk reference data version r214) (Chaumeil et al. 2022). Initial trees were calculated with FastTree2 (Price, Dehal, and Arkin 2010) and decorated with taxonomic information using PhyloRank (v0.1.12; https://github.com/donovan-h-parks/PhyloRank). Next, phylogenomic trees were inferred with IQ-Tree (v2.3.6) (Nguyen et al. 2015) using a FastTree2 with 3406 taxa as starting tree and the command “iqtree -s <input alignment> -nt 12 -m LG+C10+F+G -ft <starting tree> -pre <prefix of output files>”. Next, bootstrap trees were inferred running the same commands as above, but by adding the flag --bb 1000 for 1000 bootstrap support values via the ultrafast bootstrap approximation (UFBoot) (Hoang et al. 2018). Resulting trees were decorated with PhyloRank (v0.1.12), and visualized with ARB (Ludwig et al. 2004) and iTOL (Letunic and Bork 2021).

### Lifestyle predictions

Symbiotic lifestyle predictions were generated with the machine learning-based symbiont classifier *symcla* (https://github.com/NeLLi-team/symcla). Analyzing the 10 genomes characterized in this study along with GTDB R214 reference genomes, we followed the recommended interpretation to predict lifestyles based on the symcla scores as follows: symcla_score <= 0.42: free-living; 0.42 < symcla_score < 1.21: symbiont/host-associated; symcla_score >= 1.21: symbiont/intracellular. It should be noted that symcla was designed to minimize false positive hits for symbionts, at the expense of higher probabilities for false negative hits, i.e. symbiont/host-associated genomes might get assigned a symcla score below 0.42, and symbiont/intracellular genomes might get a symcla score lower than 1.21.

### Metabolic reconstruction

Gene calling of all MAGs was carried out with Bakta (Schwengers et al. 2021) and the resulting amino acid sequences (*.faa files) were annotated with KEGG BlastKEGG Orthology Ank Links Annotation (BlastKOALA) (https://www.kegg.jp/blastkoala/) (Kanehisa, Sato, and Morishima 2016) and gapseq (Zimmermann, Kaleta, and Waschina 2021). The resulting annotations were visualized using KEGG Mapper Reconstruction Result (Kanehisa, Sato, and Kawashima 2022). The presence of the identified genes across all MAGs was visualized using the ggplot2 library in R (Team 2014) to generate a heatmap. The sketch providing an overview of the metabolic reconstruction of the type genus was created with the open-source LibreOffice Draw (https://www.libreoffice.org). The unusually long proteins were annotated with a Quick BLASTP (Altschul et al. 2005) alignment, and by searching protein homology by domain architecture. For the latter, the Conserved Domain Architecture Retrieval Tool (CDART) (Geer et al. 2002) was used to find protein similarities across significant evolutionary distances. CDART uses sensitive domain profiles instead of direct sequence similarity, whereby a domain architecture was defined as the sequential order of conserved domains, i.e. functional units, in a protein sequence. The NCBI wide homology search of the long proteins involved obtaining a list of all 2,111,842 protein sequences assigned to the “Patescibacteria group” under the NCBI Taxonomy ID “1783273”, using the search term “txid1783273[organism:exp]”. The list was downloaded as a “Summary file” and the entries were sorted by the number of amino acids, and a threshold of 4,000 amino acids was established to retain only the longest sequences, resulting in 149 proteins. All 149 protein sequences were downloaded to create a database that was subsequently used for a HMM search. HMM profiles were created by “phmmer --domtblout output1 -A output2 --qformat fasta query_file database_file”. The analysis revealed only two hits for the same protein (QQS60272.1 MAG: fibronectin type III domain-containing protein and QQG53011.1 MAG: fibronectin type III domain-containing protein) which turned out to be self-hits to ABY1 (GCA_016699775.1) and Fred.cMAG.1 (GCA_964214775.1). However, we did find that many of these large proteins contained the conserved domains FhaB and MJ1470/ DUF2341, which were counted and visualized in a manually created gene neighborhood figure.

### Biogeography

All 16S rRNA gene profiles resulted from samples obtained from activated sludge WWTPs from the MiDAS global survey and were sequenced using Illumina HiSeq 2500 as previously described (Dueholm et al. 2022). Abundance profiles of the genomes were based on a set of 35 bacterial and 37 archaeal (in total 59) single copy marker genes, extracted with the tool singleM (Woodcroft et al. 2024) from 248,559 publicly available metagenomes at the NCBI Sequence Read Archive (SRA). An evaluation of the number of singleM markers in *Patescibacteriota* class representative genomes revealed that they contain on average 92.1% (32.2 ± 3.01) of the singleM marker genes, well above the 10% threshold for confident assignments (**Suppl. Text**). Samples from close geographic proximity were aggregated in clusters with DBSCAN (https://scikit-learn.org/stable/modules/generated/dbscan-function.html) from the sklearn python package and plotted on a world map projection using the cartopy python package (https://scitools.org.uk/cartopy).

### Phylogenetic diversity

The Phylogenetic diversity (PD) was calculated with the in-house pipeline *gtdb-phylogenetic-diversity* (https://github.com/aaronmussig/gtdb-phylogenetic-diversity). The pipeline calculates the sum of all descendant branch lengths starting from the internal node that is decorated with the phylum label. The stem of the named node (the phylum node), is included in this sum. For singletons (*i.e.* taxa that have no named node, as they are the sole representative of a phylum), the stem length from the named leaf node is used. Next, the values are normalized by the total sum of all calculated per-phylum PD scores (i.e. both named, and singletons). To calculate the standard PD values for this study, the pipeline was run on the bacterial GTDB R220 tree “bac120.tree” (https://gtdb.ecogenomic.org). To obtain the adjusted PD value, the script was run on the rank normalized version of the bac120.tree, created with the tool PhyloRank (v0.1.12; https://github.com/donovan-h-parks/PhyloRank), that calculates the relative evolutionary divergence (RED) of all taxa in a tree.

## Results

### Genome recovery and phylogeny

Binning of hybrid assemblies, *i.e.* long-read assemblies followed by short-read polishing, yielded two complete metagenome assembled genomes (MAGs) from Danish wastewater treatment plants (WWTPs). MAG ‘ABY1’ was recovered from Mariagerfjord WWTP (Singleton et al. 2021), and the second MAG ‘Fred.cMAG.1’ was obtained from the Fredericia WWTP (see Methods). Both WWTPs treat municipal wastewater using biological nutrient removal and the activated sludge process, in addition the Fredericia WWTP also treats industrial wastewater (Nierychlo et al. 2020). ‘ABY1’ and ‘Fred.cMAG.1’ represent complete, single contig genomes with sizes of 939 and 917 kbp, and coding densities of 88.2% and 85.0%, respectively (**Table 1**). Both genomes are characterized by a GC content of 35.4%, and a full set of ribosomal RNA genes (**Table 1**). The 23S rRNA genes of these complete MAGs were unusually long (4798 bp each) and characterized by two insertion sequences (**Fig. S1**) that contained several open reading frames, the longest of them was identified as coding for homing endonuclease, specifically as DNA cleaving enzymes of the LAGLIDADG family (**Fig. S2**).

**Table 1.** Genome characteristics of the metagenome-assembled genomes (MAGs) included in this study. Completeness and contamination estimates and coding density have been calculated with CheckM2. Note that the completeness of the complete circular genomes ‘ABY1’ and ‘Fred.cMAG.1’ was slightly underestimated by CheckM2. Genomes that qualify as high-quality draft MAGs according to MIMAG standards are highlighted in bold (see *Suppl. Text*). Abbreviations: Data source = including National Center for Biotechnology Information (NCBI), Singleton et al. 2021 (see Methods), Integrated Microbial Genomes & Microbiomes (IMG/M); MAG/ acc. no. = MAG name / NCBI or IMG/M accession number; Genome size (bp) = MAG size in base pairs, Con (#) = number of contigs; Comp (%) = estimated completeness in %; Con (%) = estimated contamination in %; GC (%) = % GC content; CD (%) = % coding density; Prot (#) = number of encoded proteins; 5-16-23 (#) = number of 5S, 16S, and 23S rRNA genes. Numbers of 5S and 23S genes were manually curated following Bakta annotations.

| Data source | MAG / acc. no. | Genome (bp) | Con (#) | Comp (%) | Con (%) | GC (%) | CD (%) | Prot (#) | 5-16-23 (#) |
| --- | --- | --- | --- | --- | --- | --- | --- | --- | --- |
| Singleton et al. 2021 | **ABY1<sup>TS</sup>**<br>**(GCA_016699775.1)** | 938,851 | 1 | 96.03* | 0.69 | 35.4 | 88.2 | 890 | 1-1-1 |
| This study | Fred.cMAG.1<br>(GCA_964214775.1) | 917,454 | 1 | 95.89* | 0.9 | 35.4 | 85 | 978 | 1-1-1 |
| NCBI | **GCA_002344425.1** | 752,947 | 38 | 95.69 | 0.53 | 43.9 | 92.3 | 792 | 1-1-1 |
| NCBI | GCA_002433955.1** | 748,244 | 38 | 91.4 | 0.93 | 43.9 | 91.2 | 777 | 1-2-1 |
| NCBI | GCA_002293885.1 | 690,185 | 59 | 89.88 | 0.71 | 44.2 | 90.4 | 728 | 1-0-1 |
| NCBI | GCA_002343995.1 | 746,423 | 24 | 95.51 | 1.79 | 43.9 | 93.1 | 774 | 2-1-1 |
| IMG/M | IMGM3300014059_BIN201 | 912,772 | 53 | 90.14 | 0.32 | 36.1 | 89.5 | 959 | 0-1-1 |
| IMG/M | **IMGM3300014204_BIN854** | 962,299 | 12 | 96.48 | 0.53 | 35.2 | 90.8 | 973 | 1-1-1 |
| IMG/M | **IMGM3300029288_BIN286** | 915,973 | 10 | 96.74 | 0.85 | 35.2 | 88.7 | 916 | 1-1-1 |
| IMG/M | IMGM3300030493_BIN257 | 530,683 | 60 | 69.2 | 0.23 | 34.9 | 92.2 | 583 | 0-1-0 |

<sup>TS</sup> = designated type genome of the species. \*This is a complete, single contig genome, however, CheckM2 is underestimating genome completeness. \*\* Genome GCA\_002433955.1 contains two partial 16S rRNA genes on two contigs.

We included eight additional MAGs in our analysis, four binned from public IMG/M metagenomes and four obtained from NCBI RefSeq (**Table S1**). Genome quality assessments confirmed that eight of the ten MAGs included in this study fulfilled the MIMAG standards (Bowers et al. 2017) for high-quality genomes, in terms of estimated completeness (>90%) and contamination (<5%) (**Table 1**). Taxonomic assignments by GTDB-Tk (**Fig S3; Table S2**) and a subsequent phylogenomic inference (**Fig. 1a-c)** confirmed that all ten MAGs belong to the genus UBA919, family UBA917, order BM507, and class ABY1 in the phylum *Ca.* Patescibacteria. Comparison of average nucleotide identity (ANI) values (**Table S3**) and applying a threshold of >95% for the rank of species, supported phylogenomic clusters within genus UBA919 **(Fig. 1c)** and revealed several species level lineages. ‘ABY1’ and ‘Fred.cMAG.1’ share 97.7% ANI and belong to GTDB species UBA919 sp016699775 (Parks et al. 2022). All four MAGs obtained from NCBI formed a second species (>99% ANI), whereas each of the four IMG-derived MAGs represents a separate species within the genus UBA919 **(Fig. 1c; Table S3)**.

**Fig. 1.**
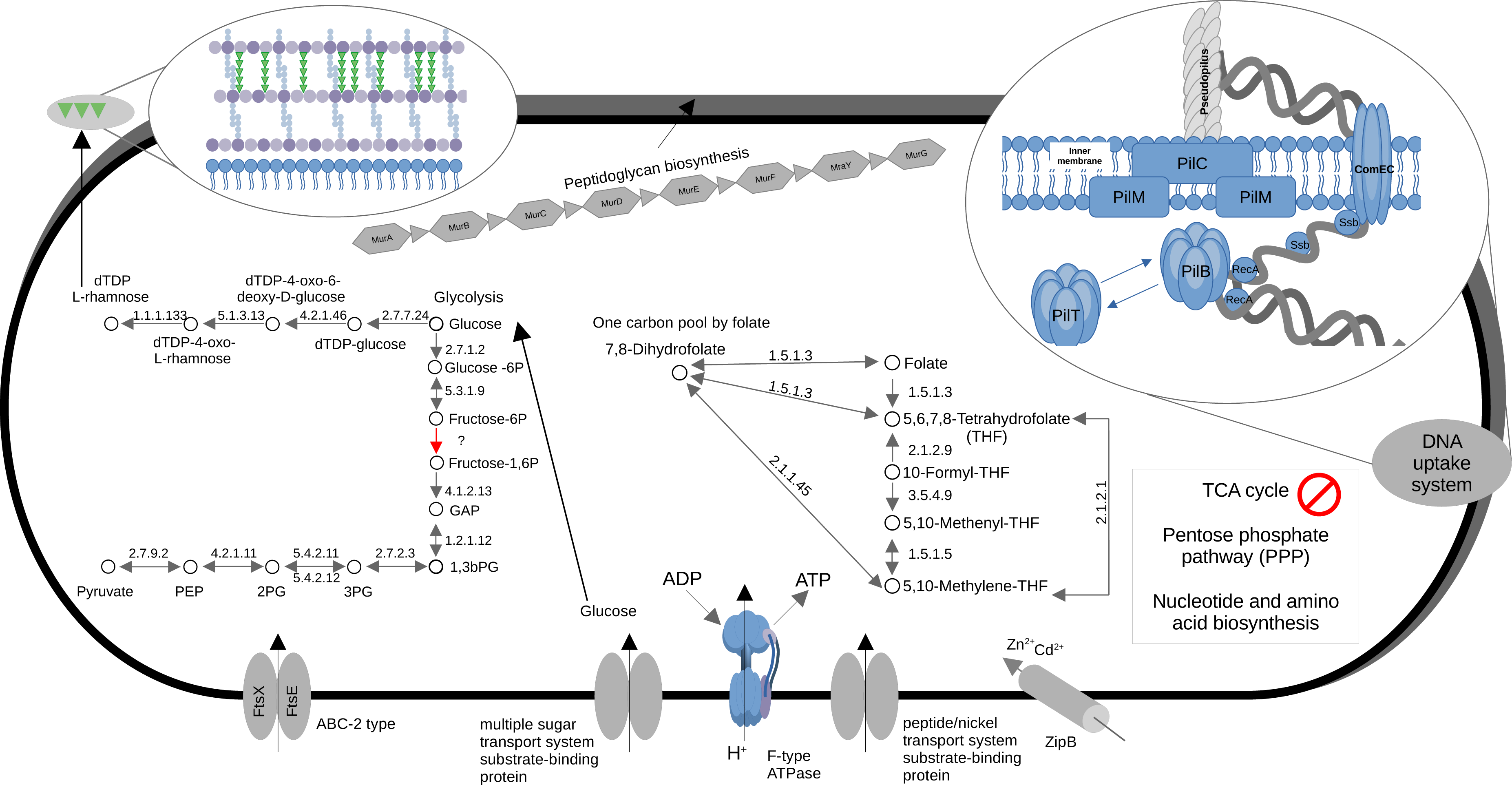
| Phylogenomic placement of the genome ABY1^TS^ within *Patescibacteriota* phyl. nov. Phylogeny was inferred with IQ-Tree from a 120 marker protein alignment of 3406 genomes. **a)** Phylogenomic tree, clustered on the phylum level, highlighting the monophyletic lineage *Patescibacteriota* phyl. nov. (gray triangle). **b)** Class level subsection of the tree, focused on *Patescibacteria* class nov. (GTDB r214 name ABY1) and its closest class level sister lineages. **c)** Order level subsection depicting *Patescibacteriales* ord. nov. (BM507) (outer gray box) containing the family *Patescibacteriaceae* fam. nov. (UBA917) (inner gray box), and the genus *Patescibacterium* gen. nov. (UBA919) (filled light gray box). The MAG ABY1^TS^ was placed within the species *Patescibacterium danicum* sp. nov. (UBA919 sp016699775) (upper dark gray box) in the genus *Patescibacterium*. For visualization purposes, the tree was rooted between the *Patescibacteriota* and all other phyla. The full tree is provided in the Supplementary Data. Ultra Fast bootstrap support values are shown below the nodes. Taxonomic assignments are based on GTDB r214 (https://gtdb.ecogenomic.org/). The MAG ABY1**^TS^** is marked with ^TS^ to indicate the status of the type species.

### Proposal of *Patescibacterium danicum* sp. nov. and associated higher taxa

Based on the recovery of two closed genomes (ABY1^TS^ and Fred.cMAG.1), the robust genome phylogeny, and ANI comparisons, we propose the species name *Patescibacterium danicum* sp. nov. This name was selected to honor the original description of the lineage ‘Patescibacteria’ based on patesco (Lat.), meaning bare, and to highlight the discovery of this lineage in samples from WWTPs in Denmark. We propose this species name under the SeqCode, based on the complete genome sequence of MAG ABY1^TS^ (GCA_016699775.1), an uncultured representative, to serve as the type genome of this species. We subsequently propose the genus *Patescibacterium* gen. nov., to serve as the nomenclatural type of the proposed family *Patescibacteriaceae* fam. nov., the order *Patescibacteriales* ord. nov., the class *Patescibacteriia* class nov. and the phylum *Patescibacteriota* phyl. nov. Detailed proposals are provided in the section “Description of new taxa according to the SeqCode” of this manuscript.

### Core metabolism of *Patescibacterium* gen. nov.

Metabolic reconstruction of all ten genomes assigned to the genus *Patescibacterium* revealed limited metabolic potential, consistent with other members of the phylum *Patescibacteriota*. *Patescibacterium* MAGs do not encode essential pathways for central carbon metabolism, including the pentose phosphate pathway (PPP) and tricarboxylic acid (TCA) cycle, and lack genes for nucleotide and amino acid biosynthesis, and for the degradation of fatty acids (**Fig. 2**). Inferred *de novo* nucleotide synthesis pathways were incomplete, with only BIN854 and BIN257 encoding the enzyme purine-nucleoside phosphorylase (punA; K03783), which is involved in adenine and guanine synthesis (**Table S4**). To potentially compensate for this absence of nucleotide synthesis, *Patescibacterium* possesses a range of encoded proteins required for the uptake of extracellular DNA. The inferred DNA uptake machinery includes a competence pseudopilus, a DNA membrane translocation compound, and an enzyme (RecA) that can catalyze the recombination of imported DNA with the bacterial chromosome (Ryan, Damke, and Shaffer 2023) (**Fig. 2, Table S5**). The pseudopilus gene neighborhood did not include genes for structures in the outer membrane, such as the major pilus subunit PilA. This finding, along with the detection of a complete cell wall synthesis operon (**Fig. 2; Table S6**), suggests that *Patescibacterium* has a cell envelope consisting of a single membrane and a murein (peptidoglycan) layer. Genes for the conversion of D-Glucose 1-phosphate to dTDP-L-rhamnose were also prevalent (**Table S6**). Rhamnose might be used instead of teichoic acids to produce rhamnose cell wall polysaccharides (**Fig. 2)**, as reported for lactic acid bacteria, such as streptococcal species (van der Beek et al. 2019). The inferred morphology is that of a rod shaped bacterium, based on the presence of genes for the rod shape determining proteins RodA, MreB and MreC, in all MAGs except for the least complete BIN257 that is missing RodA (**Table S7**). These inferred rod shape indicating proteins were also detected in the majority of genome representatives for the 24 *Patescibacteriota* classes in GTDB r214 (**Fig. S4**), indicating that rods are a common feature of this phylum.

**Fig. 2.**
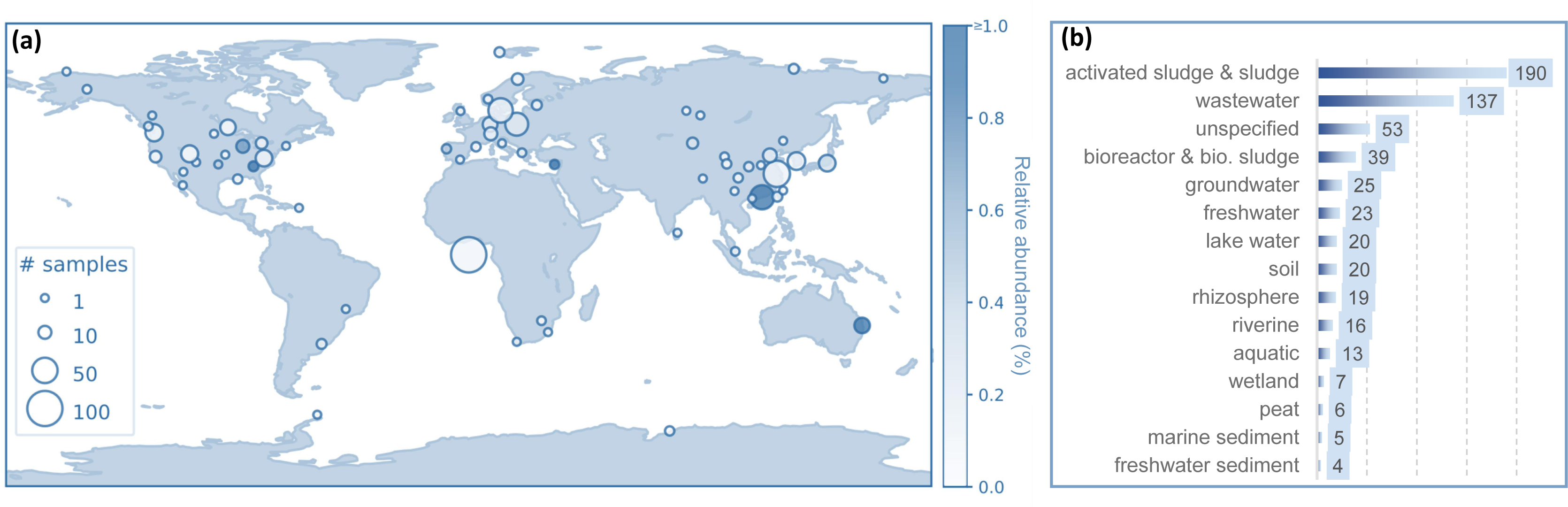
| Metabolic reconstruction of the genus *Patescibacterium gen. nov.* **(A)** Key pathways in the genus that are inferred to be present and absent. The zoom-in on the upper right shows the encoded DNA uptake machinery, including the genes PilB (assembly ATPase), PilC (inner membrane core protein), PilM (inner membrane accessory proteins), PilT (retraction ATPase), ComEC (uptake of DNA, and translocation across the inner membrane), Ssb (ssDNA binding) and RecA (catalyzes recombination with the bacterial chromosome). The presence of ComC, which is responsible for forming the pilus structure, could not be confirmed, suggesting that *Patescibacterium* gen. nov. produces only a pseudo pilus (possibly Pil A). The zoom-in on the upper left shows the cell membrane (blue) and the inferred cell wall composition consisting of murein (peptidoglycan) and rhamnose. Acronyms: GAP - Glyceraldehyde-3P; 1,3bPG - Glycerate-1,3 P_2_; 3PG - Glycerate-3P; 2PG - Glycerate-2P; PEP - Phosphoenylopyruvate; zipB - zinc and cadmium transporter; ftsX - cell division transport system permease protein; ftsE - cell division transport system ATP-binding protein.

While *Patescibacterium* lacks major carbon metabolism pathways, all ten MAGs in this genus possess a complete operon for an intermembrane F-type H+/Na+-transporting ATPase (**Fig. 2, Table S8**). A glycolysis pathway from glucose to pyruvate is encoded in all MAGs, except for phosphofructokinase-1 (PFK-1), the enzyme that catalyzes the phosphorylation of fructose-6-phosphate to fructose-1,6-bisphosphate and is considered a key regulatory step in glycolysis (**Fig. 2, Table S9**). No genes were detected for a subsequent conversion of pyruvate to acetyl-CoA nor for a fermentation to alcohol, however fermentation to lactate might be possible based on the encoded D-lactate dehydrogenase (ldhA). Prevalent was the encoded "one-carbon pool by folate", in which folate (vitamin B9) acts as a carrier of methyl groups in the form of methyl tetrahydrofolate. Genes for several transporters were present, including a zinc and cadmium transporter, a transporter (FtsX and FtsE) playing an important role in cell division machinery, a peptide/nickel transport system substrate-binding protein, and a multiple sugar transporter system (**Fig 2**, **Table S10, S11**). We did not detect any encoded flagella biosynthesis or chemotaxis proteins, suggesting that this genus is non-motile. A comparison of inferred core functions among the 10 *Patescibacterium* genomes revealed very similar patterns, although the wastewater derived genomes formed two clusters, with slight differences in pyrimidine and purine metabolism (**Fig. S5**). Remarkably, we also detected genes for unusually long proteins, including one in ABY1^TS^ and ‘Fred.cMAG.1’ consisting of 6025 and 4939 amino acids, respectively (**Table S12**). Both protein sequences contained three full length conserved domains assigned to FhaB (COG3210), a large exoprotein involved in heme utilization or adhesion (**Fig. S6a, b**). A subsequent search for protein homology by domain architecture recovered homologue proteins with a downstream autotransporter beta-domain, characterized as mediating secretion through the outer membrane (**Fig. S7**). We did not find these large encoded proteins in the other *Patescibacterium* genomes (**Fig. S6c**), nor did we recover full length homologous among 2.1 million predicted proteins assigned to ‘Patescibacteria’ at NCBI (see *Methods*). However, 149 of these 2.1 million proteins were over 4000 amino acids in length and 82 contained one or multiple Fhab and/or a DUF2341 domains (**Fig. S8**; **Table S13**). The latter domain was annotated as a predicted component of a type IV pili-like system, and was also present in the Fred.cMAG.1 protein (**Table S13**). In conclusion, these large inferred enzymes could play a role in adhesion and/or host interactions, as reported for FahB of the bacterial genus *Bordetella* (*Pseudomonadota*), that are likely to function as a toxin delivery system targeting eukaryotic hosts (Nash et al. 2024).

Overall, the emerging picture is that of a bacterial genus that is likely reliant on glycolysis, possibly using a PFK-1 alternative, as a means of energy conservation. It likely depends on a symbiotic or at least syntrophic partner to obtain essential substrates and nutrients. Encoded transporters for DNA and peptide uptake (**Fig. 2**) and the peptide cleaving potential, including peptidases such as aminopeptidases (ampS), methionyl aminopeptidases (map) and a dipeptidase (pepE) support this conclusion **(Table S14)**. Applying a recently developed machine learning based symbiont classifier, *symcla*, we used the genomic data to predict a possible symbiotic lifestyle of *Patescibacterium* (see Methods). The prediction resulted in a median score between 0.42 and 1.21, which placed this genus in the “host-associated” category (**Fig. S9**). Within this genus, the type genome ABY1^TS^ (GCA_016699775.1) of *Patescibacterium danicum* received the highest *symcla* score and is confidently placed in the middle of the same category (**Fig. S10**).

Biogeography of *Patescibacterium* gen. nov.

The 16S rRNA genes of the two MAGs (ABY1^TS^ and ‘Fred.cMAG.1’) assigned to *Patescibacterium danicum* were identical. Comparing this sequence to the MiDAS full-length 16S rRNA gene reference database, a global catalog of full length 16S rRNA gene sequences from wastewater treatment plants (WWTPs) (Dueholm et al. 2022) resulted in an assignment to the “species midas_s_17217” in the genus “midas_g_13219”. Both taxa were present in a time series across two Danish WWTPs with peaks in summer and autumn, and relative abundances of up to 0.15% (**Fig. S11)**. A comparison against a global dataset of WWTP samples revealed a prevalence of “midas_g_13219”, and the highest abundance (0.18%) of “species midas_s_17217” in a WWTP in Singapore (**Fig. S12**). These results align with a recent report that members of the phylum *Patescibacteriota* are present in WWTPs across the world (Hu et al. 2024). Our metagenomic analysis also supports the global distribution of *Patescibacterium* gen. nov., based on community profiles of 248,559 publicly available metagenomes (Woodcroft et al. 2024). In total, the genus was detected in 580 Sequence Read Archive (SRA) datasets from all seven continents (**Fig. 3a**). The most common habitat type in which *Patescibacterium* was found are aquatic, in particular wastewater and activated sludge, although it was also detected in terrestrial habitats, including soil, rhizosphere and peat (**Fig. 3b, Table S15**). In summary, our results suggest that *Patescibacterium* is globally distributed and is most commonly found in aquatic habitats, particularly engineered systems.

**Fig. 3.**
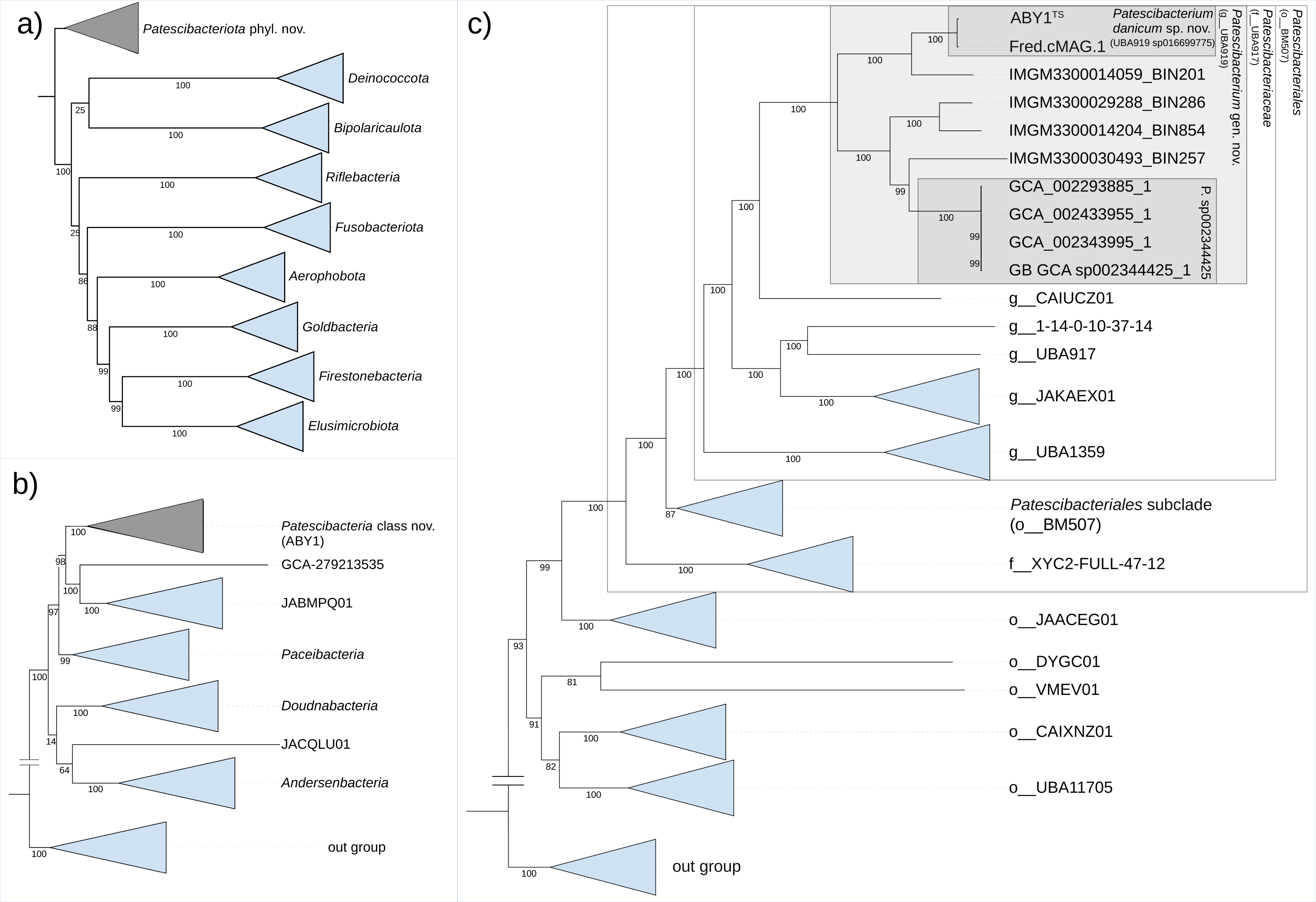
| Biogeography of *the* genus *Patescibacterium*. **(a)** Read based detection of *Patescibacterium*. A total of 580 SRA metagenomes, out of 248,559, contained hits to *Patescibacterium*, 480 of which had associated latitude/ longitude metadata and are shown here. Circle diameter indicates the number of samples per location cluster, and darker colors represent higher relative abundances (see legend). For display purposes, the abundance was capt at 1%. **(b)** Most common habitat types of *Patescibacterium* among the 580 SRA metagenomes. The list is based on the NCBI “organism” field, associated with NCBI BioSamples of metagenomic data, and has been manually curated to combine overlapping habitats. The values are counts of metagenomes per habitat. The original table is provided as **Table S15**.

### Phylogenetic diversity of *Patescibacteriota* phyl. nov.

*Patescibacteriota* was reported to comprise over 15% of the bacterial domain and to represent one of the major bacterial phyla (Schulz et al. 2017; Brown et al. 2015; Parks et al. 2018) (Coleman et al. 2021). Indeed, calculating the phylogenetic diversity (PD) (see *Methods*) on the latest bacterial GTDB reference tree, comprising over 100,000 dereplicated species, revealed *Patescibacteriota* as the fourth most diverse bacterial phylum (**Table S16**). While *Pseudomonadota* (heterotypic synonym Proteobacteria) covered most PD (18.7%), followed by Bacillota_A (12.9%) and *Bacteroidota* (10.9 %), *Patescibacteriota* were a close fourth, with a PD of 10.6% (**Table S16**). Applying PD calculations after accounting for different rates of evolutionary divergence, resulted in a slightly lower adjusted PD for *Patescibacteriota* (8.5%), possibly due to a correction of rapid evolutionary rates, that were considered to have shaped the evolutionary history of this lineage (Méheust et al. 2019). However, *Patescibacteriota* remained in fourth place, based on the adjusted PD phylum ranking (**Table S16**), confirming the status as a major bacterial phylum.

## Discussion

We proposed *Patescibacterium danicum* gen. nov., sp. nov., based on the genome ABY1^TS^ as nomenclatural type, according to the rules of the SeqCode (Whitman et al. 2024). We characterized its phylogeny, predicted its metabolic features, and mapped out its biogeographic distribution based on extant global data sets. Designating type material, i.e. nomenclatural types, is essential to establish priority of names and to ensure their uniqueness and stability (Chuvochina et al. 2019). However, names for many prokaryotic lineages were proposed without dedicated type material, i.e. a type genome, over the last decades. This malpractice has led to taxonomic instability, e.g. due to polyphyly of named lineages, resulting in an uncertain naming status for some branches in the tree of life. *Patescibacteriota* phyl. nov. has emerged as a lineage that was disproportionately affected, with over 70 names for major lineages proposed without type material (Brown et al. 2015; Anantharaman et al. 2016; Probst et al. 2018; Jaffe et al. 2020; Wrighton et al. 2014). Compounding this confusion, over a dozen of these names have since been corrected to conform with orthographic rules of nomenclature (Oren and Göker 2023). Here, we aim to contribute a modest step towards nomenclatural and taxonomic stability of this phylum by proposing the species *Patescibacterium danicum* sp. nov., the genus *Patescibacterium* gen. nov, and associated higher taxa, including the phylum *Patescibacteriota* phyl. nov., according to the SeqCode (Whitman et al. 2024).

A stable nomenclature and taxonomy for *Patescibacteriota* is desirable, since, as our results confirmed, this lineage represents one of the four major phyla in the bacterial domain. Lineages within *Patescibacteriota* have also become a popular research topic over the last decade due to their characteristic reduced genomes and host-associated lifestyles (Coleman et al. 2021), which have been experimentally verified for some members (Wang et al. 2023; Bouderka et al. 2024). *Patescibacterium danicum* has similar genome characteristics, i.e. the genome sizes of the two circular MAGs ABY1^TS^ and Fred.cMAG.1, of 939 and 917 kbp, respectively, are typical for *Patescibacteriota* genomes which have an average estimated complete genome size of 919.3 kbp ± 256 kbp, calculated based on all genomes assigned to this phylum in GTDB (**Fig. S13**). The low GC content (35.4%) of both type species MAGs also reflects the low average GC content of 33.6 ± 6.7 % of *Patescibacteriota*, which can be considered another defining feature of this lineage. Furthermore, the type species MAGs have a protein coding density (CD) of 88.17% and 86.1%, that is similar to the CD of the entire phylum (89.6 ± 2.7%), but could also indicate a symbiotic lifestyle. A recent study investigating evolutionary strategies of *Patescibacteriota*, found that the surveyed prokaryotic symbionts were characterized by a lower coding density with a median of 0.87% (Beam et al. 2020). Our machine learning-based lifestyle prediction also supports a symbiotic nature of *Patescibacterium*. However, our attempts, to calculate co-occurrence networks to predict potential partners, were inconclusive (data not shown). The search for the taxon that compensates *Patescibacterium* for its incomplete metabolic pathways is therefore still ongoing. The two long inserts in the 23S rRNA genes of ABY1^TS^ and Fred.cMAG.1, encoding a self-splicing LAGLIDADG homing endonuclease, are also a common occurrence in *Patescibacteriota*. A recent study, surveying hundreds of rRNA gene sequences assigned to this phylum, reported insertion sequences in the majority of examined 23S and nearly half of the 16S rRNA genes (Tsurumaki et al. 2024). The inserts are located in domain V of the 23S rRNA genes, i.e. insert 1 starts at position 2377 and insert 2 at position 3224 (or at 2401, when not considering insert 1) for both MAGs ABY1^TS^ and Fred.cMAG.1. Inserts in domain IV and V have been suggested to indicate that these introns are spliced (**Fig. S14**) without impacting the function of the ribosome (Tsurumaki et al. 2024), as demonstrated for 23S rRNA gene introns of hyperthermophilic bacteria of the genus *Thermotoga* (Nesbø and Doolittle 2003). In summary, the characteristics of the two MAGs assigned to *P. danicum* resemble those of a typical genome within the phylum *Patescibacteriota*. This conclusion is supported by the presence-absence pattern of predicted functions that revealed correlations of *P. danicum* and the representative of its class and most other classes in the phylum (**Fig. S15; Table S17**).

Our metabolic reconstruction suggests that the limited metabolic capacities leading to the original name patesco (Lat.), meaning bare, of *Patescibacteriota* (Rinke et al. 2013) are also a hallmark of the genus *Patescibacterium*. The absence of genes for amino acid and nucleotide biosynthesis, for the pentose phosphate pathway, the TCA cycle, and for beta oxidation, suggests a host dependent lifestyle (**Fig. 2**). This leaves the encoded glycolysis, i.e. the breakdown of glucose to pyruvate, as the most likely strategy to conserve energy. The potential to further ferment pyruvate to lactate suggests an adaptation to anaerobic habitats, which is consistent with their presence in wastewater treatment plants, bioreactor sludge and landfills (**Table S1**). Our findings are in agreement with recent studies focusing on other *Patescibacteriota* lineages, such Saccharimonadia, Parcubacteria, Gracilibacteria, and Paceibacteria (Fujii et al. 2022; Zhao et al. 2022) that also infer strongly reduced metabolic capabilities and a reliance on external nutrient sources. The remarkably large proteins (> 5000 amino acids) encoded in ABY1^TS^ and ‘Fred.cMAG.1’ that contain FhaB domains and are inferred to function as large exoproteins (FhaB) involved in adhesion, might provide first insights of how *Patescibacterium danicum* interacts with a potential host. It is tempting to speculate that these proteins are similar to FhaB homologues in lactic acid bacteria that function as cell wall-anchored proteins for the adhesion to host cells (Zhang et al. 2015). It was a surprising finding that no full-length homologues of these large proteins could be detected across all 2.1 million predicted *Patescibacteriota* proteins at NCBI, while the FhaB domain was common with one to six copies, occurring at different positions in the largest of these proteins (>4000aa). This pattern might suggest that the underlying genes undergo rapid evolution, e.g., domain reshuffling through homologous recombination. If this amounts to antigenic variation (AV), as reported for pathogens such as *Neisseria gonorrhoeae* that can replace segments of expressed genes with parts of silent genes to alter cell surface structures in order to evade the eukaryotic immune system (Voter et al. 2020), remains to be seen. Future efforts focusing on the recovery of a *P. danicum* co-culture might be able to shed light on the role of FhaB on other domains in host interactions and AV strategies.

Wastewater and activated sludge emerged as the most common *Patescibacterium* habitats in our analysis (**Fig. 3b**). Support for this finding comes from recent studies reporting that members of *Patescibacteriota* are highly abundant and diverse players in WWTP microbiomes (Kagemasa et al. 2022; Hu et al. 2024). These habitats represent nutrient-rich environments, and this could favor *Patescibacteriota* that rely on the direct uptake of nutrients, such as DNA, peptides, and sugars, from the environment. These conditions might also support syntrophic and/or symbiotic partners of this phylum, boosting the abundance of *Patescibacteriota* in return. For example, an increase in nitrogen removal rates in batch experiments coincided with an 50 times increase in relative abundances of the class *Patescibacteriia* (c_ABY1) (J. Li, Lou, and Lv 2020), suggesting that their symbiotic partners are involved in denitrification. Despite this evidence of *Patescibacteriota* being prevalent in WWTP, we cannot rule out that their relative abundances are biased by sampling procedures. For example, a groundwater study found that the average relative abundance of this phylum was nearly twice as high in a 0.1 µm filter fraction, compared to the 0.2 µm fraction that is a common standard to recover microbes from aquatic habitats (Chaudhari et al. 2021).

## Conclusion

The intention of this proposal was to provide nomenclatural and taxonomic stability by anchoring the name *Patescibacteriota* to the type genus *Patescibacterium*, in accordance with the rules of the SeqCode. In addition, we could show that MAGs assigned to this genus, and its species *P. danicum* containing the type genome ABY1^TS^, exhibit characteristics that are representative of *Patescibacteriota*, such as reduced metabolic potential and a probable host dependency. We envision that our proposal will provide a stable taxonomic foundation for further explorations of *Patescibacteriota* which have, owing to their signature trait of being small but mighty, populated most global habitats and might represent the most successful and widespread bacterial lineage on Earth.

### Description of new taxa according to the SeqCode

#### *Patescibacterium* gen. nov.

Patescibacterium (Pa.tes.ci.bac.te’ri.um. L. v. *patesco* to become visible, bare; N.L. neut. n. *bacterium* rod; N.L. neut. n. *Patescibacterium* bare bacterium).

The description of the genus is based on the single species which is characterized by a small genome size below 1 Mbp (**Table S4**) and inferred to lack nucleotide and amino acid synthesis, which is compensated by encoded DNA uptake and peptide cleaving abilities. The lack of a pentose phosphate pathway (PPP) and a complete tricarboxylic acid (TCA) cycle leaves anaerobic glycolysis, albeit without the key enzyme phosphofructokinase-1 (PFK-1), as a means of energy conservation.

The type species of the genus is *Patescibacterium danicum*, obtained from the Mariagerfjord WWTP in Denmark.

#### *Patescibacterium danicum* sp. nov.

*danicum* (da’ni.cum. N.L. neut. adj. *danicum*, Danish, named after Denmark, the country from which the type genome originates).

The species is represented by the metagenome-assembled genome (MAG) sequence ABY1^TS^, GenBank assembly accession GCA_016699775.1, obtained from metagenomic sequences from Mariagerfjord WWTP, Denmark (**Table S1**). ABY1^TS^ represents a complete, single-contig genome of 938,851 bp, with a GC content of 35.4%, a coding density of 88.17% and a predicted protein count of 890. ABY1^TS^ encodes a single copy of 5S, 16S, and 23S ribosomal RNA genes and 41 tRNAs. ABY1^TS^ meets all data quality requirements and recommendations for a MAG or SAG to serve as the nomenclatural type for a species named under the SeqCode (Hedlund et al. 2022). The metabolic predictions indicate that the organism lacks nucleotide and amino acid synthesis, which is compensated by DNA uptake and peptide cleaving abilities. The lack of a pentose phosphate pathway (PPP) and a complete tricarboxylic acid (TCA) cycle leaves anaerobic glycolysis, without the key enzyme phosphofructokinase-1 (PFK-1), as a means of energy conservation.

#### *Patescibacteriaceae* fam. nov.

*Patescibacteriaceae* (Pa.tes.ci.bac.te.ri.a.ce’ae. N.L. neut. n. *Patescibacterium* type genus of the family; -*aceae* ending to denote a family; N.L. fem. pl. n. *Patescibacteriaceae* the *Patescibacterium* family).

The family represents a monophyletic lineage based on the concatenated phylogeny of 120 protein markers and contains the genus *Patescibacterium*. The description of the family is the same as for its type genus *Patescibacterium*.

#### *Patescibacteriales* ord. nov.

*Patescibacteriales* (Pa.tes.ci.bac.te.ri.a’les. N.L. neut. n. *Patescibacterium* type genus of the order; -*ales* ending to denote an order; N.L. fem. pl. n. *Patescibacteriales* the *Patescibacterium* order).

The order represents a monophyletic lineage based on the concatenated phylogeny of 120 protein markers and contains the family *Patescibacteriaceae*. The description of the order is the same as for the family *Patescibacteriaceae*. The type genus is *Patescibacterium*.

#### *Patescibacteriia* class nov.

*Patescibacteriia* (Pa.tes.ci.bac.te.ri’ia. N.L. neut. n. *Patescibacterium* type genus of the class; -*ia* ending to denote a class; N.L. fem. pl. n. *Patescibacteriia* the *Patescibacterium* class).

The class represents a monophyletic lineage based on the concatenated phylogeny of 120 protein markers and contains the order *Patescibacteriales*. The description of the class is the same as for the order *Patescibacteriales.* The type genus is *Patescibacterium*.

#### *Patescibacteriota* phyl. nov.

*Patescibacteriota* (Pa.tes.ci.bac.te.ri.o’ta. N.L. neut. n. *Patescibacterium* type genus of the phylum; -*ota*, ending to denote a phylum; N.L. neut. pl. n. *Patescibacteriota* the *Patescibacterium* phylum).

The phylum represents a monophyletic lineage based on the concatenated phylogeny of 120 protein markers and contains the class *Patescibacteriia*. This phylum is characterized by members with a low genomic GC content (33.6 ± 6.7%), reduced genome sizes (919.3 kbp ± 256 kbp) and limited metabolic capabilities, that are likely to rely on symbiotic partners for survival. The type genus is *Patescibacterium*.

## Data availability

The MAG recovered in this study Fred.cMAG.1 has been submitted to ENA under the Project ID PRJEB78996, and was assigned the accession number GCA_964214775.1. The other nine MAGs are available via NCBI (https://www.ncbi.nlm.nih.gov/datasets/genome/), including ABY1^TS^ (GCA_016699775.1), GCA_002344425.1, GCA_002433955.1, GCA_002293885.1, GCA_002343995.1, or at IMG/M (https://img.jgi.doe.gov/cgi-bin/m/main.cgi), including IMGM3300014059, IMGM3300014204, IMGM3300029288, and IMGM3300030493. The phylogenetic tree shown in Figure 2 is provided as a newick file, under Suppl. Material.

## Acknowledgements

We thank Hermann Schwärzler and the ZID team for the support on the LEO5 compute cluster at the University of Innsbruck.

## Author contributions

CR, PH, and TW designed the project. ZD inferred phylogeny and reconstructed the metabolic features. CMS, MS and MA performed genome recovery. JCV and FS performed lifestyle predictions. AM provided phylogenetic diversity calculations. MC evaluated the proposed nomenclature. PHN and CMS contributed SSU based biogeography. CR and FS contributed the metagenome based biogeography. CR and ZD prepared the manuscript. CMS, MS, MC, TW, PHN, and PH edited and reviewed the manuscript. All authors approved the manuscript. CR supervised the study.

