## Supplementary Tables and Figures for "Proposal of *Patescibacterium danicum* gen. nov., sp. nov. in the ubiquitous ultrasmall bacterial phylum *Patescibacteriota* phyl. nov."

**Table S1** | Metadata associated with all MAGs analyzed in this study. Abbreviations: WWTP = Wastewater Treatment Plant.

| Genome sample ID | Assembly Name | Bioproject | Biosample | Country: site; habitat | Date (study) |
|---|---|---|---|---|---|
| ABY1 (GCA_016699775.1) | ASM1669977v1 | PRJNA629478 | SAMN16426283 | Denmark: Mariagerfjord WWTP | 2021-01-19 (Singleton et al. 2021) |
| Fred.cMAG.1 (GCA_964214775.1) | NA | PRJEB78996 | SAMEA115925738 | Denmark: FredericiaWWTP | This study |
| GCA_002344425.1 | ASM234442v1 | PRJNA348753 | SAMN06450679 | Australia: Thorneside WWTP | 2017-09-22 |
| GCA_002433955.1 | ASM243395v1 | PRJNA348753 | SAMN06454964 | Australia: St Lucia; bioreactor sludge* | 2017-10-04 |
| GCA_002293885.1 | ASM229388v1 | PRJNA348753 | SAMN06453158 | Australia: St Lucia; bioreactor sludge** | 2017-09-12 |
| GCA_002343995.1 | ASM234399v1 | PRJNA348753 | SAMN06450909 | Australia: Thorneside, WWTP | 2017-09-22 |
| IMGM3300014059_BIN201 | NA | PRJNA236430 | SAMN02596448 | China: Quyang, Shanghai; Engineered Wastewater, Activated Sludge | This study |
| IMGM3300014204_BIN854 | NA | PRJNA404446 | SAMN07630781 | Canada: Ontario; Engineered Solid waste, Landfill | This study |
| IMGM3300029288_BIN286 | NA | PRJNA518428 | SAMN10863920 | Canada: Ontario; Engineered Solid waste, Landfill | This study |
| IMGM3300030493_BIN257 | NA | PRJNA366145 | SAMN06268378 | United States: Espanola, New Mexico; Environmental Aquatic Groundwater Deep subsurface | This study |

* Genome binned from sequencing reads available in SRX1077115 metagenome (https://www.ncbi.nlm.nih.gov/biosample/SAMN03785553)

** Genome binned from sequencing reads available in SRX1113225 metagenome (https://www.ncbi.nlm.nih.gov/biosample/SAMN03785571)

**Table S2 | Taxonomic assignments of all ten MAGs by GTDB-tk**. Shown are the columns 'Genome ID', and 'taxonomic classification' from the GTDB-tk output. $^{TS}$ = designated type genome of the species.

| **Genome_ID** | **classification** |
|---|---|
| ABY1$^{TS}$ (GCA_016699775.1) | d__Bacteria;p__Patescibacteria;c__ABY1;o__BM507;f__UBA917;g__UBA919;s__ UBA919 sp016699775 |
| Fred_cMAG_1 | d__Bacteria;p__Patescibacteria;c__ABY1;o__BM507;f__UBA917;g__UBA919;s__ UBA919 sp016699775 |
| GCA_002293885_1 | d__Bacteria;p__Patescibacteria;c__ABY1;o__BM507;f__UBA917;g__UBA919;s__ UBA919 sp002344425 |
| GCA_002343995_1 | d__Bacteria;p__Patescibacteria;c__ABY1;o__BM507;f__UBA917;g__UBA919;s__ UBA919 sp002344425 |
| GCA_002344425 | d__Bacteria;p__Patescibacteria;c__ABY1;o__BM507;f__UBA917;g__UBA919;s__ UBA919 sp002344425 |
| GCA_002433955_1 | d__Bacteria;p__Patescibacteria;c__ABY1;o__BM507;f__UBA917;g__UBA919;s__ UBA919 sp002344425 |
| IMGM3300014059_BIN201 | d__Bacteria;p__Patescibacteria;c__ABY1;o__BM507;f__UBA917;g__UBA919;s__ |
| IMGM3300014204_BIN854 | d__Bacteria;p__Patescibacteria;c__ABY1;o__BM507;f__UBA917;g__UBA919;s__ |
| IMGM3300029288_BIN286 | d__Bacteria;p__Patescibacteria;c__ABY1;o__BM507;f__UBA917;g__UBA919;s__ |
| IMGM3300030493_BIN257 | d__Bacteria;p__Patescibacteria;c__ABY1;o__BM507;f__UBA917;g__UBA919;s__ |

**Table S3 | Genome comparison based on average nucleotide identity (ANI) values.** ANI values were calculated with fastANI (see Methods). Abbreviations: Count = count of bidirectional fragment mappings.

| QUERY_GENOME | REFERENCE_GENOME | ANI value | Count | total query fragments |
|---|---|---|---|---|
| Fred_cMAG_1.fna | Fred_cMAG_1.fna | 100 | 305 | 305 |
| Fred_cMAG_1.fna | GCA_016699775.fna | 97.5914 | 284 | 305 |
| Fred_cMAG_1.fna | IMGM3300014059_BIN201.fna | 79.5063 | 93 | 305 |
| GCA_002343995_1.fna | GCA_002343995_1.fna | 100 | 237 | 238 |
| GCA_002343995_1.fna | GCA_002344425.fna | 99.8381 | 217 | 238 |
| GCA_002343995_1.fna | GCA_002433955_1.fna | 99.7633 | 215 | 238 |
| GCA_002343995_1.fna | GCA_002293885_1.fna | 99.6336 | 192 | 238 |
| GCA_002433955_1.fna | GCA_002433955_1.fna | 99.9999 | 229 | 231 |
| GCA_002433955_1.fna | GCA_002343995_1.fna | 99.9124 | 213 | 231 |
| GCA_002433955_1.fna | GCA_002344425.fna | 99.7531 | 216 | 231 |
| GCA_002433955_1.fna | GCA_002293885_1.fna | 99.7045 | 189 | 231 |
| IMGM3300014059_BIN201.fna | IMGM3300014059_BIN201.fna | 100 | 270 | 274 |
| IMGM3300014059_BIN201.fna | Fred_cMAG_1.fna | 79.0085 | 95 | 274 |
| IMGM3300014059_BIN201.fna | GCA_016699775.fna | 78.341 | 102 | 274 |
| IMGM3300029288_BIN286.fna | IMGM3300029288_BIN286.fna | 100 | 299 | 300 |
| IMGM3300029288_BIN286.fna | IMGM3300014204_BIN854.fna | 80.6117 | 173 | 300 |
| IMGM3300029288_BIN286.fna | GCA_016699775.fna | 77.0519 | 55 | 300 |
| GCA_002293885_1.fna | GCA_002293885_1.fna | 100 | 197 | 202 |
| GCA_002293885_1.fna | GCA_002343995_1.fna | 99.9062 | 192 | 202 |
| GCA_002293885_1.fna | GCA_002344425.fna | 99.8468 | 187 | 202 |
| GCA_002293885_1.fna | GCA_002433955_1.fna | 99.7739 | 185 | 202 |
| GCA_002344425.fna | GCA_002344425.fna | 99.9999 | 233 | 234 |
| GCA_002344425.fna | GCA_002343995_1.fna | 99.8546 | 215 | 234 |
| GCA_002344425.fna | GCA_002433955_1.fna | 99.739 | 215 | 234 |
| GCA_002344425.fna | GCA_002293885_1.fna | 99.4686 | 190 | 234 |
| GCA_016699775.fna | GCA_016699775.fna | 100 | 312 | 312 |
| GCA_016699775.fna | Fred_cMAG_1.fna | 97.7155 | 282 | 312 |
| GCA_016699775.fna | IMGM3300014059_BIN201.fna | 78.8277 | 96 | 312 |
| GCA_016699775.fna | IMGM3300014204_BIN854.fna | 78.0659 | 50 | 312 |
| IMGM3300014204_BIN854.fna | IMGM3300014204_BIN854.fna | 100 | 313 | 314 |
| IMGM3300014204_BIN854.fna | IMGM3300029288_BIN286.fna | 80.4515 | 180 | 314 |
| IMGM3300014204_BIN854.fna | Fred_cMAG_1.fna | 77.6452 | 50 | 314 |
| IMGM3300014204_BIN854.fna | GCA_016699775.fna | 77.6436 | 56 | 314 |
| IMGM3300030493_BIN257.fna | IMGM3300030493_BIN257.fna | 100 | 136 | 140 |

**Table S4 | Inferred de novo nucleotide synthesis pathway of *Patescibacterium* gen. nov. genomes.** Shown are BlastKOALA annotations associated with the nucleotide synthesis pathway, separated into purine and pyrimidine metabolism. KO identifiers are provided for each encoded enzyme, and the total number of detected genes is given for each MAG.

| De novo nucleotide synthesis pathway | ABY1 GCA_016699775.1 | Fred_cMAG_1 | GCA_002344425.1 | GCA_002433955.1 | GCA_002293885.1 | GCA_002343995.1 | IMGM3300014059_BIN201 | IMGM3300014204_BIN854 | IMGM3300029288_BIN286 | IMGM3300030493_BIN257 |
|---|---|---|---|---|---|---|---|---|---|---|
| **Purine metabolism** | | | | | | | | | | |
| K00088 IMPDH, guaB; IMP dehydrogenase [EC:1.1.1.205] | 0 | 0 | 1 | 1 | 1 | 1 | 0 | 0 | 0 | 0 |
| K00525 E1.17.4.1A, nrdA, nrdE; ribonucleoside-diphosphate reductase alpha chain [EC:1.17.4.1] | 1 | 1 | 0 | 0 | 0 | 0 | 1 | 0 | 0 | 0 |
| K00526 E1.17.4.1B, nrdB, nrdF; ribonucleoside-diphosphate reductase beta chain [EC:1.17.4.1] | 1 | 1 | 0 | 0 | 0 | 0 | 1 | 0 | 0 | 0 |
| K00527 rtpR; ribonucleoside-triphosphate reductase (thioredoxin) [EC:1.17.4.2] | 1 | 1 | 1 | 1 | 1 | 1 | 1 | 1 | 1 | 0 |
| K00760 hprT, hpt, HPRT1; hypoxanthine phosphoribosyltransferase [EC:2.4.2.8] | 0 | 0 | 1 | 1 | 1 | 1 | 1 | 1 | 0 | 0 |
| K00939 adk, AK; adenylate kinase [EC:2.7.4.3] | 1 | 1 | 1 | 1 | 1 | 1 | 1 | 1 | 1 | 1 |
| K00940 ndk, NME; nucleoside-diphosphate kinase [EC:2.7.4.6] | 1 | 1 | 1 | 1 | 1 | 1 | 1 | 1 | 1 | 0 |
| K00948 PRPS, prsA; ribose-phosphate pyrophosphokinase [EC:2.7.6.1] | 0 | 0 | 0 | 0 | 0 | 0 | 0 | 1 | 0 | 0 |
| K01139 spoT; GTP diphosphokinase / guanosine-3',5'-bis(diphosphate) 3'-diphosphatase [EC:2.7.6.5 3.1.7.2] | 1 | 1 | 1 | 1 | 1 | 1 | 0 | 1 | 1 | 1 |
| K01515 nudF; ADP-ribose diphosphatase [EC:3.6.1.13 3.6.1.-] | 0 | 0 | 1 | 1 | 0 | 1 | 1 | 1 | 0 | 0 |
| K01519 rdgB, ITPA; XTP/dITP diphosphohydrolase [EC:3.6.1.66] | 0 | 0 | 0 | 0 | 0 | 0 | 1 | 0 | 0 | 0 |
| K01529 yjjX; inosine/xanthosine triphosphatase [EC:3.6.1.73] | 0 | 0 | 0 | 0 | 0 | 0 | 1 | 1 | 0 | 0 |
| K01756 purB, ADSL; adenylosuccinate lyase [EC:4.3.2.2] | 0 | 0 | 1 | 1 | 1 | 1 | 0 | 0 | 0 | 0 |
| K01923 purC; phosphoribosylaminoimidazole-succinocarboxamide synthase [EC:6.3.2.6] | 0 | 0 | 1 | 1 | 1 | 1 | 0 | 0 | 0 | 0 |
| K01939 purA, ADSS; adenylosuccinate synthase [EC:6.3.4.4] | 0 | 0 | 1 | 1 | 1 | 1 | 0 | 0 | 0 | 0 |
| K01951 guaA, GMPS; GMP synthase (glutamine-hydrolysing) [EC:6.3.5.2] | 0 | 0 | 1 | 1 | 1 | 1 | 1 | 0 | 0 | 0 |
| K01952 PFAS, purL; phosphoribosylformylglycinamidine synthase [EC:6.3.5.3] | 0 | 0 | 1 | 1 | 1 | 1 | 0 | 0 | 0 | 0 |
| K03651 cpdA; 3',5'-cyclic-AMP phosphodiesterase [EC:3.1.4.53] | 0 | 0 | 0 | 0 | 0 | 0 | 0 | 1 | 0 | 0 |
| K03783 punA, PNP; purine- | 0 | 0 | 0 | 0 | 0 | 0 | 0 | 1 | 0 | 1 |

| | | | | | | | | | | |
|---|---|---|---|---|---|---|---|---|---|---|
| nucleoside phosphorylase [EC:2.4.2.1] | | | | | | | | | | |
| K05873 cyaB; adenylate cyclase, class 2 [EC:4.6.1.1] | 2 | 1 | 1 | 1 | 1 | 1 | 2 | 1 | 0 | 2 |
| K09769 ymdB; 2',3'-cyclic-nucleotide 2'-phosphodiesterase [EC:3.1.4.16] | 1 | 1 | 1 | 1 | 1 | 1 | 1 | 1 | 1 | 0 |
| K15518 dgk; deoxyguanosine kinase [EC:2.7.1.113] | 0 | 0 | 0 | 0 | 0 | 0 | 0 | 0 | 0 | 1 |
| K15519 dck; deoxyadenosine/deoxycytidine kinase [EC:2.7.1.76 2.7.1.74] | 0 | 0 | 0 | 0 | 0 | 0 | 0 | 1 | 1 | 0 |
| **Pyrimidine metabolism** | | | | | | | | | | |
| K00254 DHODH, pyrD; dihydroorotate dehydrogenase [EC:1.3.5.2] | 1 | 1 | 0 | 1 | 1 | 1 | 1 | 1 | 1 | 0 |
| K00525 E1.17.4.1A, nrdA, nrdE; ribonucleoside-diphosphate reductase alpha chain [EC:1.17.4.1] | 1 | 1 | 0 | 0 | 0 | 0 | 1 | 0 | 0 | 0 |
| K00526 E1.17.4.1B, nrdB, nrdF; ribonucleoside-diphosphate reductase beta chain [EC:1.17.4.1] | 1 | 1 | 0 | 0 | 0 | 0 | 1 | 0 | 0 | 0 |
| K00527 rtpR; ribonucleoside-triphosphate reductase (thioredoxin) [EC:1.17.4.2] | 1 | 1 | 1 | 1 | 1 | 1 | 1 | 1 | 1 | 0 |
| K00560 thyA, TYMS; thymidylate synthase [EC:2.1.1.45] | 0 | 1 | 1 | 1 | 1 | 1 | 1 | 0 | 0 | 1 |
| K00940 ndk, NME; nucleoside-diphosphate kinase [EC:2.7.4.6] | 1 | 1 | 1 | 1 | 1 | 1 | 1 | 1 | 1 | 0 |
| K00943 tmk, DTYMK; dTMP kinase [EC:2.7.4.9] | 1 | 0 | 0 | 0 | 0 | 0 | 1 | 1 | 1 | 0 |
| K00945 cmk; CMP/dCMP kinase [EC:2.7.4.25] | 1 | 1 | 2 | 2 | 2 | 2 | 1 | 1 | 1 | 1 |
| K01465 URA4, pyrC; dihydroorotase [EC:3.5.2.3] | 0 | 0 | 1 | 1 | 1 | 1 | 0 | 1 | 0 | 0 |
| K01489 cdd, CDA; cytidine deaminase [EC:3.5.4.5] | 0 | 0 | 0 | 0 | 0 | 0 | 0 | 0 | 1 | 0 |
| K01493 comEB; dCMP deaminase [EC:3.5.4.12] | 1 | 0 | 1 | 1 | 1 | 1 | 0 | 1 | 2 | 0 |
| K01494 dcd; dCTP deaminase [EC:3.5.4.13] | 0 | 1 | 0 | 0 | 0 | 0 | 2 | 0 | 0 | 0 |
| K01937 pyrG, CTPS; CTP synthase [EC:6.3.4.2] | 1 | 1 | 1 | 1 | 1 | 1 | 1 | 1 | 1 | 1 |
| K09769 ymdB; 2',3'-cyclic-nucleotide 2'-phosphodiesterase [EC:3.1.4.16] | 1 | 1 | 1 | 1 | 1 | 1 | 1 | 1 | 1 | 0 |
| K15519 dck; deoxyadenosine/deoxycytidine kinase [EC:2.7.1.76 2.7.1.74] | 0 | 0 | 0 | 0 | 0 | 0 | 0 | 1 | 1 | 0 |
| K17828 pyrDI; dihydroorotate dehydrogenase (NAD+) catalytic subunit [EC:1.3.1.14] | 0 | 0 | 1 | 1 | 1 | 1 | 0 | 1 | 0 | 0 |

**Table S5 | Encoded enzymes of *Patescibacterium* gen. nov. genomes involved in DNA translocation into the cell.** Shown are BlastKOALA annotations associated with DNA translocation. KO identifiers are provided for each encoded enzyme, and the total number of hits is provided for each MAG.

| **DNA translocation** | **ABY1 (GCA_016699775.1)** | **Fred.cMAG.1/** | **GCA_002344425.1** | **GCA_002433955.1** | **GCA_002293885.1** | **GCA_002343995.1** | **IMGM3300014059_BIN201** | **IMGM3300014204_BIN854** | **IMGM3300029288_BIN286** | **IMGM3300030493_BIN257** |
|---|---|---|---|---|---|---|---|---|---|---|
| K02662 pilM; type IV pilus assembly protein PilM | 1 | 1 | 1 | 1 | 1 | 1 | 1 | 1 | 1 | 0 |
| K02653 pilC; type IV pilus assembly protein PilC | 1 | 1 | 1 | 1 | 1 | 1 | 1 | 1 | 1 | 1 |
| K02652 pilB; type IV pilus assembly protein PilB | 1 | 1 | 1 | 1 | 1 | 1 | 0 | 1 | 1 | 0 |
| K02669 pilT; twitching motility protein PilT | 1 | 1 | 1 | 1 | 1 | 1 | 1 | 1 | 1 | 0 |
| K02238 comEC; competence protein ComEC | 2 | 2 | 2 | 2 | 1 | 2 | 2 | 2 | 2 | 0 |
| K03111 ssb; single-strand DNA-binding protein | 1 | 1 | 1 | 1 | 1 | 1 | 1 | 1 | 1 | 1 |
| K03553 recA; recombination protein RecA | 1 | 1 | 1 | 1 | 0 | 1 | 1 | 1 | 1 | 1 |

**Table S6 | Inferred cell wall of *Patescibacterium* gen. nov. genomes consisting of murein (peptidoglycan) and rhamnose elements.** BlastKOALA annotations associated with bacterial cell wall formation and maintenance are shown. KO identifiers are provided for each encoded enzyme, and the total number of hits is provided for each MAG.

| **murein (peptidoglycan) and rhamnose elements** | **ABY1 (GCA_016699775.1)** | **Fred.cMAG.1/** | **GCA_002344425.1** | **GCA_002433955.1** | **GCA_002293885.1** | **GCA_002343995.1** | **IMGM3300014059_BIN201** | **IMGM3300014204_BIN854** | **IMGM3300029288_BIN286** | **IMGM3300030493_BIN257** |
|---|---|---|---|---|---|---|---|---|---|---|
| K00790 murA; UDP-N-acetylglucosamine 1-carboxyvinyltransferase [EC:2.5.1.7] | 1 | 1 | 1 | 1 | 1 | 1 | 1 | 1 | 1 | 1 |
| K00075 murB; UDP-N-acetylmuramate dehydrogenase [EC:1.3.1.98] | 1 | 1 | 1 | 1 | 1 | 1 | 1 | 1 | 1 | 0 |
| K01924 murC; UDP-N-acetylmuramate--alanine ligase [EC:6.3.2.8] | 1 | 1 | 1 | 1 | 1 | 1 | 1 | 1 | 1 | 0 |
| K01925 murD; UDP-N-acetylmuramoylalanine--D-glutamate ligase [EC:6.3.2.9] | 1 | 0 | 1 | 1 | 1 | 1 | 1 | 1 | 1 | 1 |
| K01928 murE; UDP-N-acetylmuramoyl-L-alanyl-D-glutamate--2,6-diaminopimelate ligase [EC:6.3.2.13] | 1 | 1 | 1 | 1 | 1 | 1 | 1 | 1 | 1 | 0 |
| K01929 murF; UDP-N-acetylmuramoyl-tripeptide--D-alanyl-D-alanine ligase [EC:6.3.2.10] | 1 | 1 | 1 | 1 | 1 | 1 | 1 | 1 | 1 | 1 |
| K01000 mraY; phospho-N-acetylmuramoyl-pentapeptide-transferase [EC:2.7.8.13] | 1 | 1 | 1 | 1 | 1 | 1 | 1 | 1 | 1 | 0 |
| K02563 murG; UDP-N-acetylglucosamine--N-acetylmuramyl-(pentapeptide) pyrophosphoryl-undecaprenol N-acetylglucosamine transferase [EC:2.4.1.227] | 1 | 1 | 1 | 1 | 1 | 1 | 1 | 1 | 1 | 0 |
| K00973 rfbA, rmlA, rffH; glucose-1-phosphate thymidylyltransferase [EC:2.7.7.24] | 1 | 1 | 1 | 1 | 1 | 1 | 1 | 1 | 1 | 1 |
| K01710 rfbB, rmlB, rffG; dTDP-glucose 4,6-dehydratase [EC:4.2.1.46] | 0 | 0 | 1 | 1 | 0 | 1 | 1 | 1 | 0 | 1 |
| K01790 rfbC, rmlC; dTDP-4-dehydrorhamnose 3,5-epimerase [EC:5.1.3.13] | 0 | 0 | 1 | 1 | 1 | 1 | 0 | 1 | 0 | 1 |
| K00067 rfbD, rmlD; dTDP-4-dehydrorhamnose reductase [EC:1.1.1.133] | 1 | 1 | 1 | 1 | 0 | 1 | 1 | 1 | 1 | 1 |

**Table S7 | Inferred rod shape determining proteins of *Patescibacterium* gen. nov. genomes.** BlastKOALA annotations of genes associated with shape determination are shown. KO identifiers are provided for each encoded enzyme and hits for each MAG.

| **rod shape determining proteins** | **ABY1 (GCA_016699775.1)** | **Fred.c MAG.1/** | **GCA_002344425.1** | **GCA_002433955.1** | **GCA_002293885.1** | **GCA_002343995.1** | **IMGM3300014059_BIN201** | **IMGM3300014204_BIN854** | **IMGM3300029288_BIN286** | **IMGM3300030493_BIN257** |
|---|---|---|---|---|---|---|---|---|---|---|
| K05837 rodA, mrdB; rod shape determining protein RodA | 1 | 1 | 1 | 1 | 1 | 1 | 1 | 1 | 1 | 0 |
| K03569 mreB; rod shape-determining protein MreB and related proteins | 2 | 2 | 2 | 2 | 2 | 2 | 2 | 2 | 2 | 2 |
| K03570 mreC; rod shape-determining protein MreC | 1 | 1 | 1 | 1 | 1 | 1 | 1 | 1 | 1 | 1 |

**Table S8 | The encoded operon for an intermembrane F-type H+/Na+-transporting ATPase of *Patescibacterium* gen. nov. genomes.** BlastKOALA gene annotations are shown, with KO identifiers provided for each encoded enzyme, and hits provided for each MAG.

| **F-type ATPase** | **ABY1 (GCA_016699775.1)** | **Fred.c MAG.1/** | **GCA_002344425.1** | **GCA_002433955.1** | **GCA_002293885.1** | **GCA_002343995.1** | **IMGM3300014059_BIN201** | **IMGM3300014204_BIN854** | **IMGM3300029288_BIN286** | **IMGM3300030493_BIN257** |
|---|---|---|---|---|---|---|---|---|---|---|
| K02108 ATPF0A, atpB; F-type H+-transporting ATPase subunit a | 1 | 1 | 1 | 1 | 1 | 1 | 1 | 1 | 1 | 1 |
| K02109 ATPF0B, atpF; F-type H+-transporting ATPase subunit b | 1 | 1 | 1 | 1 | 1 | 1 | 1 | 1 | 1 | 1 |
| K02110 ATPF0C, atpE; F-type H+-transporting ATPase subunit c | 1 | 1 | 1 | 1 | 1 | 1 | 1 | 1 | 1 | 1 |
| K02111 ATPF1A, atpA; F-type H+/Na+-transporting ATPase subunit alpha [EC:7.1.2.2 7.2.2.1] | 1 | 2 | 1 | 0 | 1 | 1 | 1 | 1 | 1 | 1 |
| K02112 ATPF1B, atpD; F-type H+/Na+-transporting ATPase subunit beta [EC:7.1.2.2 7.2.2.1] | 1 | 1 | 1 | 1 | 1 | 1 | 1 | 1 | 1 | 1 |
| K02113 ATPF1D, atpH; F-type H+-transporting ATPase subunit delta | 1 | 1 | 0 | 0 | 0 | 0 | 1 | 0 | 0 | 0 |
| K02114 ATPF1E, atpC; F-type H+-transporting ATPase subunit epsilon | 1 | 1 | 1 | 1 | 1 | 1 | 1 | 1 | 1 | 1 |
| K02115 ATPF1G, atpG; F-type H+-transporting ATPase subunit gamma | 1 | 1 | 1 | 1 | 1 | 1 | 1 | 1 | 1 | 0 |

**Table S9 | Encoded glycolysis pathway from glucose to pyruvate and lactate of *Patescibacterium* gen. nov. genomes.** BlastKOALA gene annotations are shown, with KO identifiers provided for each encoded enzyme, and hits provided for each MAG.

| **glycolysis pathway** | **ABY1 (GCA_016699775.1)** | **Fred.cMAG.1/** | **GCA_002344425.1** | **GCA_002433955.1** | **GCA_002293885.1** | **GCA_002343995.1** | **IMGM3300014059_BIN201** | **IMGM3300014204_BIN854** | **IMGM3300029288_BIN286** | **IMGM3300030493_BIN257** |
|---|---|---|---|---|---|---|---|---|---|---|
| K25026 glk; glucokinase [EC:2.7.1.2] | 1 | 1 | 1 | 1 | 0 | 1 | 1 | 1 | 1 | 1 |
| K15916 pgi-pmi; glucose/mannose-6-phosphate isomerase [EC:5.3.1.9 5.3.1.8] | 1 | 1 | 1 | 1 | 1 | 1 | 1 | 1 | 1 | 1 |
| K16305 fructose-bisphosphate aldolase / 6-deoxy-5-ketofructose 1-phosphate synthase [EC:4.1.2.13 2.2.1.11] | 1 | 1 | 1 | 0 | 1 | 1 | 1 | 1 | 1 | 1 |
| K0013 GAPDH, gapA; glyceraldehyde 3-phosphate dehydrogenase (phosphorylating) [EC:1.2.1.12] | 1 | 1 | 1 | 1 | 1 | 1 | 1 | 1 | 1 | 1 |
| K00927 PGK, pgk; phosphoglycerate kinase [EC:2.7.2.3] | 1 | 1 | 1 | 1 | 1 | 1 | 1 | 1 | 1 | 0 |
| K15634 gpmB; 2,3-bisphosphoglycerate-dependent phosphoglycerate mutase [EC:5.4.2.11] | 0 | 0 | 0 | 0 | 0 | 0 | 0 | 0 | 0 | 1 |
| K15633 gpmI; 2,3-bisphosphoglycerate-independent phosphoglycerate mutase [EC:5.4.2.12] | 1 | 1 | 1 | 1 | 1 | 1 | 1 | 1 | 1 | 0 |
| K01689 ENO1_2_3, eno; enolase 1/2/3 [EC:4.2.1.11] | 1 | 1 | 1 | 1 | 1 | 1 | 1 | 1 | 1 | 1 |
| K01007 pps, ppsA; pyruvate, water dikinase [EC:2.7.9.2] | 1 | 1 | 1 | 1 | 1 | 1 | 0 | 1 | 1 | 1 |

**Table S10 | Encoded transporters in *Patescibacterium* gen. nov. genomes.** Transporter list predicted by BLASTKOALA. KO identifiers are provided for each encoded enzyme, and hits for each MAG.

| **GENOME/ Transporter** | **ABY1 (GCA_016699775.1)** | **Fred.cMAG.1/** | **GCA_002344425.1** | **GCA_002433955.1** | **GCA_002293885.1** | **GCA_002343995.1** | **IMGM3300014059_BIN201** | **IMGM3300014204_BIN854** | **IMGM3300029288_BIN286** | **IMGM3300030493_BIN257** |
|---|---|---|---|---|---|---|---|---|---|---|
| **ABCB-BAC subgroup** K06147 ABCB-BAC; ATP-binding cassette, subfamily B, bacterial | 1 | 1 | 0 | 1 | 1 | 1 | 1 | 2 | 2 | 1 |
| **ABC Transporters** | | | | | | | | | | |
| **Putative multiple sugar transporter** K02027 ABC.MS.S; multiple sugar transport system substrate-binding protein | 1 | 1 | 1 | 1 | 1 | 1 | 1 | 1 | 1 | 0 |
| **Peptide and nickel transporters** K02035 ABC.PE.S; peptide/nickel transport system substrate-binding protein | 1 | 0 | 1 | 1 | 1 | 1 | 1 | 1 | 1 | 0 |
| **ABC-2 type and other transporters** | | | | | | | | | | |
| **Cell division transporter** K09811 ftsX; cell division transport system permease protein | 1 | 1 | 1 | 1 | 1 | 1 | 0 | 1 | 1 | 0 |
| **Cell division transporter** K09812 ftsE; cell division transport system ATP-binding protein | 1 | 1 | 1 | 1 | 1 | 1 | 0 | 1 | 1 | 0 |
| **Putative ABC transporter** K02004 ABC.CD.P; putative ABC transport system permease protein | 2 | 2 | 2 | 2 | 2 | 2 | 2 | 3 | 2 | 2 |
| **Putative ABC transporter** K02003 ABC.CD.A; putative ABC transport system ATP-binding protein | 1 | 1 | 1 | 1 | 1 | 1 | 1 | 2 | 1 | 0 |
| **Other transporters** | | | | | | | | | | |
| **Pores ion channels [TC:1]** K03282 mscL; large conductance mechanosensitive channel | 1 | 1 | 1 | 1 | 1 | 1 | 1 | 1 | 1 | 1 |
| **Electrochemical potential-driven transporters [TC:2]** K16267 zipB; zinc and cadmium transporter | 0 | 0 | 1 | 1 | 1 | 1 | 0 | 1 | 1 | 0 |

**Table S11 | Transporter list predicted by gapseq of *Patescibacterium* gen. nov. genomes.** Listed are 6 transporters with the highest e-value (cut off 1.28E-162) and bitscore (more than 500) Fred.cMAG.1/ ABY1.

| Predicted protein ID | Substrate | MAG |
|---|---|---|
| O68460 | Pyrophosphate | Fred.cMAG.1/ ABY1 (GCA_016699775.1) |
| Q8PYZ7 | Sodium | Fred.cMAG.1/ ABY1 (GCA_016699775.1) |
| O32220 | Cadmium, copper, iron, lead | Fred.cMAG.1/ ABY1 (GCA_016699775.1) |
| Q7A3E6 | Copper, iron, lead | Fred.cMAG.1/ ABY1 (GCA_016699775.1) |
| P11021 | Glucose | Fred.cMAG.1 |
| AMR68964 | Zinc | ABY1 (GCA_016699775.1) |

**Table S12 | Protein sequences of the two largest proteins encoded by *P. danicum* sp. nov., genomes.** Note that both proteins were annotated as “hypothetical” proteins by BLASTKOALA.

Inferred ‘Fred.cMAG.1’ protein consisting of 4939 amino acids.

```
>MOBHLK_04680 hypothetical protein_Fred
MEDGEYVRMTFEHLLENTRDISMYARPTKSDQSATIELYTIDGELISTSPLIDHEDIFRLFLRDLKTPANVFDFKIIGNVEIDFILDPPSITIEEIKNGKLVT
ISAAEVQKSIESVALVPVRIEIPELFKIEQKNTISVFWKNQDSENADNAENPEKEFTTYDENKNGLIDHIEWLAPHSGKQTFEVQYLFKALRLDSAKKN
LENIYEYISQQDEKSVSLLDGQYVQMKFENILDETRDIILFATPTHTGQSASIEVYTTEGQLVTVLENIDQADKYIISLRDLKTPTDIFDLKIIGNLDIDYIV
DPPKVVVESTTTGKIVTVSATEEQKATDLMVDIPVHIEIPELLKVDELNNIKIKWQTNGNQLVQFSGKDFDNNGYLDTVEWVIPELTTEKFEIIYISRAL
RLDSQKNKIEDIYDLVKARDQVYASLNDTEYVRVTFDTELDNTKDITIYAKPRDLDQPAYIEVYTLNNQLLAAYPPIDQDGKYRIVLNNLQTPTDTFDL
KIIGNVDIDFIIDPPANAYWVGGTGDWSDAANHWASTSNGTPGIANLPDATSNVFFDASSGSGATTITIDSNFSIGSMSISGYTGTIVQNADLTITDSG
AQSGDYSQTSAVTFTAIDPANNTFSATGSFSVTAGTFRRYTGLGISIDPFMVFDVYGLQGMKTNLSSSYKLNNDINASVTSGWNSGEGFFPVGDN
TTSYSGIFDGDNHTISNLYINSTTRNYVGLFGYSNNSIKNVSLTGVSITAIGTAGSNGSNGADGTSGVTPTAGGTGTAGTVGSALYIGGVVGYNIGSI
LNVSIAGSVTGTGGTGGTAGNAGNGGNTTSGATVAGAGGASVSNAAAVGGIVYAGGLVGYSQQGSITADSSVTVTGTGGTGGASGNSGNGGTN
SGSGIGGAGGAVGAPGAGGIAYVGGLIGWVNSGGIREAFSTGSVTATGGTGGAAGTAGNGGAGGTAAIGGVGGAGGVGGAGGASYAGGLIANN
LGTIQNGSYATGVVQATGGTGGTPSAGGIAGASATATTGTAATGGLGGAGGAGGIGYAGGLAALNSSDGVITGVASIVYATGTVTANGGSGINGQ
SGGVGGNGSSGTAHTGGTGGAGGVGGGGGSGNAGGLIGLNAGVTITTNSSTGLVTVVGGTGGVGGTGGSGGAKGSSGANGAGGASGTGGSA
NVTSSYGGGLVGDNTGIIETSYSTNAVNATGGAGGAGGVGVAGLTGIAGAAGGTGGAGEIINLGGITGRNTLAVIESYHTTGALVGIAGSGGAAGA
GGAGGTGALAGAAGTATAGAIIYVGGVVGSNSGTSILDVYSTGSVTGTSGTGGAGANGGVGATSLAGIAGSVGAIGGIAYVGGVVGYTANGEIFN
SYSTGDVTVAGGAGGAGGSGGNGGVGAASTTGGAGAAGAAGGAGGAAYGGGVVGSYLLGPITEIYNTGNITVTGGAGGNGGVGGSGGTGGT
NAIGGVGAIGGAGGTGGIAYAGGIAGQSTGVIQGISYSTANITANGGTPGDGGIGGNGGATTATFNSATAAAGGNGGAGGAGAIAYAGGIVGNNT
NIISNSYVSGDITATASNGKNGANPGAAGDGSAGGSSHVGAIGATGGAGGAGAIAYAGGIAGLNTSTITNLYIEAGSVTATSGNGGNGANGSVGGT
GGASAIGGAGGVGGGGGVGGVAYGGGVVAHNLGGIIQDDSHSTVAVIVTSGSSGTGGTGGAGASRAVGGAGGASGGAGAIAYGGGLVGLSSG
TINDSYATGSITVTGGGGGNPGSGGNGGSGTIGATGGANGGTDVAANVLLRAGGAAYAGGLLGYNTTNGIVQGVSYATGITTATGGAGGAGASG
ANGGNSSSAVAGAAGIAGISGGAGGIGVAGGLVGWNIAIVSDDSYASGNVSATGGAGGASGNAGSGGSSGAGIAGVAGVAGGTGGIGGISYGGG
VIAYNGIIGTTISTVNSAGTITSIGGAGNSGGSGGTGGTGGSTTGAIGGTGGAGGAGGAGGISYGGGIAGVNIGLIRGVTYAITTITVVGGAAGNGGS
GGNAGASGNIAGSTGATAGAGGTGGAGGATYVGGIAGDNSGVIISAAVTGSISASNKPGGNGGTGGAGGAGTLGTGGHNGGVGGAGGVGGAS
GNLFMGGLVGQGNGNISSSSSTSGTMTVSVSLGGGDGGVGGAGGASGASGRNGTPGAGGAGAVTGAVFVGGAIGHSSLADVVFDTIISSNHVV
VNADAGGSGGSGTAGTNGAIGSVGGAGGAGGITQLTALNAGGLIGDIVGAGGSIVDSFATGNITVVGDNGGSGGVGGAGGAGAASMTGGAGAIG
GAGGGGSNAYGGGLIGFATTNVYIGTSYATGNVTVTSGNGGVAGNGGAGGSGGSSSAGGAGAIGGAAGAGGISYTGGLAGFITSGSIVSSYSTGI
VQGISGAGGVSGNGGAGGTGGSGAIGGIGAAGTVGAVGGANFTGGLVGDNSGTIQSSYHIIGAVTATSGAGGVGGSGGTGGASGSITNGTAAIG
GAGAAGGLAGAVSAGGLIGRNNTVIALINSSYATTNVTSIGGAGGAGGLGGDGGDGSAGTNHIGGAGGIGGAGGSAPVTVAASLQAGGLVGQNV
LGTITSSYAQTGTVTATGGAGGASAQGGLGGTGTALDGLAGAGGAGGAGGSIYVGGLVGYTTGVIQTTSYSKTEIVATGGAGANGSDGRSGAVN
SGVAGSAGGAGGASGILYVGGLVGYTTSDVVSSYASYHAGSGRGNITTTGGVTGTGGNGGNGGSALATGGAGGAGAIGGIGGIATGSFVGGVA
GYSAVTGLFTTSYSEILTISGNEQPAGNGGNGGSGGLGIGAFNGGIGATGGAGGAGGVRSFGGLIGDNLGTIQGTSFSTVNIAVTGSTGGNGGTG
GNGGTSGSTISSGTAAVGGAGGAGGAGAPTYGGGLVGRMTGTTQLLQILDSYATGSVTVTGGTGGIGGTGGTGGDGSTGTSTHIGSAGALGGA
AGVGGEANAGGLVGLSNFTTGTSNFIHDYATGNVTANGGNAGTAGTSGNGGASRTTVAGGAGGGAVAAAAGGAAYAGGLVGNNRVTTGLSDNI
SSFATGTVTSTGGTGSTGSIGGNGANGAASSIGGAGGVGGTGGVGGEANSGGLAGYFSVTTGGSRSFGSYATGNSVVATGGTGGIGRSAGEPG
SGGSGAAGGTVGAGGAGGVGGAAYAGGLTGQMSGGVFTTSSYSTKDVTANGGISGNGATGVVGGTGGSSANGGVGGAGGATGVGGGSFAG
GLVASNTGIIGGTSYATGNVISNSGATGTPGVGGAGGVSGSISAGTAATGGAGGAGAAGGASYAGGLTANNAATGVIINSYATGNVTSLASNGAN
GATGGVGGDGSAGTGNHIGGAGGVGGAAGAGGASYSGGLIGNNISTTNTVNTSYSTGTTVTGTSGNGGTGGTGGAGGNPGSIGVGGIGGAGG
AAGAGGVLYTGGFVGYNNGTIQLHSYSTTNVVGNVGSGGTGGAGGSGVNGFPGSGGNGAVGGAGGAATAGGAGGIGYGGGFVGWNNTSRTI
KDSYTTGTLTLSGGAGGAGGAGGNGGDGGDGVTTFTGGTGGIAGASGAGTVGGIKYGGGFVGYNIGIINTINTVGNVSVTGSAGGVGGNGGNG
GNGGDGGPSGTGGVGGAQGATAAAGAGGILHAGGLAGYNFTAGLIQNSWSAGNVTTVGGAGGVAGVRGISGTGGAGGTLAGGNGGAAVAGG
AGGAGGASFVGGLIGSNLTTGSFNTSYASGTVTMTGGNGATGGTGGKGGNGGAGGSGGSGNGGTGTVGFVGGVGGVGGAVTGGGLVGDNT
GAISNAYALGSVTVTAGTPGNGGTGGDGGNGGAAGGGSAGANGNGANGGVGGAGASAFAGGLLGANATAVITNTYSIGVPTGTKTSGATGGVG
GAAGTGGTGGSNGSTGATGASSALQYVGGLTGNPTSGTYSSNYWDTTTSGKNNGVGNGADPGGMTGAITSVMKQQTTYSGWDFTILPIWNIVE
NTTYPRFNFQTLPDPPTNIVAVRGNGEATITFTPPANNGGSAITSYTVTSSPGGFTGTGGSSPITVTGLTNGTAYTFTITATNAIGTSIASATSNSVTP
ATVPDAPTIGTATGGNGQATVTFTPPGSNGGSAITSYTVTSSPGGFTATGGASPLTVTGLTNGVSYTFTVTATNAVGTSSASGTSNSVGPATIPDA
PTIGTATRGNGEATVTFTPPINNGGSAITSYTVTSSPGGFTGTGGASPITVTGLTNGTSYTFTVTATNAVGTSSASGTSNSVTPASVPGAPTIGTAT
RGNGEATITFSAPGSDGGSAITSYTVTSSPGGFTASGGSSPLTVTGLTNGTAYTFTVTATNAVGTGSASSASNSVTPATVPDAPTIGTATRGNGQA
TVTFTPPVFNGGSAITSYTVTSSPGGFTGTGGASPIVVTGLTNSVTYTFTVTATNDVGTSVASAASNPIVPATVPGAPTIGTATAGNGEATITFSAPG
SDGGSAITSYTVTSSPGGFTASGGSSPLTVTGLTNGTAYTFTVTATNAVGTGSASSASNSVTPVTVPGAPTNVSAVRGNGEATITFTPPSSDGGSA
ITSYTVTSSPGGFTGTGGSSPITVTGLTNGTTYTFTVTATNAVGTGIASSTSNSVVPATVPDAPTIGTATRGNGQAIVTFTPPVNNGGSAITSYTVTS
SPGGFTGTGGSSPITVTGLTNGTTYTFTVTATNAVGTSIASSASGGATPATVPDAPTIGTATRGNGEATITFTPPVNNGGSVITSYTVTSSPGGFTA
SGGASPLTVTGLTNGTTYTFTVTATNAVGTSSSSGTSNSVVPATVPDAPTIGTATRGNGEATITFTPPVNNGGSVITSYTVTSSPGGFTASGGASPL
TITGLTNGTTYTFTVTATNAVGTSSSSGTSNSVVPATVPDAPTIGTATRGNGEATITFTPPVNNGGSVITIYCDFKSWRLYCFGRSFANHYYWIN
```

Inferred ABY1$^{TS}$ protein consisting of 6025 amino acids.

```
>ALDGBO_04135 hypothetical protein_GCA_775
MLLRSIQKFLIIITAISLVGSYPAQAVLDSGLLLENSNASMVEITNTPVINVSENLPIIISSQELETKSLSDSPISVEVNSSNNSNSKTDNFTSSTEENISN
EENSNSTPPENPIVEPAPVIDPIIIPAPDPIDEEKVEETIEPIIVPEPDPILEENNEPKILETEKDLLPTILIQPTTTGKIITVSFLDEQKLTTPIINFPVHIEIPEI
YPVGQENKINLTWKNNEDKKIDFTVTDDDKNGLLDSMDWVVPEISAETPNHTFEVSYIFKALRLDSEKKSLENIYDYIFSQDEKFVSMADGEYVRMT
FEHLLENTRDISIYARPTKSDQSATIELYTIEGELIATSPLIDHEDTYRLFLRDLKTPANVFDFKIIGNVEIDFILDPPIITTEEIENGKLVTVSAAEVQKSIE
SVALVPVRIEIPELFKIEQKNTISVFWKNQDSENADNAVNPEKEFTTYDENQNGLIDHIEWLAPHSGKQTFEVQYLFKALRLDSAKKNLEDIYEYISQ
QDEKSVSLLDGQYVRMKFENILDETRDIILFATPTHTGQSASIEVYTTEGQLVTVLDNIDQADKYIISLRDLKTPTDIFDLKIIGNLDIDYIVDPPKVVVES
TTTGKIVTVSATEEQKATDLMVDIPVHIEIPELLKVDQLNQIKIKWQTNGNQLVHFSGKDFDNNGYLDTVEWIIPELSTEKFEIIYISRALRLDSQKNKIE
DIYDLVKARDQVYASLNDTEYVRVTFDTELDNTKDITIYAKPRDLDQLAYIEVYTLNNQFLAAYPPIDQDGKYRIVLNNLLTPTDSFDLKIIGNVDIDFII
DPPANAYWVGGTGDWSDAANHWASTSNGTPGIANLPDATSNVFFDASSGSAATTITIDSNFSIGSMSISGYTGTIVQNADLTITDSGAQSGDYSQT
SAATFTAIDPANNTFSATGSFSVTAGTFRRYTGLGISVDPFMVFDVYGLQGMKTNLSSSYKLNNAINASVTSGWNSGEGFIPVGDNTTSYSGIFDG
DNYTISNLYINSTTKNYVGLFGYSNNSIKNVSLTGVSITAIGTAGSNGSNGADGTSGATPTAGGTGTAGTVGSALYIGGVVGYNIGSILNVSIDGSVT
GTGGTGGAAGNAGNGGNTTSGGTVSGAGGASASNAVAIGGIVYAGGLVGYSQQGSITADSSVTVIGTGGTGGASGNGGTGGTNSGSGIGGAG
GAVGAPGAGGIAYVGGLIGWVNSGGIRESFATGTVTATGGTGGAAGSAGNGGNGGTAAIGGVGGAGGVGGAGGASYSGGLIGNNLGTIQNGSY
ATGVVQATGGTGGTPSAGGIAGASATATTGTAATGGLGGAGGAGGIGYAGGLVALNSSDGVITGVTSIVYATGTVTANGGSGINGQSGGVGGNG
SSGTAHTGGTGGAGGVGGGGGSGNAGGLIGLNAGVTITTNSSTGLVTVVGGTGGVGGTGGSGGAKGSSGANGAGGASGTGGSANVTSSYGG
GLVGDNTGIIETSYSTNAVNATGGAGGAGGVGVAGLTGIAGAAGGTGGAGEIINLGGITGRNTLAVIESYHTTGALVGIAGSGGAAGAGGAGGTGA
LAGAAGTATAGAIIYVGGVVGSNSGTSILDVYSTGSVTGTSGTGGAGANGGVGATSLAGIAGSVGAIGGIAYVGGVVGYTANGEIFNSYSTGDVTV
```

AGGAGGAGGAGGNGGVGAASTTGGAGAAGAAGGAGGAAYGGGVIGNYILGPITEIYNTGNITVTGGAGGNGGVGGSGGTGGTNAIGGVGAIGG
AGGTGGIAYAGGIAGQSTGVIQGISYSTANITANGGTPGNGGIGGNGGATTATFNSATAAAGGNGGAGGAGAIAYAGGIVGNNTNIISNSYVSGDIT
ATASNGKNGANPGAAGDGSAGGSSHVGAIGATGGAGGAGAIAYAGGIAGLNTSTITNLYIEAGSVTATSGNGGNGANGSVGGTGGASAIGGAGG
VGGGGGVGGVAYGGGVVAHNLGGIIQDDSHSTVAVIVTSGSSGTGGTGGAGASRAVGGAGGASGGAGAIAYGGGLVGLSSGTINDSYATGSITV
TGGGGGNPGSGGNGGSGTIGATGGANGGTDVAANVLLRAGGAAYAGGLLGYNTTNGIVQGVSYATGITTATGGAGGAGASGANGGNSSSAVA
GAAGIAGISGGAGGIGVAGGLVGWNIAIVSDDSYASGNVSATGGAGGASGNAGSGGSSGAGIAGVAGVAGGTGGIGGISYGGGVIAYNGIIGTTIS
TVNSAGTITSIGGAGNSGGSGGTGGTGGSTTGAIGGTGGAGGAGGAGGISYGGGIAGVNIGLIRGVTYAITTITVVGGAAGNGGTGGDAGASGNIA
GSTGATAGAGGTGGAGGATYVGGIVGDNSGVIISAAVTGSISASNKPGGNGGTGGAGGAGTLGTGGHNGGVGGAGGVGGASGNLFMGGLVGQ
GNGNISSSSSTSGTMTVSVSLGGGDGGVGGAGGASGASGRNGTPGAGGAGAVTGAVFVGGAIGHSSLADVVFDTIISSNHVVVNADAGGSGGA
GTAGTNGAIGSVGGAGGAGGITQLTALNAGGLIGDIVGAGGSIVDSFATGNITVVGDNGGSGGVGGAGGAGAASMTGGAGAIGGAGGGGSNAY
GGGLIGFATTNVYIGTSYATGNVTVTSGNGGVAGNGGAGGSGGSSSVGGAGAIGGAAGAGGISYTGGLAGFITSGSIVSSYSTGIVQGISGAGGV
SGNGGAGGTGGSGAIGGIGAAGTVGAVGGANFTGGLVGDNSGTIQSSYHIIGAVTATSGAGGVGGSGGTGGASGSITNGTAAIGGAGAAGGLAG
AVSAGGLIGRNNTVIALINSSYATTNVTSIGGAGGAGGLGGDGGDGSAGTNHIGGAGGIGGAGGSAPVTVAASLQAGGLVGQNVLGTITSSYAQT
GTVTATGGAGGASAQGGLGGTGTALDGLAGAGGAGGAGGSIYVGGLVGYTTGVIQTTSYSKTEIVATGGAGTNGSDGRSGAVNSGVAGSAGGA
GGASGILYVGGLVGYTTSDVVSSYASYHAGSGRGNITTTGGVTGTGGNGGNGGSALATGGAGGAGAIGGIGGIATGSFVGGVAGYSAVTGLFTT
SYSEILTISGNEQPAGNGGNGGSGGLGIGAFNGGIGATGGAGGAGGVRSFGGLIGDNLGTIQGTSFSTVNIAVTGSTGGNGGTGGNGGTSGSTIS
SGTAAVGGAGGAGGAGAPTYGGGLVGRMTGTTQLLQILDSYATGSVTVTGGTGGIGGTGGTGGDGSTGTSTHIGSAGALGGAAGVGGEANAG
GLVGLSNFTTGTSNFIHDYATGNVTANGGNAGTAGTSGNGGASRTTVAGGAGGGAVAAAAGGAAYAGGLVGNNRVTTGLSDNISSFATGTVTST
GGTGSTGSIGGNGANGAASSIGGAGGVGGTGGVGGEANSGGLAGYFSVTTGGSRSFGSYATGNSVVATGGTGGIGRSAGEPGSGGSGAAGGT
VGAGGAGGVGGAAYAGGLTGQMSGGVFTTSSYSTKDVTANGGISGNGATGVVGGTGGSSANGGVGGAGGATGVGGGSFAGGLVASNTGIIGG
TSYATGNVISNSGATGTPGVGGAGGVSGSISAGTAATGGAGGAGAAGGASYAGGLTANNAATGVIINSYATGNVTSLASNGANGATGGVGGDGS
AGTGNHIGGAGGVGGAAGAGGASYSGGLIGNNISTTNTVNTSYSTGTTVTGTSGNGGTGGTGGAGGNPGSIGVGGIGGAGGAAGAGGVLYTGG
FVGYNNGTIQLHSYSTTNVVGNVGSGGTGGAGGSGVNGFPGSGGNGAVGGAGGAATAGGAGGIGYGGGFVGWNNTSRTIKDSYTTGTLTLSG
GAGGAGGAGGNGGDGGDGVTTFTGGTGGIAGASGAGTVGGIKYGGGFVGYNIGTINTINTVGNVSVTGSAGGVGGNGGNGGNGGDGGPSGT
GGNGGAQGATAAAGAGGALHAGGLAGYNFTAGLIQNSWSAGNVTTVGGAGGVAGVRGISGTGGAGGTLAGGAGGAGITGGAGGAGGASFVG
GLIGSNLTTGSFNTSYASGTVTMTGGNGATGGTGGKGGNGGAGGSGGSGNGGAGTVGFVGGVGGVGGAVTGGGLVGDNTGTISNAYALGSVT
VTAGTPGNGGNGGDGGNGGAAGGGSAGANGNGANGGVGGAGASAFAGGLLGANATAVITNTYSIGVPTGTKTSGATGGAGGAAGTGGTGGS
AGSSGAAGASSALQYVGGLTGNPTSGTYSSNYWDTTTSGKNNGVGNGADPGGMTGAITSVMKQQATYSGWDFTILPIWNIVETSTYPRFNFQTL
PDPPTNIVAVRGNGEATITFTPPANNGGSAITSYTVTSSPGGFTGTGGSSPITVTGLTNGTAYTFTVTATNAIGTSIASATSNSVTPATVPDAPTIGTA
TGGNGQATVSFTPPINNGGSAITSYTVTSSPGGFTATGGASPLTVTGLTNGVSYTFTVTATNAVGTSSASGTSNSVGPATIPDAPTIGTATRGNGE
ATVTFTPPINNGGSAITSYTVTSSPGGFTGTGGASPITVTGLTNGTSYTFTVTATNAVGTSSASGTSNSVTPASVPGAPTIGTATRGNGEATITFSAP
GSDGGSAITSYTVTSSPGGFTASGGSSPLTVTGLTNGTAYTFTVTATNAVGTGSASSASNSVTPATVPDAPTIGTATRGNGQATVTFTPPVFNGG
SAITSYTVTSSPGGFTGTGGASPIVVTGLTNSVTYTFTVTATNDVGTSVASAASNPIVPATVPGAPTIGTATAGNGEATITFSAPGSDGGSAITSYTV
TSSPGGFTASGGSSPLTVTGLTNGTAYTFTVTATNAVGTGSASSASNSVTPVTIPGAPTNVSAVRGNGEATITFTPPSSDGGSAITSYTVTSSPGG
FTGTGGSSPITVTGLTNGTTYTFTVTATNAVGTGIASSTSNSVVPATVPDAPTIGTATRGNGQAIVTFTPPVNNGGSAITSYTVTSSPGGFTGTGGS
SPITVTGLTNGTTYTFTVTATNAVGTSIASSASGGATPATVPDAPTIGTATRGNGEATITFTPPVNNGGSVITSYTVTSSPGGFTASGGASPLTITGLT
NGTTYTFTVTATNAVGTSSSSGTSNSVVPATVPDAPTIGTATRGNGEATITFTPPVNNGGSVITSYTVTSSPGGFTASGGASPLTITGLTNGTTYTF
TVTATNAVGTSSSSGTSNSVVPATVPDAPTIGTATRGNGEATITFTPPVNNGGSVITSYTVTSSPGGLTASGGASPITITGLTNGTAYTFTVTATNAL
GTSSASLASNSVTPATVPDAPTNLAAAIGDSSATLTWTAPAFDGGSAITSYTVTSSPGGFTASGSSPLTVTGLTNGVTYTFTATATNAIGTSSPSGT
SNSVTPASVPDVPTNLAATIDDSSVDLSWTAPVSNGGSPITDYVVQYKLTVDGTWATFADGVSTTPATTVINLSNNNSYDFRVSAKNLVGTGSPSV
SVSATPGSPAQVLIQSFPSLVVPNIGTAVRITNEGSTAYEYQYTWCVTTADNNFCGGGDDIFSSTAAKLIQPAENFDTTLNSTVPTAGNYWFHISVQ
FGSESSEAAQSFTATGTGGGGGGGGGGGGGGGGTPPDITPPTNVSVTIAAGVAVINTTSVVLTMTAIGANYMMISDNSAFTGATWETYVTSKP
WFLSPGNGVKNVFVKFKDLAGNESLVVSDSITLAIDSVTPPGNQRAIDVADDAVALTRILGIERHTNWEADNISRVIAIAQQSGVIINNYDLTVAVNFV
TYGVSDVTIRQGWVERLKLVGDQLQTLGYINVYALNEISDGVKPNKRNVVYEQKQFSAVSQAFRTLVGRNYNPADLTDDLAWHTMMYRLRFPRNI
VKENAAVAHYIKVFKRKPVSAFDWSVVRAWAYVLSGRGQVIIKETGNILVIKSTPTGWLRVRSAPSLTGTIMGSVYPNQRYTFSQEKNGWFNITISG
NRKGWVSGQYVTVVSRATPEKITIPEPKKIIVPKAKTVTVQATPTGWLNVRSIPSTQGTIRTRVYPSQTYVYTKYEQGWYLITLNNGTTGWVTEQYV
K

**Table S13 | Predicted long proteins containing the Fhab and MJ1470 domains**. Out of the investigated 149 long protein sequences assigned to the taxonomy "Patescibacteria group", obtained from NCBI, 82 contained a Fhab and/or a MJ1470 domain. Overall, 75 contain Fhab, of which 59 contained only Fhab, and 7 contained only MJ1470 (COG5306) The latter is annotated as "uncharacterized conserved protein MJ1470, contains DUF2341 domain, predicted component of type IV pili-like system" at NCBI (https://www.ncbi.nlm.nih.gov/research/cog/cog/COG5306/).

| Protein ID | BioProject | Fhab<br>Large exoprotein involved in heme utilization or adhesion [Intracellular trafficking, secretion, and vesicular transport] | MJ1470/ DUF2341<br>Uncharacterized conserved protein MJ1470, contains DUF2341 domain, predicted component of type IV pili-like system [General function prediction only] |
|---|---|---|---|
| **GIW59270** | PRJDB11609 | 6 | 0 |
| **OHA62620** | PRJNA288027 | 5 | 0 |
| **KND49831** | PRJNA283788 | 4 | 0 |
| **PCI28011** | PRJNA391950 | 4 | 0 |
| **PIZ79246** | PRJNA362739 | 4 | 0 |
| **PJA68901** | PRJNA362739 | 4 | 0 |
| **PJC42213** | PRJNA362739 | 4 | 0 |
| **QQG53011** | PRJNA640378 | 4 | 0 |
| **USN92160** | PRJNA648801 | 3 | 7 |
| **USN88639** | PRJNA648801 | 3 | 6 |
| **KKU23721** | PRJNA273161 | 3 | 1 |
| **AKQ02518** | PRJNA262935 | 3 | 0 |
| **KKQ53758** | PRJNA273161 | 3 | 0 |
| **KKQ61136** | PRJNA273161 | 3 | 0 |
| **KKQ63205** | PRJNA273161 | 3 | 0 |
| **KKQ65401** | PRJNA273161 | 3 | 0 |
| **KKQ69978** | PRJNA273161 | 3 | 0 |
| **KKQ72541** | PRJNA273161 | 3 | 0 |
| **KND46770** | PRJNA283788 | 3 | 0 |
| **OGC80521** | PRJNA288027 | 3 | 0 |
| **OGC87865** | PRJNA288027 | 3 | 0 |
| **OGF37239** | PRJNA288027 | 3 | 0 |
| **OGF39440** | PRJNA288027 | 3 | 0 |
| **OGF41578** | PRJNA288027 | 3 | 0 |
| **OGG53755** | PRJNA288027 | 3 | 0 |
| **OGK62849** | PRJNA288027 | 3 | 0 |
| **OGK64048** | PRJNA288027 | 3 | 0 |
| **OGK65894** | PRJNA288027 | 3 | 0 |
| **OGK70299** | PRJNA288027 | 3 | 0 |
| **OGK74041** | PRJNA288027 | 3 | 0 |
| **OGY60217** | PRJNA288027 | 3 | 0 |
| **OGY60473** | PRJNA288027 | 3 | 0 |
| **OGY62004** | PRJNA288027 | 3 | 0 |
| **QQS60272** | PRJNA629478 | 3 | 0 |
| **QSH39715** | PRJNA701838 | 3 | 0 |
| **TSC88693** | PRJNA387583 | 3 | 0 |
| **USN87822** | PRJNA648801 | 3 | 0 |
| **WKZ27142** | PRJNA983097 | 3 | 0 |
| **WKZ28703** | PRJNA983097 | 3 | 0 |
| **KKU23075** | PRJNA273161 | 2 | 1 |
| **OGG71089** | PRJNA288027 | 2 | 1 |
| **OGG85015** | PRJNA288027 | 2 | 1 |
| **OGY83032** | PRJNA288027 | 2 | 1 |
| **KKP83066** | PRJNA273161 | 2 | 0 |
| **KKQ13942** | PRJNA273161 | 2 | 0 |

| | | | |
|---|---|---|---|
| **KKQ17080** | PRJNA273161 | 2 | 0 |
| **KKR12461** | PRJNA273161 | 2 | 0 |
| **KKS38360** | PRJNA273161 | 2 | 0 |
| **KKT43414** | PRJNA273161 | 2 | 0 |
| **KKT97993** | PRJNA273161 | 2 | 0 |
| **KPJ85734** | PRJNA270657 | 2 | 0 |
| **OGI27043** | PRJNA288027 | 2 | 0 |
| **OGZ68747** | PRJNA288027 | 2 | 0 |
| **PLX27174** | PRJNA387015 | 2 | 0 |
| **QHO62982** | PRJNA414521 | 2 | 0 |
| **TAK04260** | PRJNA514088 | 2 | 0 |
| **TSC87231** | PRJNA387583 | 2 | 0 |
| **UMX47940** | PRJNA802382 | 2 | 0 |
| **KKT80597** | PRJNA273161 | 1 | 2 |
| **OGF36494** | PRJNA288027 | 1 | 2 |
| **OGF38657** | PRJNA288027 | 1 | 2 |
| **OGF40974** | PRJNA288027 | 1 | 2 |
| **KKQ54817** | PRJNA273161 | 1 | 1 |
| **KKQ64009** | PRJNA273161 | 1 | 1 |
| **KKQ66643** | PRJNA273161 | 1 | 1 |
| **KKQ71114** | PRJNA273161 | 1 | 1 |
| **QQR65269** | PRJNA629478 | 1 | 1 |
| **KKU92066** | PRJNA273161 | 1 | 0 |
| **OGG19417** | PRJNA288027 | 1 | 0 |
| **OGI87241** | PRJNA288027 | 1 | 0 |
| **OGL60671** | PRJNA288027 | 1 | 0 |
| **OGL70302** | PRJNA288027 | 1 | 0 |
| **OGL82989** | PRJNA288027 | 1 | 0 |
| **OGL84435** | PRJNA288027 | 1 | 0 |
| **OQB18202** | PRJNA321808 | 1 | 0 |
| **KKQ54234** | PRJNA273161 | 0 | 2 |
| **KKQ63270** | PRJNA273161 | 0 | 2 |
| **KKQ65612** | PRJNA273161 | 0 | 2 |
| **KKQ70002** | PRJNA273161 | 0 | 2 |
| **KKQ72757** | PRJNA273161 | 0 | 2 |
| **OGF26850** | PRJNA288027 | 0 | 2 |
| **PLX27049** | PRJNA387015 | 0 | 2 |

**Table S14 | Inferred peptide cleaving potential of *Patescibacterium* gen. nov. genomes.** BlastKOALA annotations included peptidases such as aminopeptidases (ampS), methionyl aminopeptidases (map) and dipeptidases (pepE). KO identifiers are provided for each encoded enzyme, and hits for each MAG.

| **Inferred peptide cleaving potential** | **ABY1 (GCA_016699775.1)** | **Fred.cMAG.1/** | **GCA_002344425.1** | **GCA_002433955.1** | **GCA_002293885.1** | **GCA_002343995.1** | **IMGM3300014059_BIN201** | **IMGM3300014204_BIN854** | **IMGM3300029288_BIN286** | **IMGM3300030493_BIN257** |
|---|---|---|---|---|---|---|---|---|---|---|
| K19689 ampS, pepS, ampT; aminopeptidase [EC:3.4.11.-] | 0 | 0 | 1 | 1 | 1 | 1 | 0 | 0 | 0 | 0 |
| K01265 map; methionyl aminopeptidase [EC:3.4.11.18] | 1 | 1 | 1 | 1 | 1 | 1 | 1 | 1 | 1 | 1 |
| K05995 pepE; dipeptidase E [EC:3.4.13.21] | 0 | 0 | 0 | 0 | 0 | 0 | 2 | 2 | 1 | 0 |

**Table S15 | Habitat types.** Original list of the NCBI “organism” field, associated with NCBI BioSamples of metagenomic data. Data were obtained from sandpiper v0.3.0 (see Methods). NCBI labels were simplified (see column “Simplified labels”) and the following were later combined: activated sludge & sludge and bioreactor & bioreactor sludge.

| Simplified labels | NCBI “organism” | Count |
|---|---|---|
| wastewater | wastewater metagenome | 137 |
| activated sludge | activated sludge metagenome | 130 |
| unspecified | metagenome | 53 |
| sludge | sludge metagenome | 35 |
| bioreactor | bioreactor metagenome | 25 |
| groundwater | groundwater metagenome | 25 |
| freshwater | freshwater metagenome | 23 |
| lake water | lake water metagenome | 20 |
| soil | soil metagenome | 20 |
| rhizosphere | rhizosphere metagenome | 19 |
| riverine | riverine metagenome | 16 |
| bioreactor sludge | bioreactor sludge metagenome | 14 |
| aquatic | aquatic metagenome | 13 |
| wetland | wetland metagenome | 7 |
| peat | peat metagenome | 6 |
| marine sediment | marine sediment metagenome | 5 |
| freshwater sediment | freshwater sediment metagenome | 4 |
| landfill | landfill metagenome | 4 |
| marine | marine metagenome | 4 |
| biofilter | biofilter metagenome | 3 |
| permafrost | permafrost metagenome | 3 |
| salt lake | salt lake metagenome | 3 |
| biofilm | biofilm metagenome | 2 |
| environmental samples | environmental samples | 2 |
| gut | gut metagenome | 2 |
| anaerobic digester | anaerobic digester metagenome | 1 |
| cave | cave metagenome | 1 |
| cold seep | cold seep metagenome | 1 |
| cold spring | cold spring metagenome | 1 |
| fish | fish metagenome | 1 |
| uncultured bacterium | uncultured bacterium | 1 |
| wood decay | wood decay metagenome | 1 |

**Table S16 | Phylogenetic diversity (PD) of the top 50 bacterial phyla in GTDB.** The PD was calculated on the latest bacterial R220 GTDB reference tree, and that PD value is shown in the column “standard_pd”. After accounting for different rates of evolutionary divergence (see Methods), the PD was recalculated and this adjusted PD value is shown in the column “adjusted_pd”. *Patescibacteriota* phyl. nov. is highlighted in **bold**.

| Phylum | standard_pd | adjusted_pd |
|---|---|---|
| p__Pseudomonadota | 18.703 | 18.1896 |
| p__Bacillota_A | 12.8671 | 11.3428 |
| p__Bacteroidota | 10.9206 | 9.5019 |
| **p__*Patescibacteriota phyl. nov.*** | **10.5838** | **8.4943** |
| p__Actinomycetota | 6.8158 | 7.0201 |
| p__Bacillota_I | 3.6072 | 3.9408 |
| p__Chloroflexota | 3.6878 | 3.7858 |
| p__Planctomycetota | 3.6578 | 3.4727 |
| p__Desulfobacterota | 2.558 | 3.2217 |
| p__Acidobacteriota | 2.0499 | 2.4183 |
| p__Verrucomicrobiota | 2.7563 | 2.3824 |
| p__Bacillota | 1.8494 | 2.3143 |
| p__Spirochaetota | 1.6476 | 1.9707 |
| p__Cyanobacteriota | 1.3583 | 1.7081 |
| p__Myxococcota | 1.4758 | 1.5414 |
| p__Omnitrophota | 1.1661 | 1.365 |
| p__Bdellovibrionota | 1.1257 | 1.0403 |
| p__Campylobacterota | 0.8331 | 0.9333 |
| p__Bacillota_B | 0.6151 | 0.9258 |
| p__Nitrospirota | 0.5434 | 0.7153 |
| p__Gemmatimonadota | 0.6078 | 0.6837 |
| p__Armatimonadota | 0.6036 | 0.6628 |
| p__Elusimicrobiota | 0.5127 | 0.5425 |
| p__Bacillota_C | 0.383 | 0.5358 |
| p__Chlamydiota | 0.6766 | 0.5062 |
| p__Bacillota_E | 0.3124 | 0.434 |
| p__Marinisomatota | 0.378 | 0.4226 |
| p__Bacillota_G | 0.252 | 0.4209 |
| p__WOR-3 | 0.3127 | 0.3552 |
| p__Dependentiae | 0.3434 | 0.3192 |
| p__Zixibacteria | 0.2342 | 0.3103 |
| p__Synergistota | 0.2566 | 0.3034 |
| p__Margulisbacteria | 0.2444 | 0.2917 |
| p__Eremiobacterota | 0.2332 | 0.2832 |
| p__Desulfobacterota_B | 0.1995 | 0.2637 |
| p__Bacillota_D | 0.1713 | 0.2446 |
| p__Cloacimonadota | 0.2044 | 0.2397 |

| | | |
|---|---|---|
| p__Fibrobacterota | 0.1976 | 0.2293 |
| p__Deinococcota | 0.1783 | 0.2101 |
| p__Nitrospinota | 0.151 | 0.1995 |
| p__Myxococcota_A | 0.1667 | 0.1921 |
| p__KSB1 | 0.114 | 0.1822 |
| p__Thermotogota | 0.1563 | 0.1798 |
| p__Krumholzibacteriota | 0.151 | 0.1791 |
| p__UBA10199 | 0.1329 | 0.1774 |
| p__Bipolaricaulota | 0.1489 | 0.1731 |
| p__Fusobacteriota | 0.1164 | 0.1681 |
| p__Hydrogenedentota | 0.1286 | 0.1669 |
| p__Aquificota | 0.1487 | 0.1638 |
| p__Latescibacterota | 0.1071 | 0.1514 |

**Table S17 | Genome representatives of the 25 Patescibacteriota classes present in GTDB r214.** The representatives were selected according to criteria proposed in *Parks et al. 2018**.

| CLASS | REPRESENTATIVE GENOME R214 |
|---|---|
| C__PACEIBACTERIA | GCA_000995965 |
| C__KAZAN-3B-28 | GCA_001029795 |
| C__GRACILIBACTERIA | GCA_001430755 |
| C__CPR3 | GCA_001771135 |
| C__ANDERSENBACTERIA | GCA_001817055 |
| C__ABY1 | GCA_001818315 |
| C__GCA-2792135 | GCA_002792135 |
| C__CPR2_A | GCA_002792735 |
| C__DOJKABACTERIA | GCA_002840365 |
| C__SACCHARIMONADIA | GCA_004138405 |
| C__SOKK01 | GCA_004374725 |
| C__SICC01 | GCA_009694835 |
| C__JACMRA01 | GCA_014377045 |
| C__JAHJAL01 | GCA_018814055 |
| C__JABMPQ01 | GCA_018818205 |
| C__4484-211 | GCA_018824115 |
| C__DYJS01 | GCA_020723005 |
| C__UBA1384 | GCA_021157605 |
| C__JACPGU01 | GCA_022703645 |
| C__CPR2 | GCA_023473595 |
| C__DOUDNABACTERIA | GCA_023484515 |
| C__MICROGENOMATIA | GCA_024277325 |
| C__CG2-30-54-11 | GCA_903889905 |
| C__WWE3 | GCA_943354835 |
| C__JAEDAM01 | GCF_021057185 |

* Parks, Donovan H., Maria Chuvochina, David W. Waite, Christian Rinke, Adam Skarshewski, Pierre-Alain Chaumeil, and Philip Hugenholtz. 2018. 'A Standardized Bacterial Taxonomy Based on Genome Phylogeny Substantially Revises the Tree of Life'. Nature Biotechnology 36 (10): 996–1004. https://doi.org/10.1038/nbt.4229.

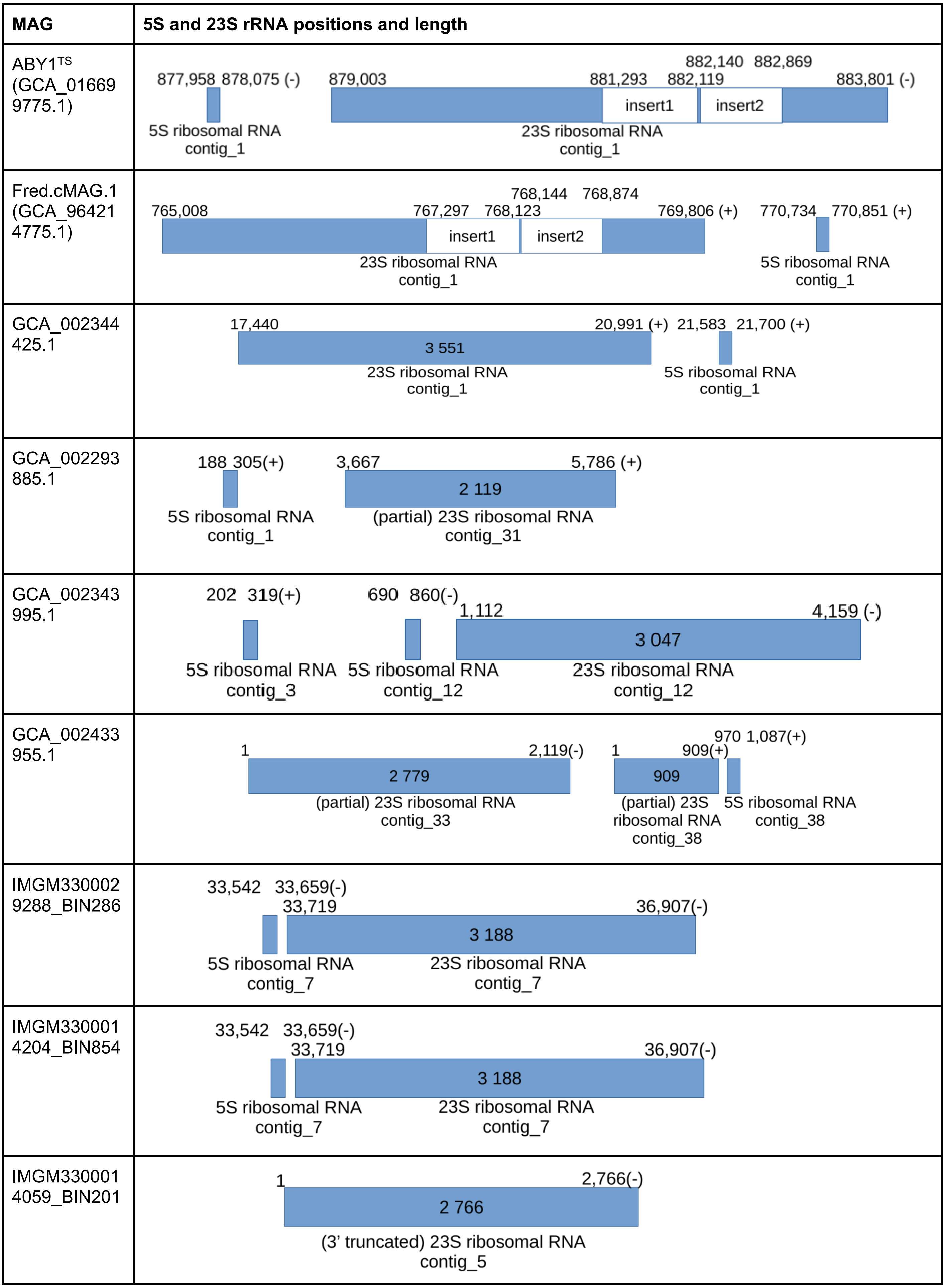
MAG
5S and 23S rRNA positions and length
ABY1TS (GCA_016699775.1)
877,958 878,075 (-)
879,003
881,293
882,119
882,140 882,869
883,801 (-)
insert1
insert2
5S ribosomal RNA contig_1
23S ribosomal RNA contig_1
Fred.cMAG.1 (GCA_964214775.1)
765,008
767,297 768,123
768,144 768,874
769,806 (+)
770,734 770,851 (+)
insert1
insert2
23S ribosomal RNA contig_1
5S ribosomal RNA contig_1
GCA_002344425.1
17,440
20,991 (+)
21,583 21,700 (+)
3 551
23S ribosomal RNA contig_1
5S ribosomal RNA contig_1
GCA_002293885.1
188 305(+)
3,667
5,786 (+)
2 119
5S ribosomal RNA contig_1
(partial) 23S ribosomal RNA contig_31
GCA_002343995.1
202 319(+)
690 860(-)
1,112
4,159 (-)
3 047
5S ribosomal RNA contig_3
5S ribosomal RNA contig_12
23S ribosomal RNA contig_12
GCA_002433955.1
1
2,119(-)
1
909(+)
970 1,087(+)
2 779
909
(partial) 23S ribosomal RNA contig_33
(partial) 23S ribosomal RNA contig_38
5S ribosomal RNA contig_38
IMGM3300029288_BIN286
33,542 33,659(-)
33,719
36,907(-)
3 188
5S ribosomal RNA contig_7
23S ribosomal RNA contig_7
IMGM3300014204_BIN854
33,542 33,659(-)
33,719
36,907(-)
3 188
5S ribosomal RNA contig_7
23S ribosomal RNA contig_7
IMGM3300014059_BIN201
1
2,766(-)
2 766
(3' truncated) 23S ribosomal RNA contig_5

**Fig. S1 | Ribosomal 5S and 23S RNA genes of all nine MAGs**. Provided are the contig coordinates for all nine MAGs containing a 23S rRNA gene. The bars representing the rRNA genes are scaled according to gene length. The rather incomplete MAG IMGM3300030493_BIN257 did not contain a 5S or 23S rRNA and is therefore not shown. Analysis of 23S ribosomal RNA genes revealed that both *Patescibacterium danicus* sp. nov. MAGs, Fred.cMAG.1 and ABY1$^{TS}$ (GCA_016699775.1), contain two large insertions. The gap between 5S and 23S is exactly 928bp in Fred and ABY1$^{TS}$, the 23S rRNA gene length is also exactly the same, i.e. 4798 bp, although both sequences differed by 5 positions: pos1934 Fred.cMAG.1(G), ABY1$^{TS}$(A); pos1935 Fred.cMAG.1(C), ABY1$^{TS}$(T); pos2429 Fred.cMAG.1(C), ABY1TS(T); pos2679 Fred.cMAG.1(G), ABY1TS(A); pos3584 Fred.cMAG.1(T), ABY1TS(C)). The 23S rRNA gene of MAG GCA_002433955.1 is split into two fragments, both of which were identified as 23S rRNA genes of *Ca.* Patescibacteria by the NCBI's BLASTN, however, no overlap was found between these fragments which are part of different contigs. The MAG GCA_002343995.1 contains two 5S rRNA genes that are located on two different contigs.

| | |
|---|---|
| **Insert1** | |
| Predicted ORF | 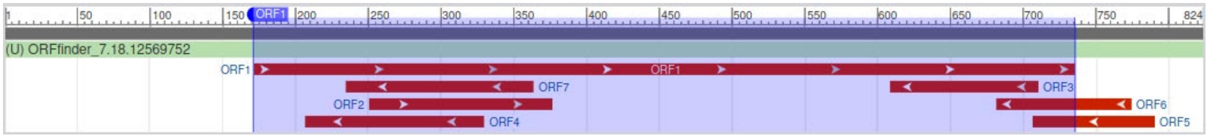 |
| Predicted protein sequence | >lcl\|ORF1<br>MIKSENVSRADNQQERLRRLMLIGWIVGFVDGEGCFSIGIFMQQDREEQN<br>RIRRGYATGYQVFHEFSVTQGEKSKSALEELQNFFQVGKLYLNKRHDNHT<br>ENLWKYVVRKRIDLINVIIPFFMENKLRTSKNLDFLKFVKCLEIINQGEH<br>LTRAGVIKIAEIAATMNRRKSRDSIIRILRDYMPNPS |
| Blast hit | LAGLIDADG family homing endonuclease (domain architecture ID 10469810)<br><br>LAGLIDADG family homing endonuclease belongs to a large family of homing endonucleases that each contain one or two copies of a motif that resembles the consensus sequence LAGLIDADG and is directly involved in the DNA cutting process |
| **Insert2** | |
| Predicted ORF | 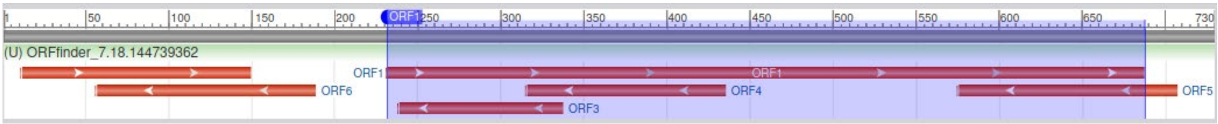 |
| Predicted protein sequence | >lcl\|ORF1<br>MDINSYITGFVDGEGCFLVSFSLRSKMKLGIEVRPSFSVSQHKCSQDIIF<br>FLQRFFRCGGVRFSNKDQNYKYEVRNITELVKIIIPHFKKYPLMTAKKDE<br>FERFAKICELIYSNHHLSKVGLEEILMLSEKLNIDGNKKYRRTDLLKMIT<br>R |
| Blast hit | LAGLIDADG family homing endonuclease (domain architecture ID 10469810)<br><br>LAGLIDADG family homing endonuclease belongs to a large family of homing endonucleases that each contain one or two copies of a motif that resembles the consensus sequence LAGLIDADG and is directly involved in the DNA cutting process |

**Fig. S2 | Analysis of inserts in 23S rRNA gene sequences of Fred.cMAG.1 and ABY1$^{TS}$ (GCA_016699775.1).** The sequences of insert 1 and insert 2 are identical in Fred.cMAG.1 and ABY1$^{TS}$. The insert identification analysis, including the prediction of open reading frames (ORFs) and subsequent blastp homology searches, identified the large ORF1 fragments as homing endonuclease, in both inserts. Several other ORFs are present, but none of them yielded specific hits in blastp searches.

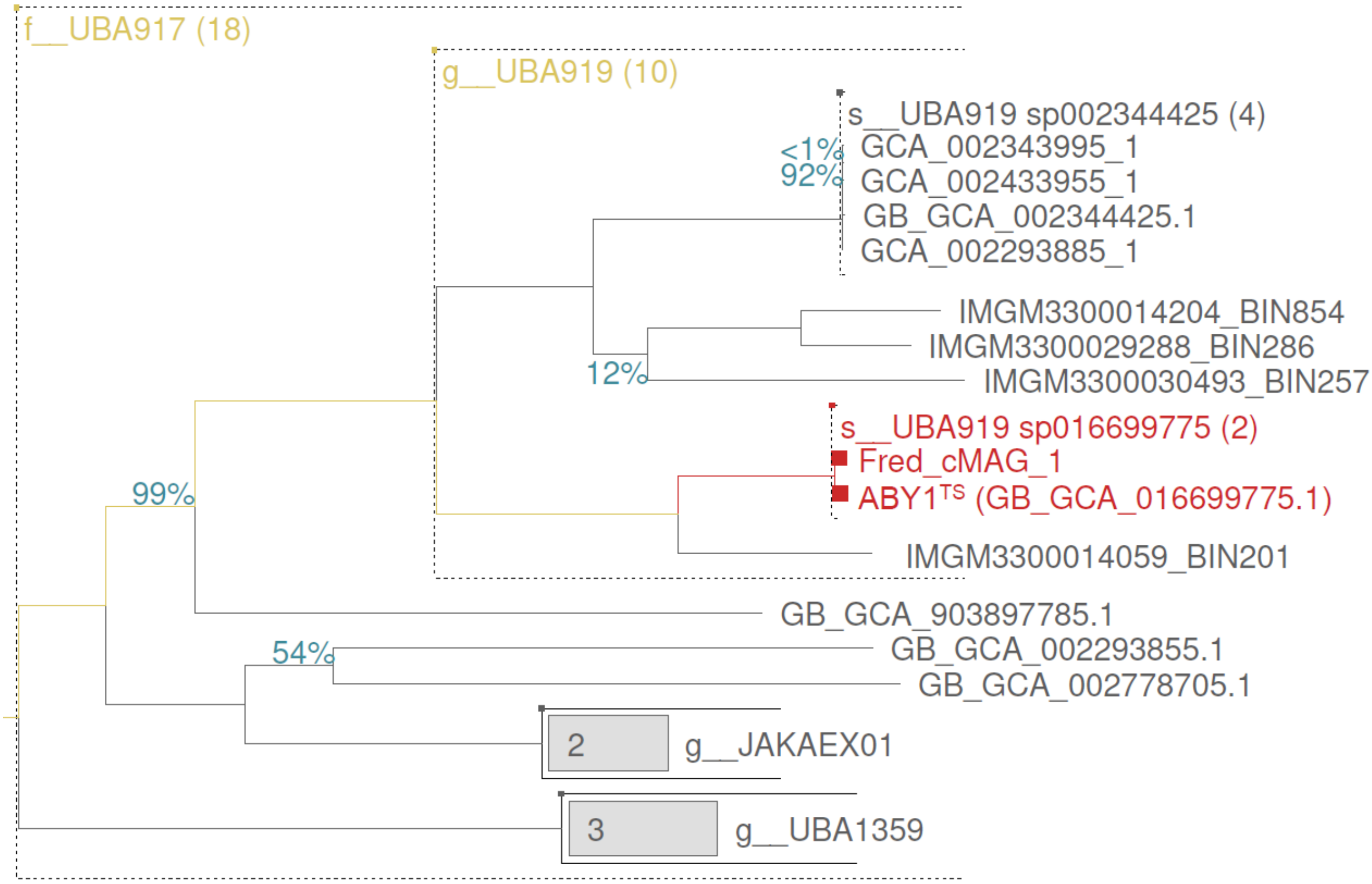


**Fig. S3 | Phylogenomic assignment on the genus and family level of all MAGs included in this study.** Depicted is the genus UBA919 and its sister genera within the family UBA917, representing a section of a phylogenomic tree inferred with FastTree2 from a multiple sequence alignment of 120 marker proteins of 3406 genomes. See **Suppl. Material** for the full tree, confirming the placement of genus UBA919 within *Ca.* Patescibacteria**.** FastTree support values below 100% are shown as numerical values on internal nodes. Nodes without support values have a 100% FastTree support value. Taxonomic assignments are based on GTDB r214 (https://gtdb.ecogenomic.org/). GTDB species UBA919 sp016699775, containing the MAGs ABY1 and Fred_cMAG_1, is highlighted in red, and the former is marked with TS for type species. Dashed lines indicate the genus and family boundaries.

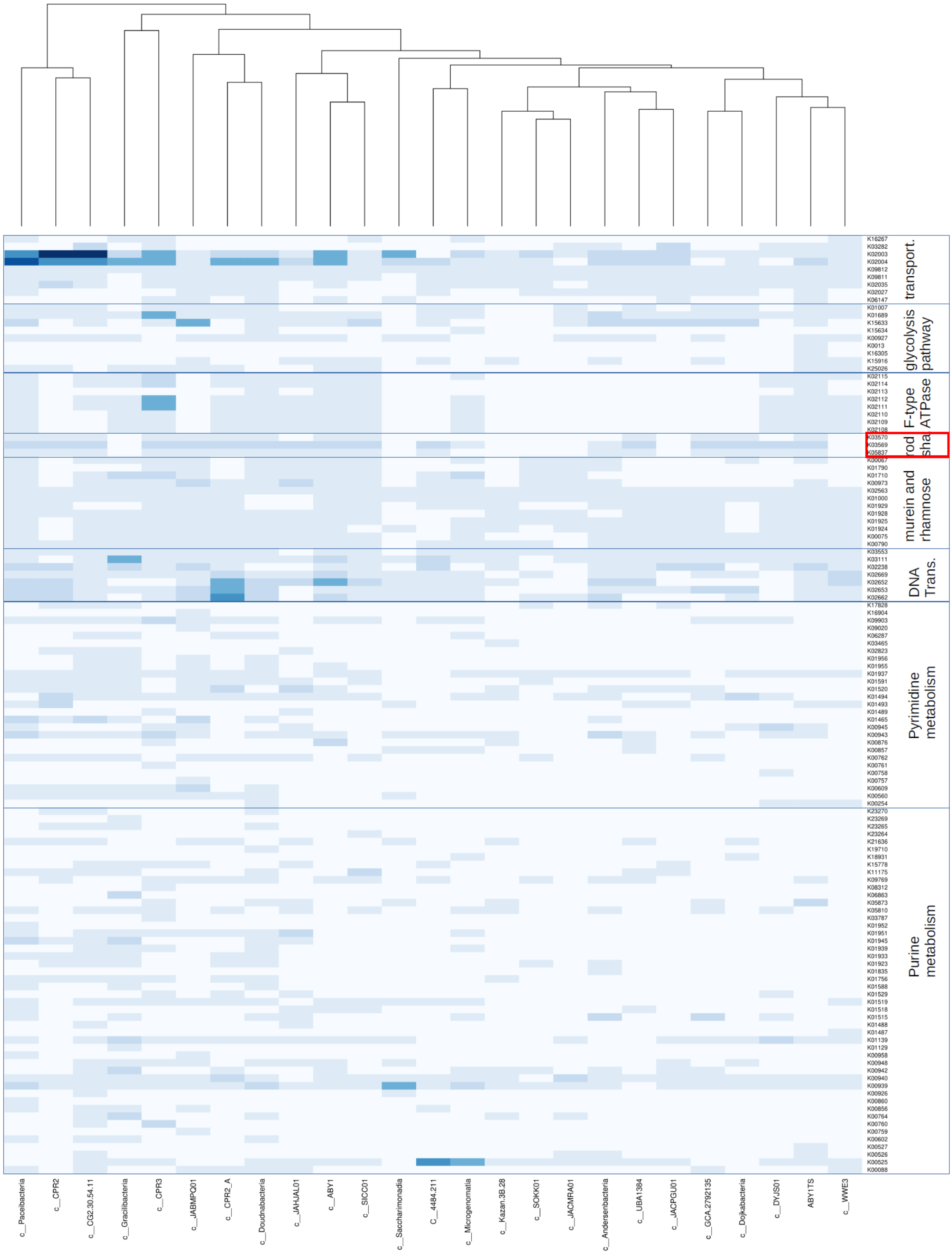


**Fig. S4 | Heat map of inferred functions across *Patescibacteriota* classes.** Shown are the genes counts (white = 0, light blue = 1; blue = 2; dark blue =3) for each genome representative of 24 *Patescibacteriota* classes, that is all except for "c__JAEDAM01" (GCF_021057185.1, *Absconditicoccus praedator*) that lacked BlastKOLA annotations for most of its predicted proteins, present in GTDB r214 and the genome of the proposed type genus and type species P. danicum, labelled as “ABY1TS” (GCA_016699775.1), shown on the second sample column from the right. A detailed list of the class representative genomes is provided in **Table S17**. The functional category “rod shape determining proteins” is highlighted with a **red box**. Note that genes for rod shape determining proteins are present in the majority of class representatives (***17 out of 25***), except for the following eight: c__Gracilibacteria, c__Saccharimonadia, c__4484-211, c__Kazan-3B-28, c__SOKK01, c__JACMRA01, c__JACPGU01, c__WWE3.

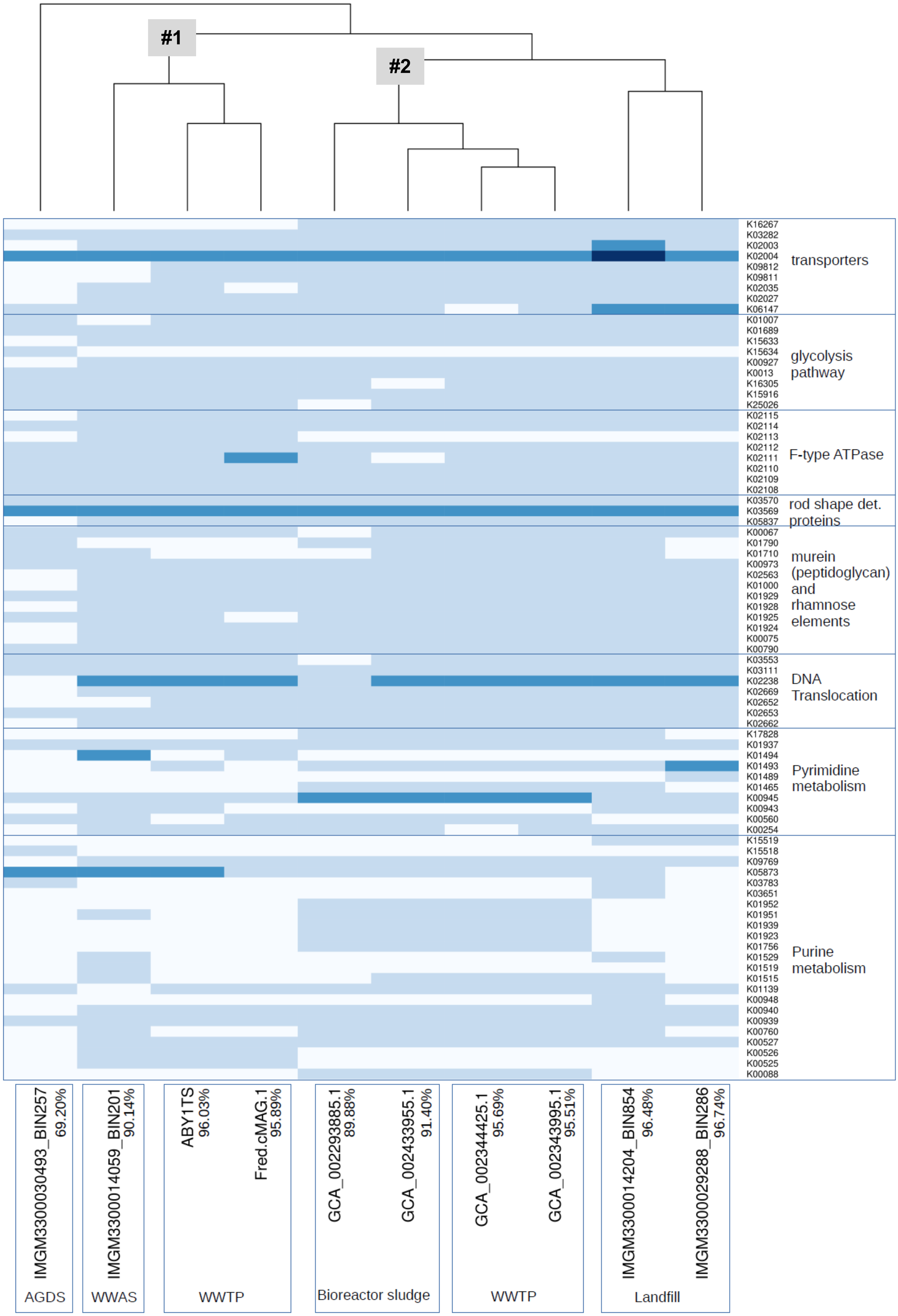
#1
#2
K16267
K03282
K02003
K02004
K09812
K09811
K02035
K02027
K06147
transporters
K01007
K01689
K15633
K15634
K00927
K0013
K16305
K15916
K25026
glycolysis pathway
K02115
K02114
K02113
K02112
K02111
K02110
K02109
K02108
F-type ATPase
K03570
K03569
K05837
rod shape det. proteins
K00067
K01790
K01710
K00973
K02563
K01000
K01929
K01928
K01925
K01924
K00075
K00790
murein (peptidoglycan) and rhamnose elements
K03553
K03111
K02238
K02669
K02652
K02653
K02662
DNA Translocation
K17828
K01937
K01494
K01493
K01489
K01465
K00945
K00943
K00560
K00254
Pyrimidine metabolism
K15519
K15518
K09769
K05873
K03783
K03651
K01952
K01951
K01939
K01923
K01756
K01529
K01519
K01515
K01139
K00948
K00940
K00939
K00760
K00527
K00526
K00525
K00088
Purine metabolism
IMGM3300030493_BIN257 69.20%
AGDS
IMGM3300014059_BIN201 90.14%
WWAS
ABY1TS 96.03%
Fred.cMAG.1 95.89%
WWTP
GCA_002293885.1 89.88%
GCA_002433955.1 91.40%
Bioreactor sludge
GCA_002344425.1 95.69%
GCA_002343995.1 95.51%
WWTP
IMGM3300014204_BIN854 96.48%
IMGM3300029288_BIN286 96.74%
Landfill

**Fig. S5 | Comparison of core encoded proteins (KEGG IDs) of all 10 genomes, assigned to the genus Patescibacterium.** Note that the landfill samples cluster together, and that there are two clusters (#1 and #2) of wastewater related MAGs, whereby #1 contains the type genome of *P. danicum* (ABY1TS). The differences between the wastewater clusters were mainly restricted to pyrimidine and purine metabolism, whereas the landfill MAGs had a higher copy number of two transporters.

Colors indicate the copy number of the inferred proteins: white = 0, light blue = 1; dark blue = 2; darkest blue = 3 (annotated proteins). Abbreviations: WWAS = waste water activated sludge; AGDS = aquatic groundwater deep-subsurface, WWTP = waste water treatment plant.

**(A)**

### Conserved domains on [lcl|Query_2889086]

MOBHLK_04680 hypothetical protein_Fred

View Standard Results

#### Protein Classification

**beta strand repeat-containing protein**( domain architecture ID 11460828)

beta strand repeat-containing protein similar to extracellular S-layer proteins and ice nucleation proteins

#### Graphical summary

Zoom to residue level | hide extra options « | Show site features | Horizontal zoom: × 1 | Update graph

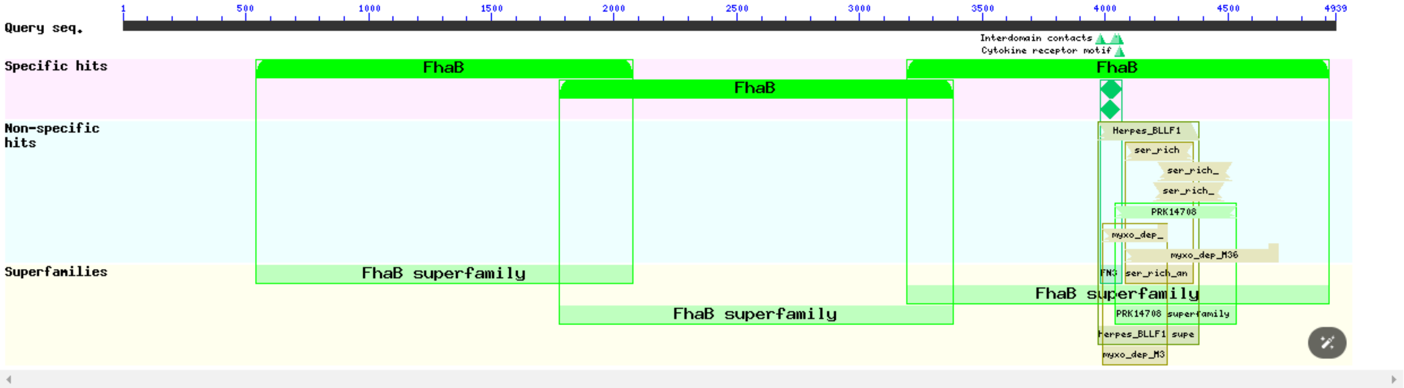


Search for similar domain architectures | Refine search

#### List of domain hits

| Name | Accession | Description | Interval | E-value |
|---|---|---|---|---|
| [+] FhaB | COG3210 | Large exoprotein involved in heme utilization or adhesion [Intracellular trafficking, ... | 3194-4914 | 3.59e-35 |
| [+] FhaB | COG3210 | Large exoprotein involved in heme utilization or adhesion [Intracellular trafficking, ... | 539-2075 | 1.79e-12 |
| [+] FhaB | COG3210 | Large exoprotein involved in heme utilization or adhesion [Intracellular trafficking, ... | 1777-3380 | 5.39e-12 |
| [+] Herpes_BLLF1 | pfam05109 | Herpes virus major outer envelope glycoprotein (BLLF1); This family consists of the BLLF1 ... | 3972-4382 | 2.17e-09 |
| [+] ser_rich_anae_1 | NF033849 | serine-rich protein; This serine-rich protein belongs to a family with large size (over 1000 ... | 4081-4359 | 2.49e-07 |
| [+] FN3 | cd00063 | Fibronectin type 3 domain; One of three types of internal repeats found in the plasma protein ... | 3983-4068 | 3.65e-07 |
| [+] FN3 | smart00060 | Fibronectin type 3 domain; One of three types of internal repeat within the plasma protein, ... | 3983-4058 | 2.91e-06 |
| [+] ser_rich_anae_1 | NF033849 | serine-rich protein; This serine-rich protein belongs to a family with large size (over 1000 ... | 4217-4517 | 4.34e-06 |
| [+] ser_rich_anae_1 | NF033849 | serine-rich protein; This serine-rich protein belongs to a family with large size (over 1000 ... | 4199-4482 | 7.59e-05 |
| [+] PRK14708 | PRK14708 | flagellin; Provisional | 4040-4536 | 6.28e-04 |
| [+] myxo_dep_M36 | NF038112 | myxosortase-dependent M36 family metallopeptidase; Members of this bacterial protein family ... | 3990-4252 | 2.72e-03 |
| [+] myxo_dep_M36 | NF038112 | myxosortase-dependent M36 family metallopeptidase; Members of this bacterial protein family ... | 4083-4703 | 3.14e-03 |

**Blast search parameters**

**Data Source:** Live blast search RID = 8FKWD5JB016

**User Options:** Database: CDSEARCH/cdd  Low complexity filter: no  Composition Based Adjustment: yes  E-value threshold: 0.01  Maximum number of hits: 500

**(B)**

### Conserved domains on [lcl|Query_3699361]

ALDGBO_04135 hypothetical protein_GCA_775

View Standard Results

#### Graphical summary

Zoom to residue level | show extra options »

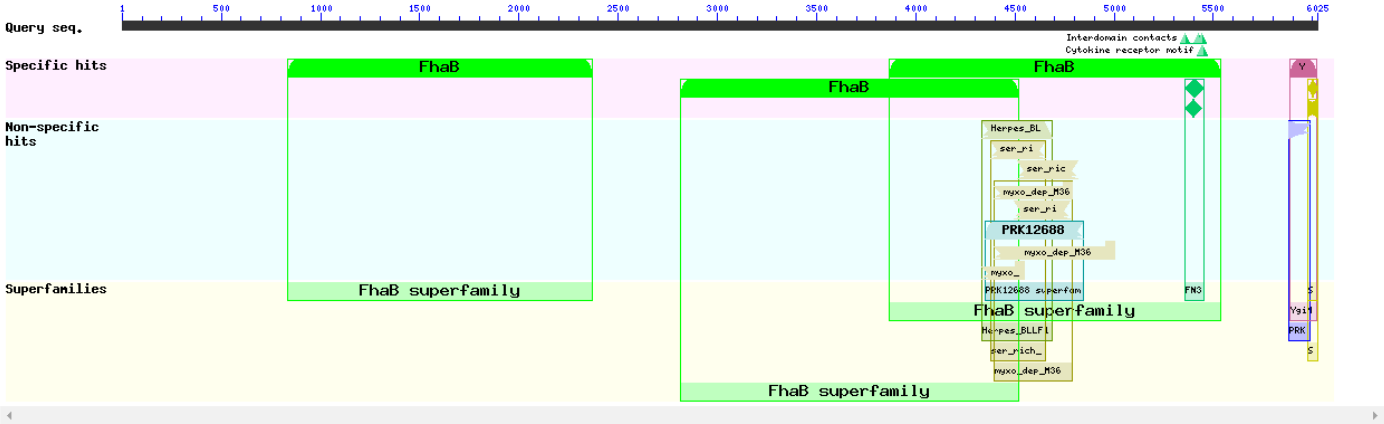


Search for similar domain architectures | Refine search

#### List of domain hits

| Name | Accession | Description | Interval | E-value |
|---|---|---|---|---|
| [+] FhaB | COG3210 | Large exoprotein involved in heme utilization or adhesion [Intracellular trafficking, ... | 3867-5534 | 5.42e-40 |
| [+] FhaB | COG3210 | Large exoprotein involved in heme utilization or adhesion [Intracellular trafficking, ... | 2813-4518 | 3.10e-20 |
| [+] YgiM | COG3103 | Uncharacterized conserved protein YgiM, contains N-terminal SH3 domain, DUF1202 family ... | 5886-6019 | 5.70e-15 |
| [+] FN3 | cd00063 | Fibronectin type 3 domain; One of three types of internal repeats found in the plasma protein ... | 5359-5450 | 5.92e-15 |
| [+] FhaB | COG3210 | Large exoprotein involved in heme utilization or adhesion [Intracellular trafficking, ... | 836-2372 | 6.19e-14 |
| [+] Herpes_BLLF1 | pfam05109 | Herpes virus major outer envelope glycoprotein (BLLF1); This family consists of the BLLF1 ... | 4331-4688 | 2.96e-09 |
| [+] FN3 | smart00060 | Fibronectin type 3 domain; One of three types of internal repeat within the plasma protein, ... | 5359-5441 | 1.87e-08 |
| [+] ser_rich_anae_1 | NF033849 | serine-rich protein; This serine-rich protein belongs to a family with large size (over 1000 ... | 4378-4656 | 7.72e-08 |
| [+] ser_rich_anae_1 | NF033849 | serine-rich protein; This serine-rich protein belongs to a family with large size (over 1000 ... | 4514-4814 | 1.86e-06 |
| [+] SH3_3 | pfam08239 | Bacterial SH3 domain; | 5975-6025 | 2.91e-06 |
| [+] myxo_dep_M36 | NF038112 | myxosortase-dependent M36 family metallopeptidase; Members of this bacterial protein family ... | 4397-4786 | 8.41e-06 |
| [+] ser_rich_anae_1 | NF033849 | serine-rich protein; This serine-rich protein belongs to a family with large size (over 1000 ... | 4496-4779 | 3.07e-05 |
| [+] PRK12688 | PRK12688 | flagellin; Reviewed | 4348-4843 | 4.13e-05 |
| [+] SH3b | smart00287 | Bacterial SH3 domain homologues; | 5973-6020 | 4.76e-04 |
| [+] myxo_dep_M36 | NF038112 | myxosortase-dependent M36 family metallopeptidase; Members of this bacterial protein family ... | 4394-5000 | 1.33e-03 |
| [+] PRK13914 | PRK13914 | invasion associated endopeptidase; | 5882-5986 | 6.21e-03 |
| [+] myxo_dep_M36 | NF038112 | myxosortase-dependent M36 family metallopeptidase; Members of this bacterial protein family ... | 4332-4549 | 7.49e-03 |

**Blast search parameters**

**Data Source:** Live blast search RID = 8FN6YDNM016

**User Options:** Database: CDSEARCH/cdd  Low complexity filter: no  Composition Based Adjustment: yes  E-value threshold: 0.01  Maximum number of hits: 500

**(C)** Protein size distribution across all MAGs

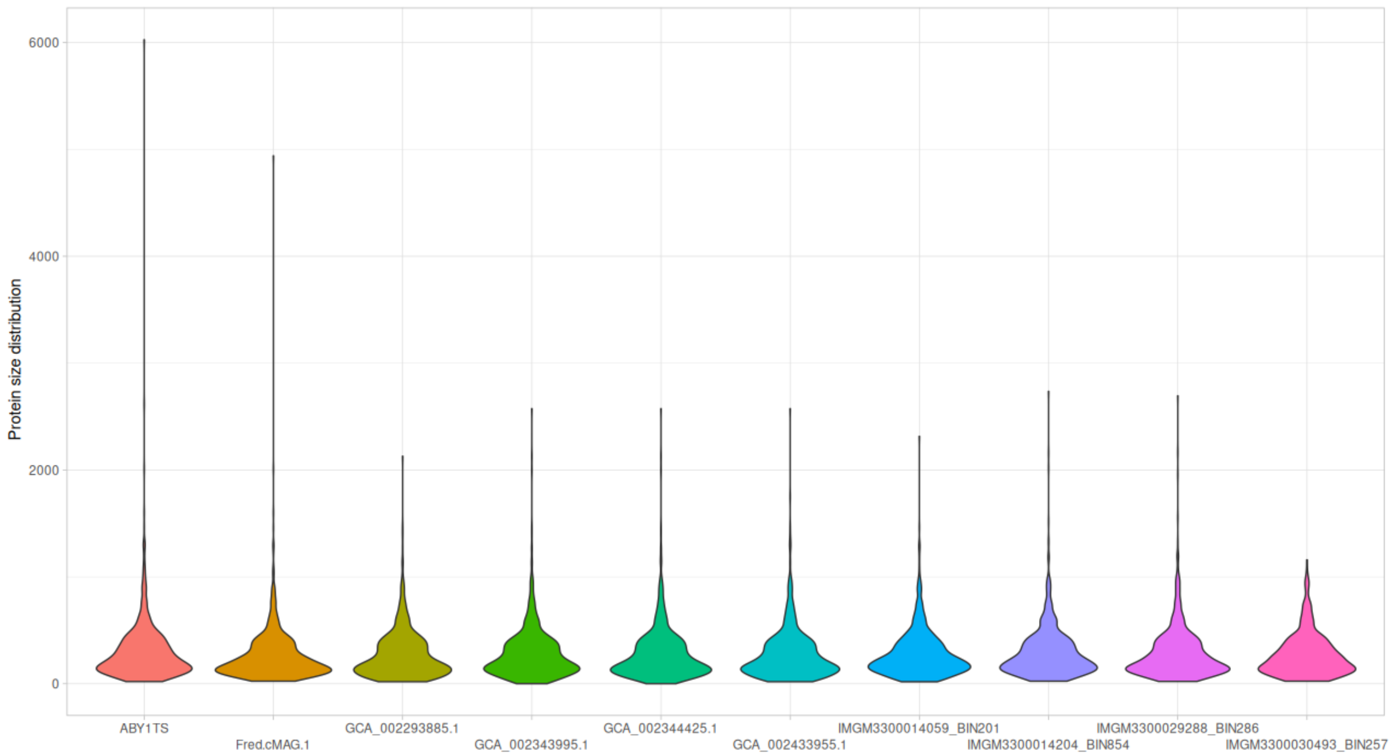


**Fig. S6 | Annotations of large proteins in *Patescibacterium* gen. nov.** Blastp search results of the largest protein encoded in the genome of 'Fred.cMAG.1' consisting of 4939 amino acids (**A**) and ABY1$^{TS}$ consisting of 6025 amino acids (**B**). Both of these large proteins have multiple hits, with considerably low e-value, to FhaB (COG3210), a large exoprotein involved in heme utilization or adhesion. **(C)** Protein size (amino acid counts) distribution across all MAGs.

### ABY1TS:

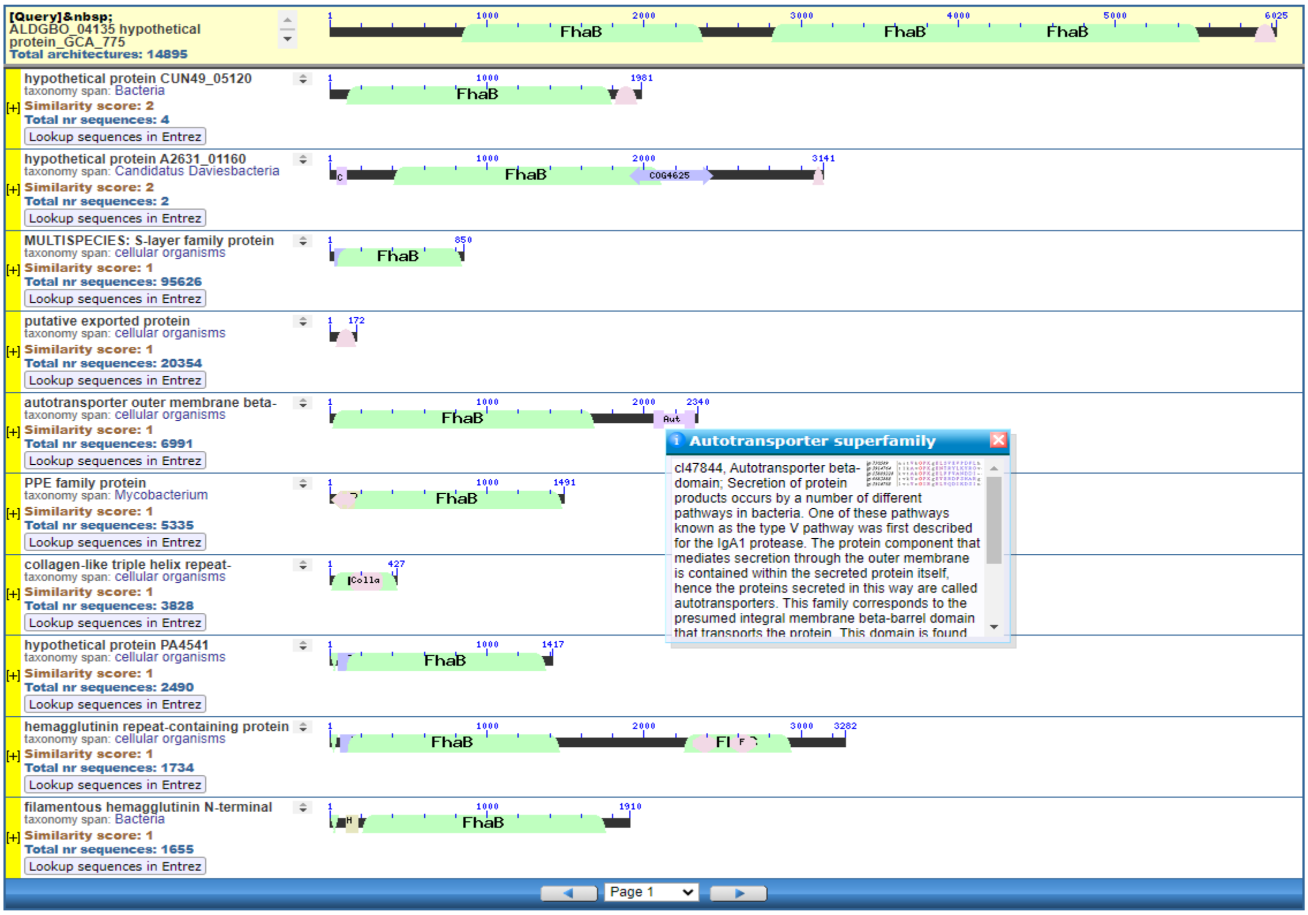


### Fred.cMAG.1:

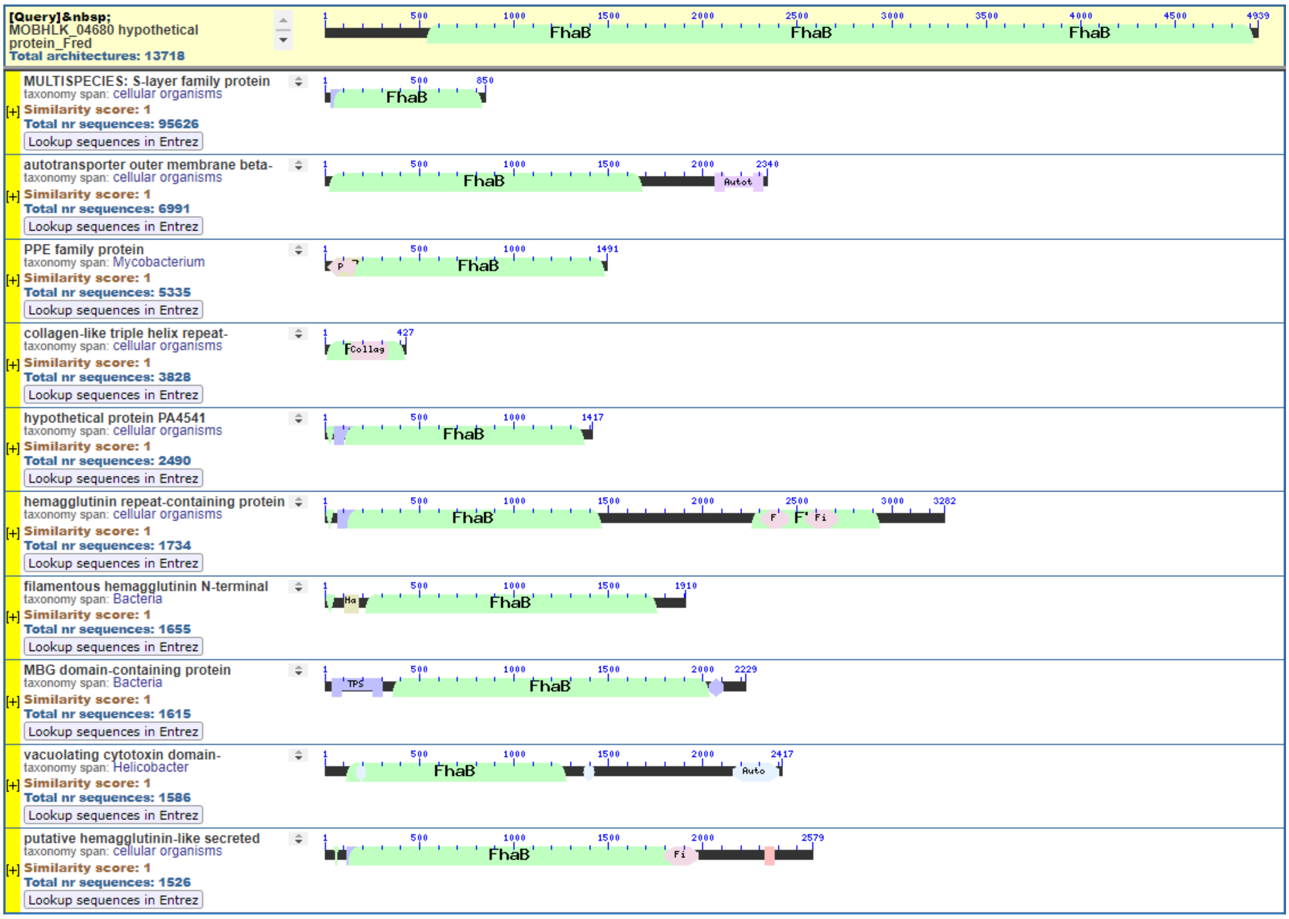

**Fig. S7 | Searching protein homology by domain architecture.** The Conserved Domain Architecture Retrieval Tool (CDART) (https://www.ncbi.nlm.nih.gov/Structure/lexington/docs/cdart_about.html) was used to find protein similarities (domain architectures) across significant evolutionary distances. A domain architecture is defined as the sequential order of conserved domains (functional units) in a protein sequence. Note that hits for Fred.cMAG.1 and ABY1$^{TS}$ contain proteins with downstream autotransporter beta-domains, e.g. hits number 2 and 9 of the query Fred ('Fred.cMAG.1'). These protein components are known to mediate secretion through the outer membrane and are thereby contained within the secreted protein itself, therefore the proteins secreted in this way are called autotransporters.

**GIW59270** 9304 aa

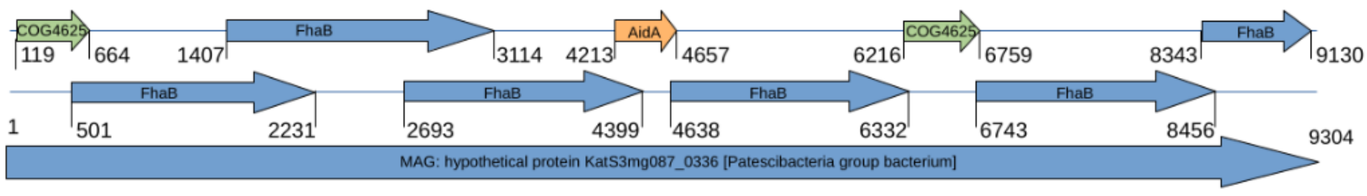


**NCBI protein:** hypothetical protein KatS3mg087_0336 [Patescibacteria group bacterium]
**GTDB taxonomy:** p__Patescibacteria;c__Microgenomatia;o__Daviesbacterales;f__;g__;s__*

**OGG71089** 6999 aa

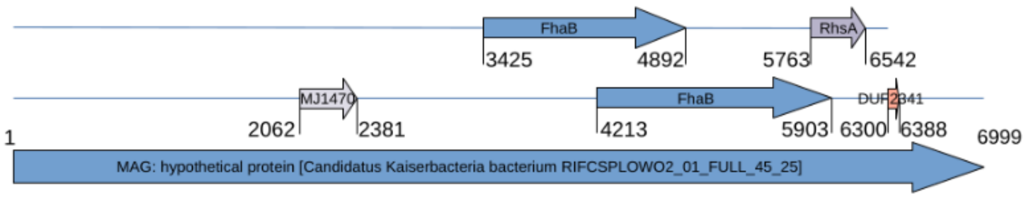


**NCBI protein:** hypothetical protein A3G90_03050 [Candidatus Kaiserbacteria bacterium RIFCSPLOWO2_12_FULL_45_26]
**GTDB taxonomy** (GCA_001781805.1):
p__Patescibacteria;c__Paceibacteria; o__UBA9973; f__UBA918; g__OLB19; s__OLB19 sp001781415

**QQS60272** 6025 aa

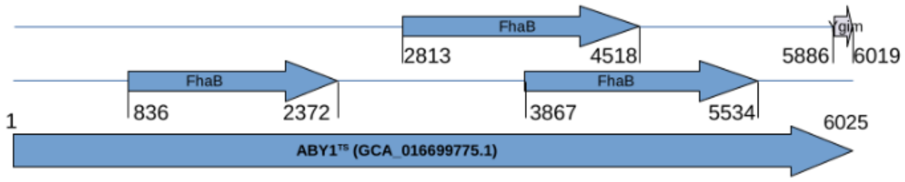


ABY1$^{TS}$ (GCA_016699775.1)

**QQR65269** 5626 aa

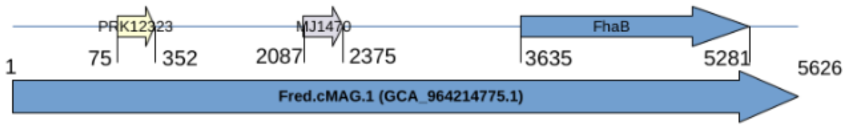


Fred.cMAG.1 (GCA_964214775.1)

**taxonomy* was obtained from the Suppl. Table S3 published in (Kato et al., 2022) #4439

**Fig. S8 | Domains of large *Patescibacteriota* proteins.** Shown are the unusually long proteins detected in the genomes of ABY1$^{TS}$ (GCA_016699775.1) and Fred.cMAG.1 (GCA_964214775.1) compared to two representative proteins, from the 149 long proteins obtained from NCBI. Note that the **DUF2341** domain, predicted component of type IV pili-like system, is shown as the COG annotation "Uncharacterized conserved protein **MJ1470** (COG5306)". ***Annotations***: **COG4625** =Uncharacterized conserved protein, contains a C-terminal beta-barrel porin domain [Function unknown];

**FhaB** = Large exoprotein involved in heme utilization or adhesion [Intracellular trafficking, secretion, and vesicular transport]; **AidA** = Autotransporter adhesin AidA [Cell wall/membrane/envelope biogenesis, Intracellular trafficking, secretion, and vesicular transport]; **COG4625** = Uncharacterized conserved protein, contains a C-terminal beta-barrel porin domain [Function unknown]; **MJ1470** = Uncharacterized conserved protein MJ1470, contains DUF2341 domain, predicted component of type IV pili-like system [General function prediction only]; **RhsA** = Uncharacterized conserved protein RhaS, contains 28 RHS repeats [General function prediction only]; **DUF2341** = Domain of unknown function (DUF2341); **YgiM** = Uncharacterized conserved protein YgiM, contains N-terminal SH3 domain, DUF1202 family [General function prediction only]; **PRK12323** = DNA polymerase III subunit gamma/tau.

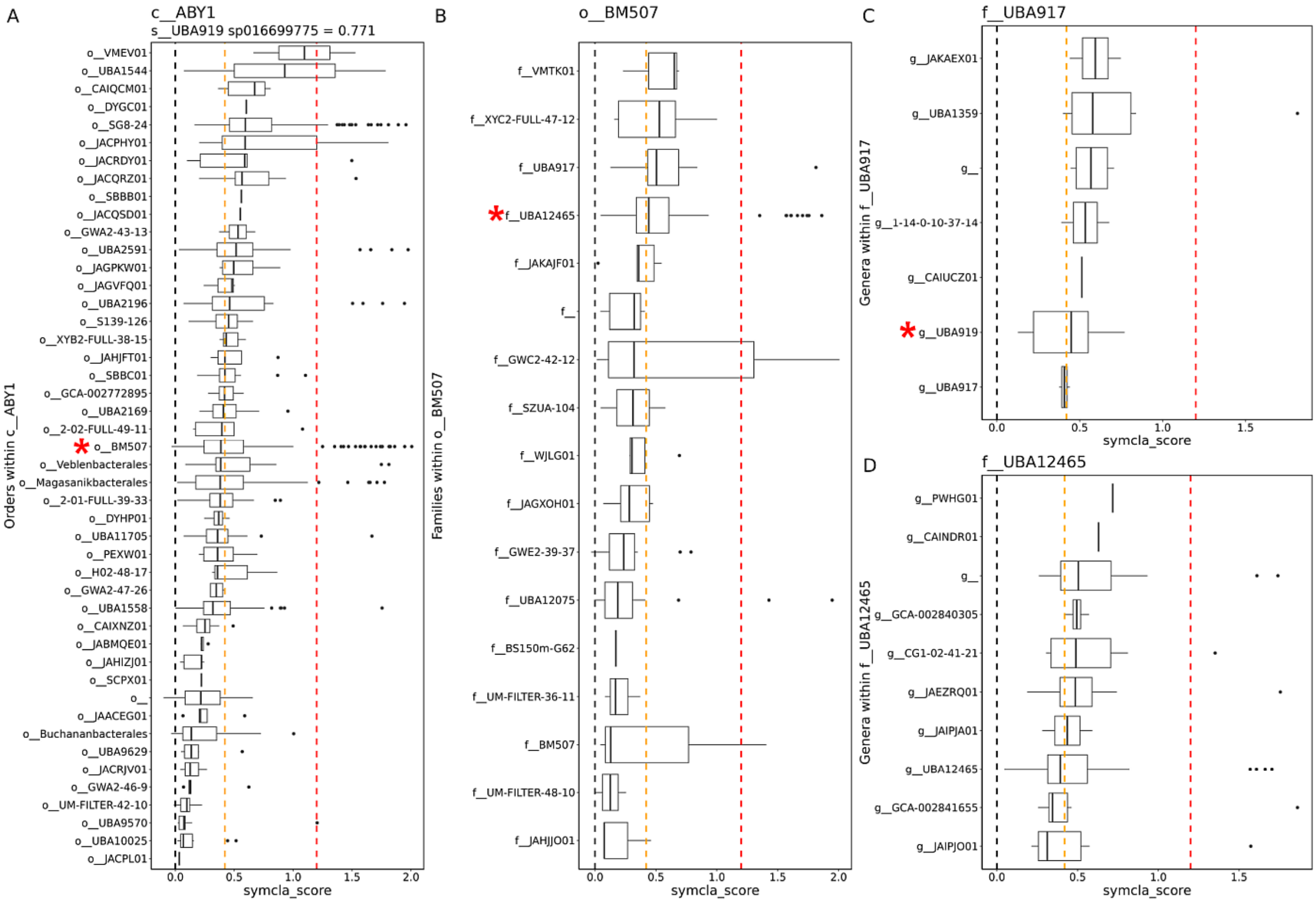


**Fig. S9 | Predicting the lifestyle of *Patescibacteria* gen. nov. (UBA919).** Shown are the scores of the symcla analysis of Patescibacteriota phyl. nov. lineages. **(A)** Symcla scores for all orders of the class ABY1. Order BM507 is marked with a red asterisk. **(B)** Symcla scores for all families within the order BM507. Family UBA917 is marked with a red asterisk. **(C)** Symcla scores of all genera in the family UBA917, compared to the symcla scores in the sister family UBA12465 **(D)**. Note that the following categories have been defined based on symcla scores: < 0.42 = free-living (orange dashed line), >= 0.42 - <= 1.20= symbiont host-associated, >= 1.20 symbiont intracellular (red dashed line). According to the developers of symcla, this tool was designed to minimize the rate of false positives for symbionts, at the expense of increased false negatives (i.e. some symbionts might still get a lower symcla score).

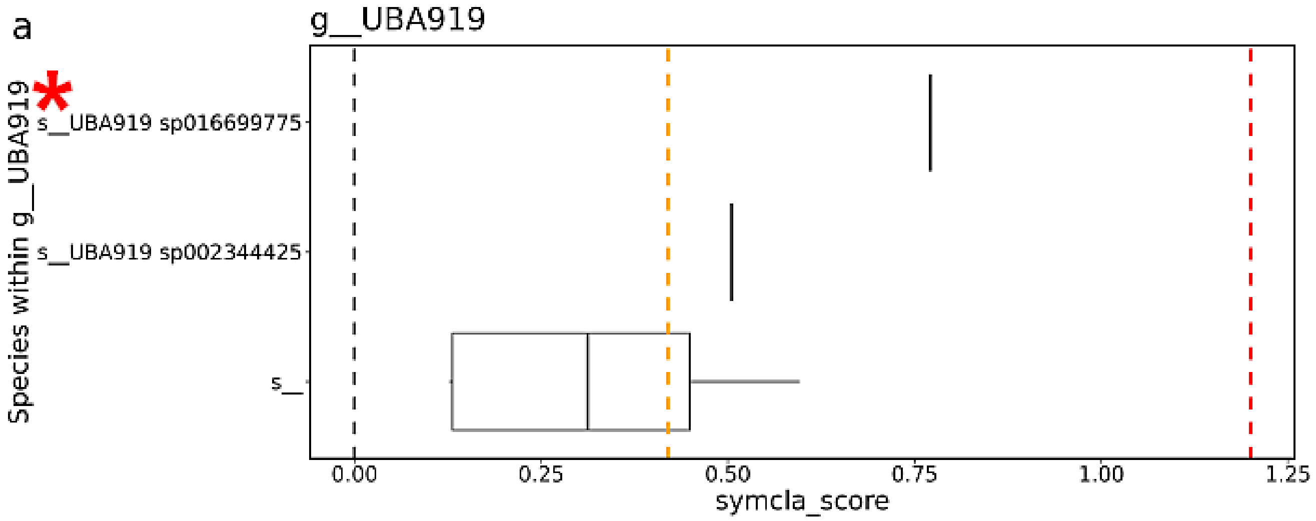


**Fig. S10 | Predicting the lifestyle of *Patescibacteria* gen. nov. (UBA919).** Shown are the symcla analysis scores of species within the genus Patescibacteria gen. nov., (former UBA919). The genome ABY1$^{TS}$ (GCA_016699775.1), that has been proposed as the type genome of the species *Patescibacterium danicum* sp. nov., is shown as s__UBA919 sp016699775 (highlighted with a red asterisk) in this plot and has a symcla score of 0.771, that is by far the highest in this genus, and is right in the middle of the host associated symbiont category (0.42 >= x < 1.2). For the purpose of this analysis, all 4 MAGs recovered from IMG were treated as one lineage, shown here as "s__". Note that the following categories have been defined based on symcla scores: < 0.42 = free-living (orange dashed line), >= 0.42 - <= 1.20= symbiont host-associated, >= 1.20 symbiont intracellular (red dashed line).

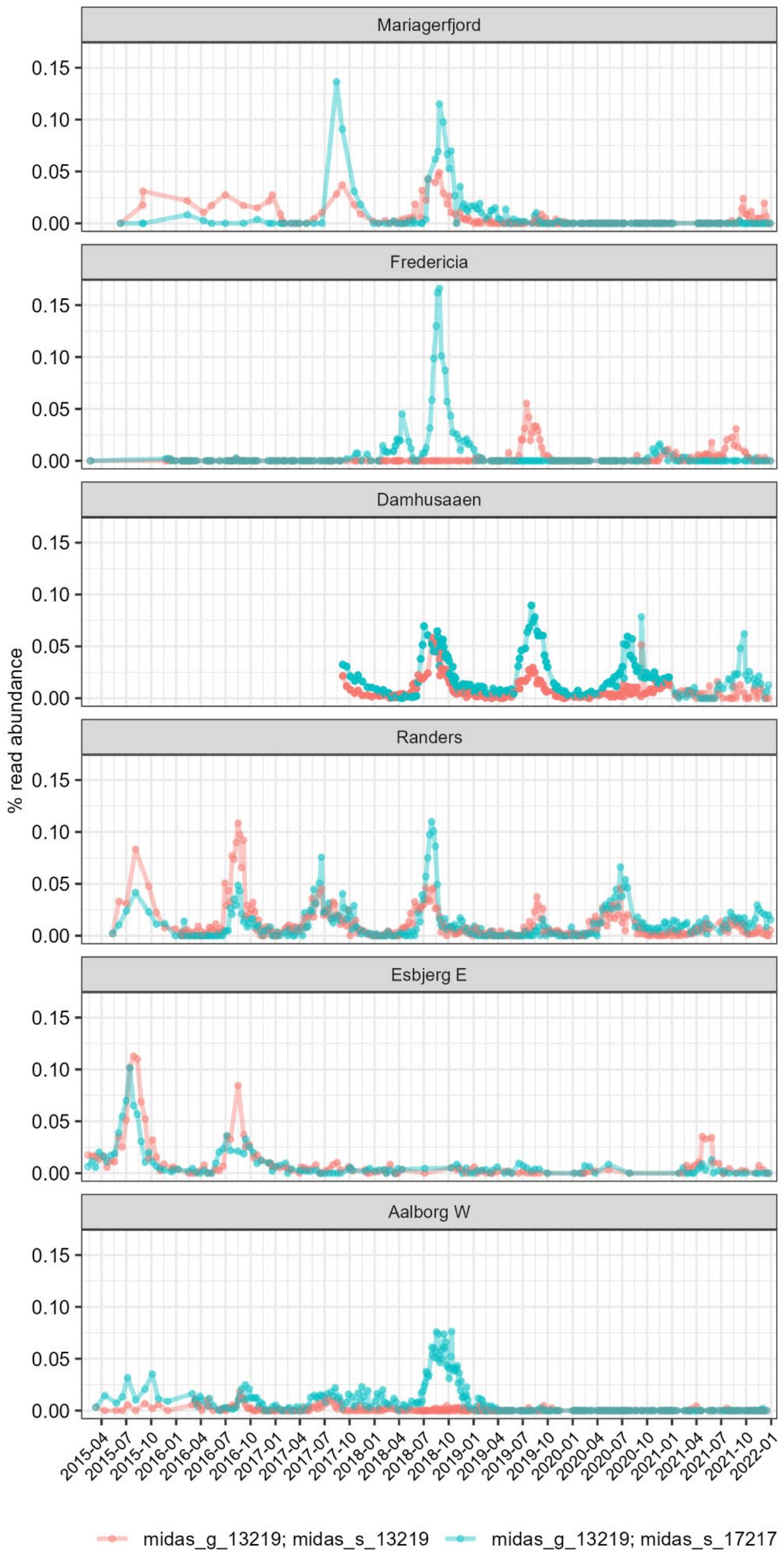


**Fig. S11 | Relative 16S rRNA gene abundances of *Patescibacterium danicum* during a time series obtained from Danish WWTPs**. The species “midas_s_17217” (shown in cyano) includes the 16S rRNA gene sequence of the MAGs ABY1TS and Fred.cMAG.1 assigned to *Patescibacterium danicum*. The taxon midas_s_13219 (shown in salmon) is another species in the same genus, only known on the 16S level. Note, that we see peaks in abundance for summer/autumn. The time series included data over 4 - 6 years, sampled weekly, from 6 Danish WWTPs with nutrient removal.

| | C Australia AU-30 | C India IN-15 | C Uruguay UY-14 | C Uruguay UY-15 | C,N,DN Belgium BE-13 | C,N,DN China CN1-01 | C,N,DN Malaysia MY-16 | C,N,DN Malaysia MY-17 | C,N,DN Poland PL-18 | C,N,DN United States US1-106 | C,N,DN United States US1-107 | C,N,DN United States US1-123 | C,N,DN,P Malaysia MY-07 | C,N,DN,P Singapore SG-01 |
|---|---|---|---|---|---|---|---|---|---|---|---|---|---|---|
| midas_g_13219; midas_s_32630 | 0 | 0.257 | 0 | 0 | 0 | 0 | 0.065 | 0.125 | 0 | 0 | 0 | 0 | 0 | 0.073 |
| midas_g_13219; midas_s_13219 | 0.119 | 0 | 0.061 | 0.050 | 0.041 | 0 | 0 | 0 | 0 | 0 | 0.229 | 0 | 0 | 0 |
| midas_g_13219; midas_s_47379 | 0 | 0 | 0 | 0 | 0 | 0 | 0 | 0 | 0 | 0.047 | 0 | 0.337 | 0 | 0 |
| **midas_g_13219; midas_s_17217** | 0 | 0.068 | 0 | 0 | 0.001 | 0.070 | 0.004 | 0.001 | 0.074 | 0 | 0 | 0.002 | 0 | 0.179 |
| midas_g_13219; midas_s_55049 | 0 | 0 | 0 | 0 | 0 | 0 | 0 | 0 | 0 | 0 | 0.080 | 0 | 0.077 | 0.057 |

**Fig. S12 | Biogeography of *Patescibacteriota* based on 16S rRNA gene surveys.** Shown are relative abundances calculated from Illumina 16S rRNA gene profiles. Included are taxa from genus “midas_g_13219” in the MIDAS database, i.e. the species “midas_s_17217” (highlighted in bold), which includes the 16S rRNA gene sequence of the MAGs ABY1$^{TS}$ and Fred.cMAG.1 assigned to *Patescibacterium danicum*. Samples were obtained from WWTPs across a range of countries and continents (see Methods). Samples include WWTPs designed for carbon removal with nitrification and denitrification (C, N, DN), for carbon removal only (C), and for carbon removal with nitrogen and enhanced biological phosphorus removal (C, N, DN, P).

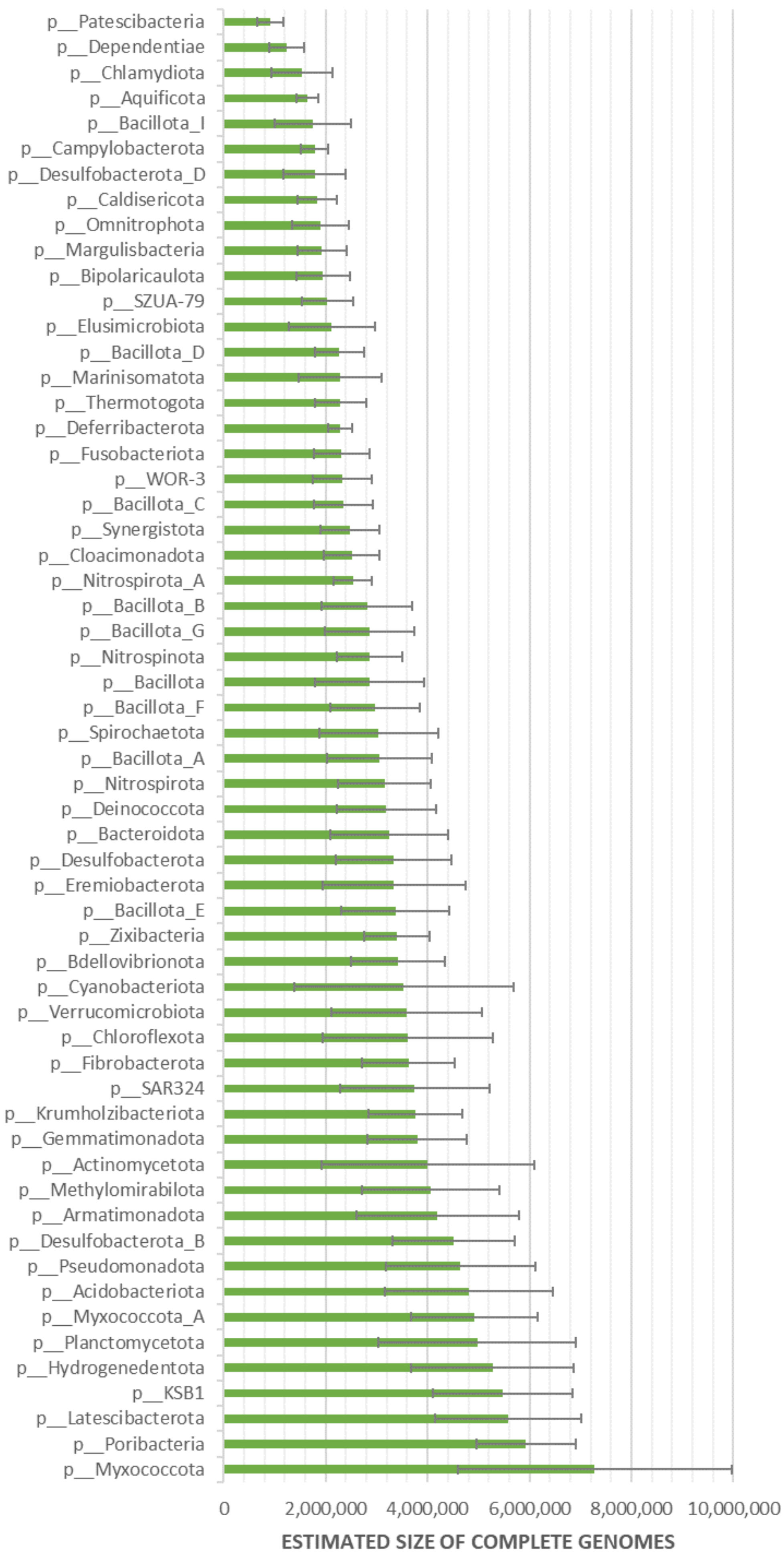


**Fig. S13 | Estimated complete genome sizes for phyla in GTDB.** The 'complete' genome sizes were calculated based on all genomes assigned to a phylum. Note that only phyla present in the GTDB release 09-RS220, that contained at least 100 genomes, were included in the analysis. The resulting average complete genome sizes per phylum ranged from 919.3 kbp ∓ 256 kbp in *Patescibacteriota* (here shown as Patescibacteria, the current name used in GTDB) to 7.3 Mbp ∓ 2.7 Mbp in *Myxococcota*.

**(a)** ABY1$^{TS}$ LSU_rRNA_bacteria with insert **(b)** Fred.cMAG.1 LSU_rRNA_bacteria with insert

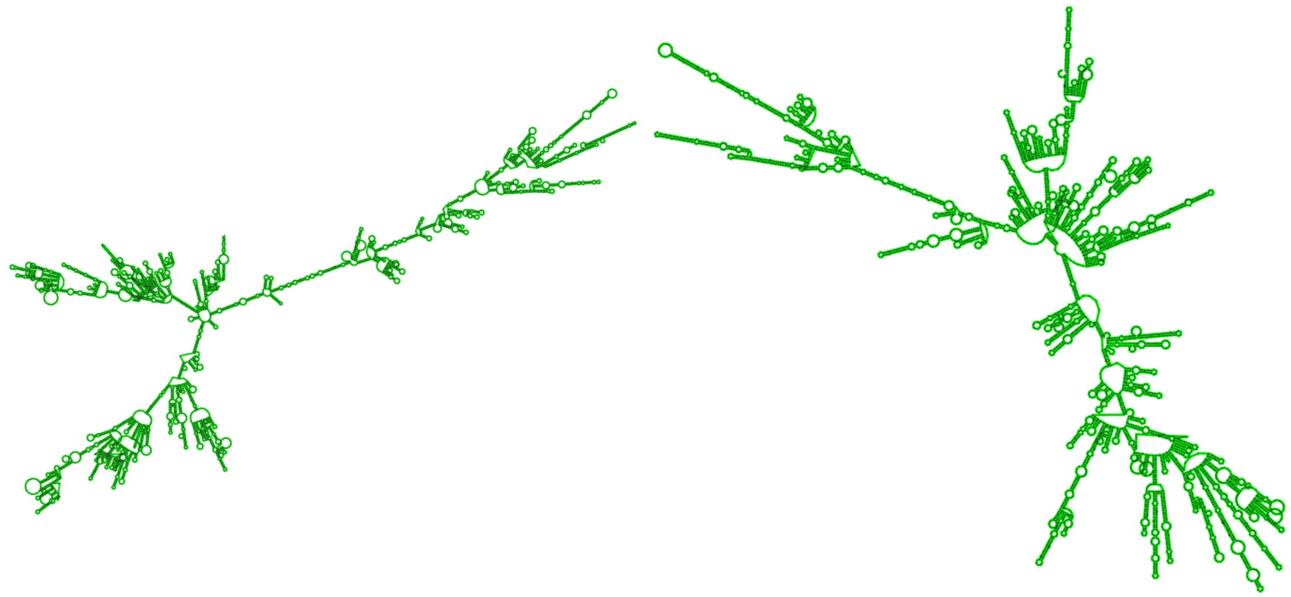

**(c)** ABY1$^{TS}$ LSU_rRNA_bacteria without insert **(d)** Fred.cMAG.1 LSU_rRNA_bac without insert

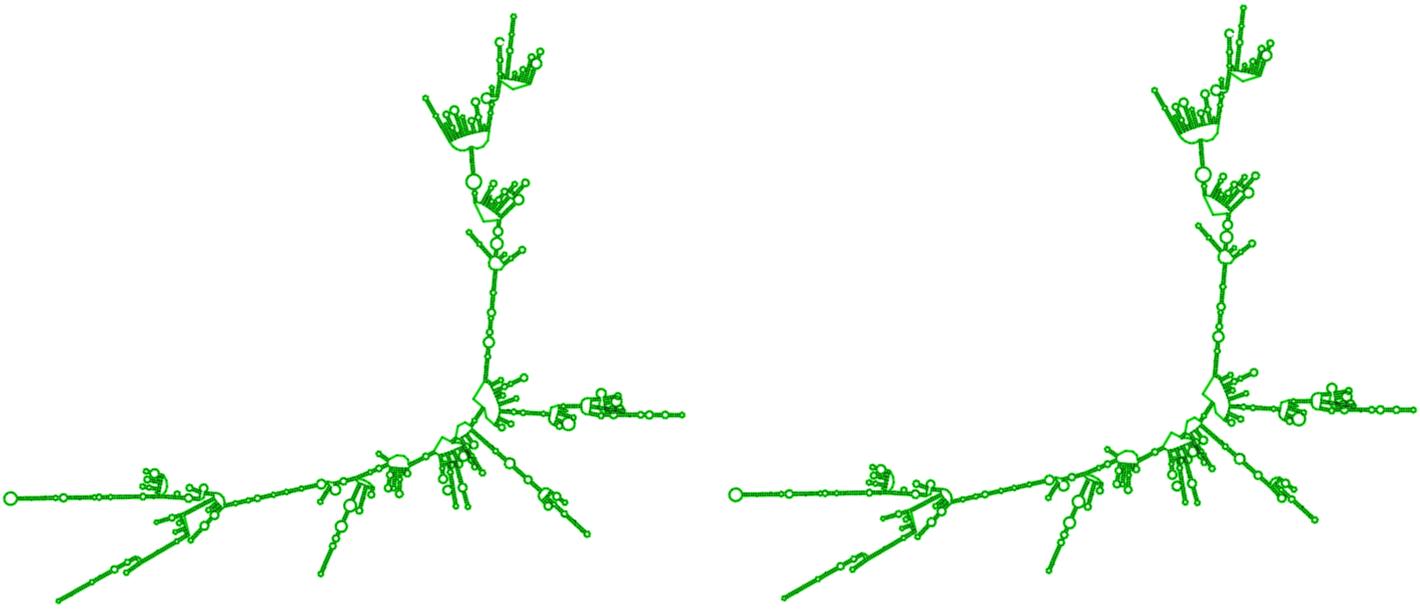

**Fig. S14 | Predicted secondary structure of ribosomal 23S RNA before (a, b) and after intron splicing (c, d).** All predictions were made using the web server StructRNAfinder (see Methods). The post-splicing secondary structure of ABY1$^{TS}$ and Fred LSU (c and d) resembles a functional 23S rRNA.

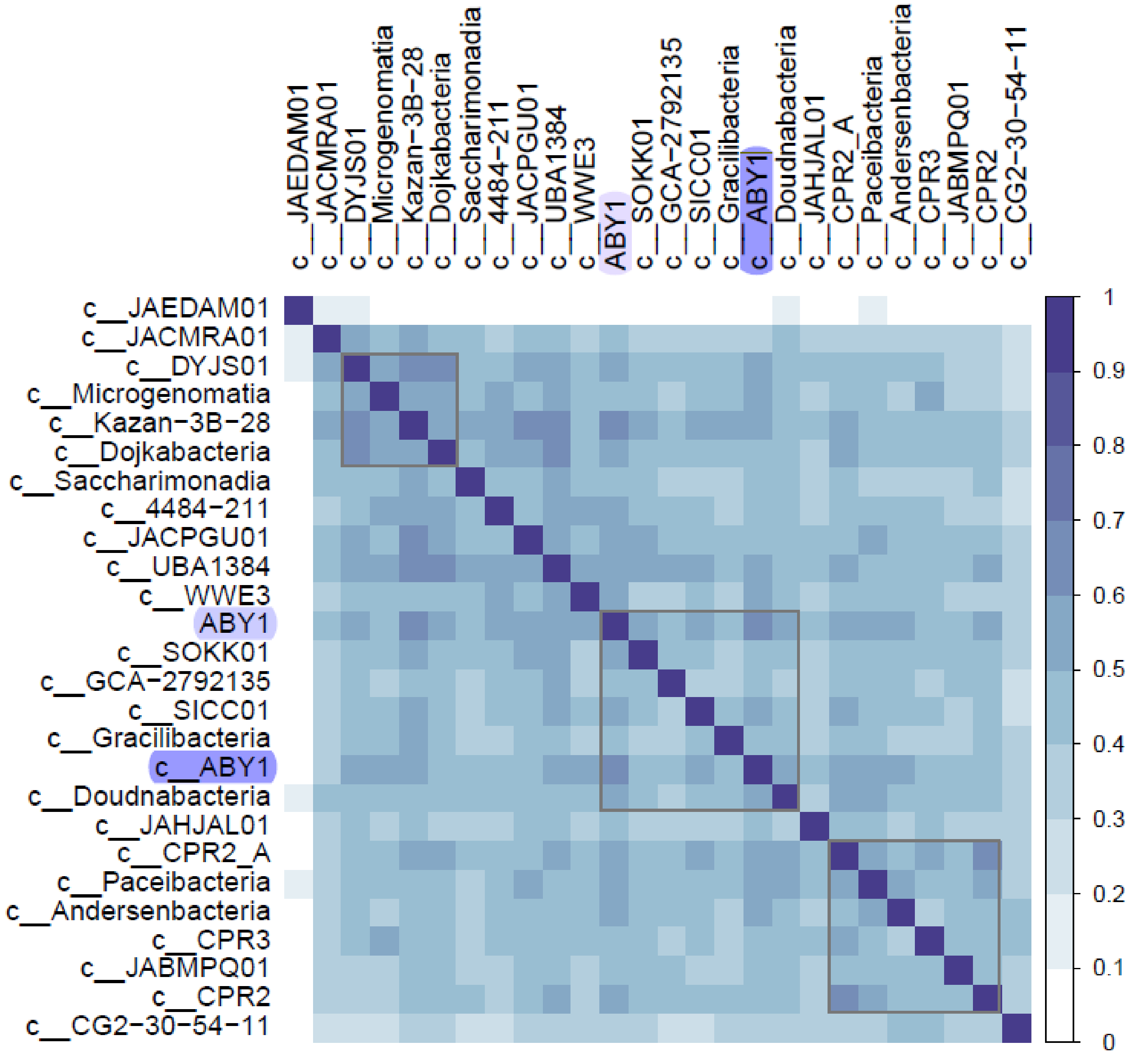


**Fig. S15 | Inferred functions of *Patescibacterium*, the type genus of the phylum, and *Patescibacteriota* classes.** The presence/absence patterns are based on encoded proteins (KEGG IDs) for each genome representative of the 25 *Patescibacteriota* classes present in GTDB r214 and the genome of the proposed type genus and type species *P. danicum*, ABY1$^{TS}$ (GCA_016699775.1). Colors of the scale on the right correspond to the degree of correlation, i.e. 0 stands for no correlation, and 1 for maximal correlation. Note, the lowest correlation was observed for the representative of the class "c__JAEDAM01" (GCF_021057185.1, *Absconditicoccus praedator*) that lacked BlastKOLA annotations for most of its predicted proteins. **Abbreviations**: ABY1 = the genome ABY1$^{TS}$ (GCA_016699775.1); **c__ABY1** = representative of the class ABY1 (Patescibacteria). A detailed list of all class representative genomes is provided in **Table S17.**
